# Cortical gradient perturbation in attention deficit hyperactivity disorder correlates with neurotransmitter-, cell type-specific and chromosome- transcriptomic signatures

**DOI:** 10.1101/2023.04.05.535657

**Authors:** Zhiyi Chen, Ting Xu, Xuerong Liu, Benjamin Becker, Wei Li, Kuan Miao, Zheng Gong, Rong Zhang, ZhenZhen Huo, Bowen Hu, Yancheng Tang, Zhibin Xiao, Zhengzhi Feng, Ji Chen, Tingyong Feng

**Author notes:** Corresponding at: Zhiyi Chen or Zhengzhi Feng, Experimental Research Center for Medical and Psychological Science (ERC-MPS), School of Psychology, Third Military Medical University, Chongqing, 400038 P.R. China; or Tingyong Feng, Faculty of Psychology, Southwest University, Chongqing, 400415, P.R. China.

## Abstract

Neurofunctional dysregulations in spatially discrete areas or isolated pathways have been suggested as neural markers for attention deficit hyperactivity disorder (ADHD). However, multiscale perspectives into the neurobiological underpins of ADHD spanning multiple biological systems remain sparse. This points to the need of multi-levels of analysis encompassing brain functional organization and its correlation with molecular and cell-specific transcriptional signatures are stressed. Here, we capitalized on diffusion mapping embedding model to derive the functional connectome gradient, and deployed multivariate partial least square (PLS) method to uncover the enrichment of neurotransmitomic, cellular and chromosomal connectome-transcriptional signatures of ADHD. Compared to typical control, ADHD children presented connectopic cortical perturbations in lateral orbito-frontal and superior temporal regions, which had also been validated in another independent sample. This gradient-derived variants in ADHD further aligned spatially with distributions of GABA_A/BZ_ and 5-HT_2A_ receptors and co-varied with genetic transcriptional expression. Cognitive decoding and gene-expression annotation showed the correlates of these variants in memory, emotional regulation and spatial attention. Moreover, the gradient-derived transcriptional signatures of ADHD exhibited enriched expression of oligodendrocyte precursors and endothelial cells, and were mainly involved as variants of chromosome 18, 19 and X. In conclusion, our findings bridged in-vivo neuroimging assessed functional brain organization patterns to a multi-level molecular pathway in ADHD, possibly shedding light on the interrelation of biological systems that may coalesce to the emergence of this disorder.

## INTRODUCTION

Attention deficit hyperactivity disorder (ADHD) is a neurodevelopmental disorder characterized by persistent symptoms of inattention, hyperactivity and impulsivity with an estimated 5 % of children worldwide being affected^1, 2^. ADHD has been associated with cognitive and social impairments across the lifespan, and in turn an increase in the risk for psychiatric disorders^3^. Further, the economic strain that ADHD puts on the public healthcare systems has strongly increased, with $ 21,000 per year on average^4^. Despite the tremendous negative consequences of ADHD as a multi-scale biologically hierarchical deficit, the precise neuro-pathophysiological mechanisms of ADHD have been rarely investigated, particularly the neuroimage-derived neurotransmitomic, molecular and transcriptomic vulnerability.

Given that ADHD has been conceptualized as brain disconnection syndrome^5^, examining how the brain functional connectome pattern develops and whether this could be a robust neural phenotype for neurodevelopmental dysregulations in ADHD patients is a promising venue. Functional connectome approaches that capture temporal correlation of coherent spontaneous fluctuations among brain regions have been widely applied for identifying brain biomarkers for clinical conditions^6^. Regarding childhood ADHD, different lines of large-scale brain network studies have reported dysfunctional connectivity patterns in the default mode and attention relevant brain networks, which were linked with ADHD symptoms, such as impaired executive control and distractibility^7^. However, it remains unclear whether ADHD-related functional network connectivity alterations reflect dysregulations in a single global pattern or multiple patterns across levels of cortical hierarchical architecture organization. Within this context, the recently described spatial gradients of functional integration from primary sensory network to transmodal hierarchy may allow to determine the precise alterations^8–10^.

Hierarchical organization changes in the primary-to-transmodal cortex during childhood and its developmental trajectories may support a facilitation of cognition, such as working memory and attention^11, 12^. A recent study suggested dysfunctional pattern of the primary-to-transmodal connectome gradient in childhood-onset autism spectrum disorder, which may underlie neurodevelopmental deficits in social interaction and cognition during childhood^13^. While this underscores the neurodevelopment importance of brain connectome gradients, their potential as biomarkers in ADHD remains to be clarified .

Despite numerous potential neuromarkers for ADHD, diagnosis by means of ADHD-specific neural perturbations remains challenging, which may imply the presence of intermediate phenotypes^14^. To determine the underlying phenotypes, a growing number of studies utilized imaging transcriptomic which combines genetic variance with imaging-derived phenotypes (IDPs, e.g., connectome gradient) to improve identifying the biological pathophysiological mechanisms underlying neurodevelopmental disorders. Specifically, both neurotransmitter imbalance and their cellular characterization/structures that code vulnerability genes show specific correlations with morphometric or connectome gradient phenotypes dysfunctions in adult depression^15^ and childhood autism^16^. An initial study has reported significant association between gene-set expression levels with brain size in ADHD adolescence and adults^17^. This research although focusing on brain structural vulnerability, supports for in-depth examination of functional neuropathological underpinnings for ADHD-specific brain connectome gradient perturbations are needed.

To this end, we determined neurotransmitomic profiles and genetic risk indices for ADHD-specific IDP on brain connectome gradient perturbations, by using both imaging-transcriptomic association analysis and a knowledge-based meta-analytic decoding approach (**Fig. 1**). We firstly examined ADHD-specific-functional connectome gradient perturbations with a multisite resting-state functional MRI dataset (i.e., ADHD-200), and further decoded cognitive functions associated with gradient changes in ADHD patients. To reveal the neurotransmitomic molecular associations of these neural changes in ADHD, we capitalized on a partial least square (PLS) regression model to map a normative neurotransmitter atlas into cortical changes of connectome gradients. Further, we specified the correlation of ADHD-specific connectome gradient perturbations to microassay-determined brain spatial gene expression levels for clarifying transcriptomic signatures of IDP of ADHD. Based on meta-analytic decoding toolkit, these transcriptomic markers were decoded as risky gene sets for correlating to cognitive, neural, neurological and neuropsychiatric diseases. Finally, we performed parallel enrichment analyses to reveal gene-set-related ontological clusters and pathways for IDP of ADHD by insights into cellular diversities (i.e., cell type-specific and chromosome-enrichment). Workflow and technical details for the current study are depicted in **Fig. 1**. By means of this multi-methodal and multi-level approach, we aimed to advance our understanding of the molecular pathophysiological mechanisms of ADHD.

**Fig. 1.**
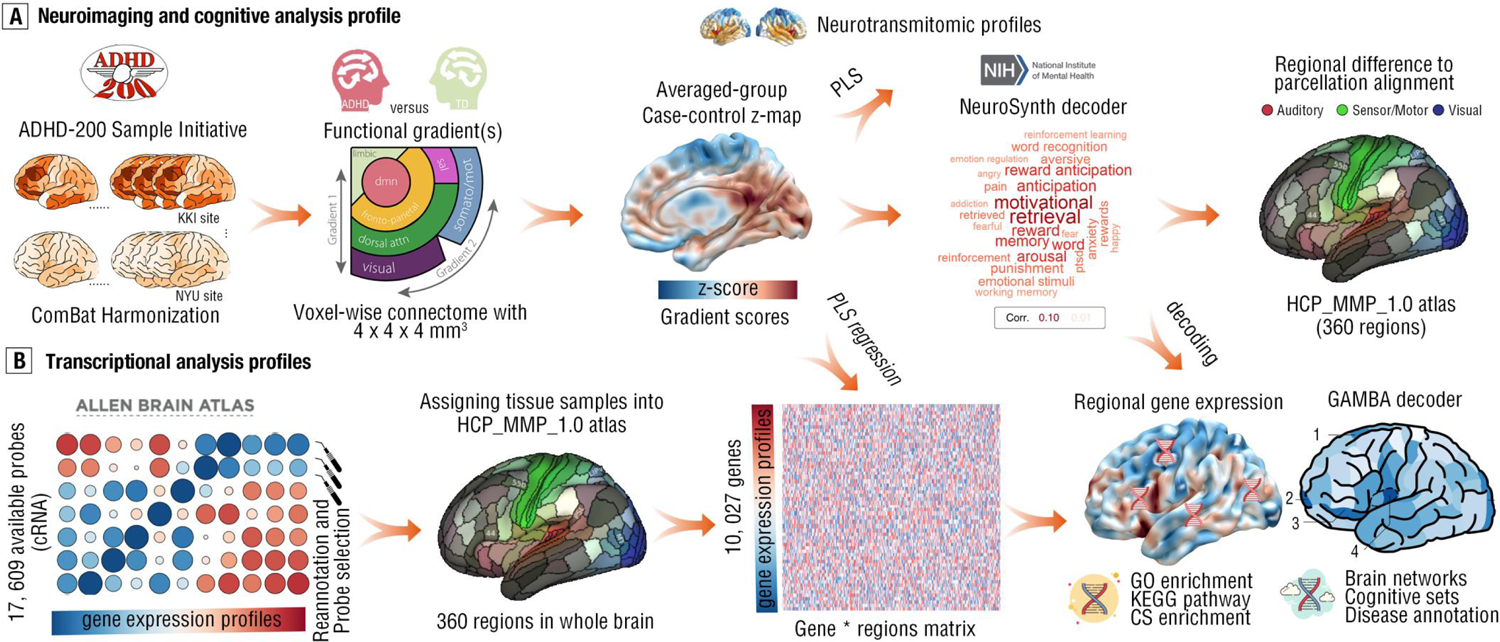
Study flowchart. **(a)** Neuroimaging and cognitive analysis workflow. We utilized high-quality resting-state functional connectivity MRI data from an open data initiative entitled “ADHD-200 Sample”, which included 93 ADHD patients with combined symptoms and 93 matched TD controls across five image sites. Further, by using the embedding mapping method, the principal and second-third gradients for each participant were estimated in voxel-wise network for global and brain subsystems. Then, general linear model was used to produce the case-control z map for vertex-wise statistics with gradient scores. To reveal the neurotransmitomic profiles of these gradient differences (i.e., case-control z map), we aligned this z map into Schaefer-100 atlas, and performed partial least square (PLS) regression fitting Hansen neurotransmitomic matrix that defined by Schaefer-100 atlas (19 x 100, with 19 receptors or transporters) for z-value vector (1x 100). To uncover the cognitive terms of such gradient differences, we decoded the case-control z map by using NeuroSynth decoder, and provided word-cloud chart here, with large font size for high correlation. Final step in neuroimaging and cognitive analysis was to parcel case-control z map (i.e., regional gradient differences) into HCP_MMP_1.0 atlas with 360 regions. **(b)** Transcriptional analysis profiles. We preprocessed gene expression level from AHBA (Allen Human Brain Atlas) dataset by canonical pipelines (https://github.com/BMHLab/AHBAprocessing), such as reannotation and probe selection from samples. Then, the HCP_MMP_1.0 atlas was used to assign samples into whole brain with 360 regions, and finally produced gene x region matrix. Likewise, the PLS regression model was used to fit this matrix for case-control z-value vector so as to reveal the regional gene expression pattern of ADHD-specific gradient changes. Further, gene ontology (GO) enrichment and Kyoto Encylopedia of Gene, Genomes (KEGG) pathways and chromosome (CS)-enrichment were estimated for these transcriptional signatures. In addition, the GAMBA (Gene Annotation by Macroscale Brain-imaging Association) decoder was used to reveal the associations of brain networks, cognitive components (sets), and diseases for these transcriptional signatures.

## MATERIALS AND METHODS

### Participants and images preprocessing

We capitalized on an ADHD cohort from a high-quality open data-sharing initiative entitled ADHD-200 Sample (http://fcon_1000.projects.nitrc.org/indi/adhd200/index.html). To stringently control confounding factors (e.g., diagnostic criterion, imaging quality, disease subtypes and comorbidity), we included pharmacological treatment-naive children and adolescents who were diagnosed for ADHD symptomatology using validated criteria such as DSM-V only after visual inspections on image quality (Supplemental Method 1). The final sample included 94 ADHD patients and 94 matched typically developing (TD) control individuals (**Tab. 1**, Supplemental Method 1 and **Figure S1-2**). Ethics approval was granted in the context of the original studies and further ethics, and thus were waived by the IRB of Third Military Medical University (PLA, China) given that no original human data.

**Tab. 1.**
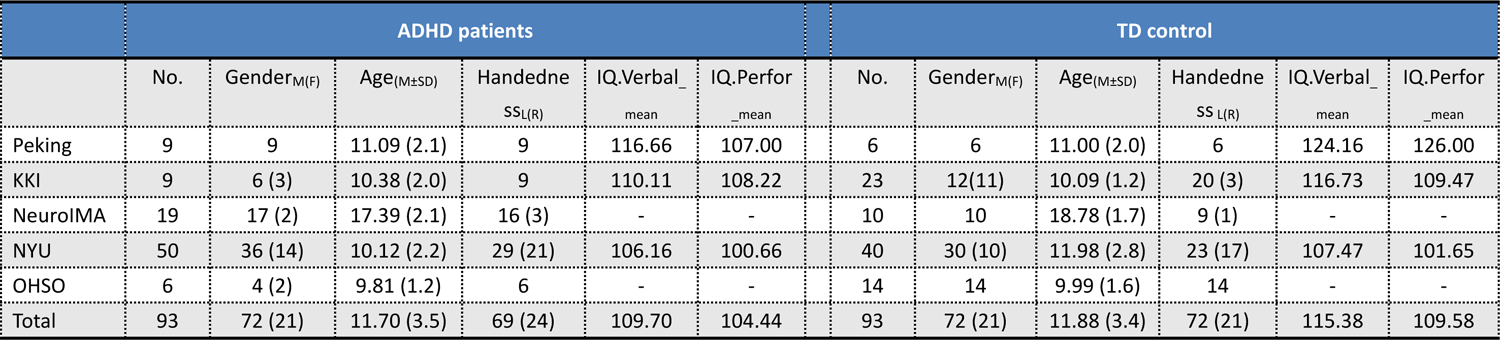
Demographic characteristics for ADHD patients and TD control individuals. The number of (No.) males is provided in the table, alongside with the number of females. Mean value for age has been reported in the table, alongside with standard derivation (S.D.). Handedness describes the number of Left-handedness, alongside with right ones. Verbal IQ and IQ performance were tested by Wechsler Intelligence Scale for Children, Fourth Edition (WISC-IV). “-” means missing data in the original records.

Image acquisition have been performed by five independent sites in final sample (i.e., Peking, KKI, NeuroIMAGE, NYU and OHSO). Protocol for scanning have been detailed into the Supplementary Data 1. Image preprocessing was followed by NeuroBureau NIAKPipeline^18^. Full details for image preprocessing can be found in Supplement Method 2.

### Connectome gradient estimates

To estimate the hierarchy of cortical organization, we estimated the voxel-wise “connectopies”, which capture gradient(s) over high-dimensional spaces, rather in a discrete parcellation atlas^9, 19^. We firstly downsampled original preprocessed images into 4 x 4 x 4 mm^3^ resolution to balance computational requirements leading to 18,933 voxels (nodes) for each participant^20^. Then, inter-correlationsbetween each pair of these voxels were estimated. Subsequently, we included the top 10 % connections for each voxel. For building connectomes, cosine similarity was estimated for each pair of selected voxels mentioned above to generate similarity matrix. Each resultant matrix was further scaled and normalized into angle matrix^21^. For gradient detection, we capitalized on the diffusion map embedding method to examine gradient(s) that could reflect the hierarchical functional connectivity pattern of the functional connectome. Full details can be found in the Supplemental Method 3. Given the multi-site heterogeneity, we carried out ComBat harmonization (https://github.com/Jfortin1/neuroCombat) to correct the potential noise for statistics with respect to connectome gradient.

### Neurotransmitter-distribution signatures

By collating multi-site positron emission tomography (PET) data in a large-scale dataset (n = 1,200), Hasson et al (2022) provided a neurotransmitter atlas to quantify 19 neurotransmitters (receptors and translators) distribution^22^. This work provided the neurotransmitter atlas for distribution density of 19 receptors/translators by using Schaefer-100 cortical parcellation (100 nodes), and we thus obtained a 19 x 100 neurotransmitter matrix. Then, the voxel-wise case-control z-maps for gradient scores in each gradient axis (i.e., gradient 1, 2, 3) were separately aligned to the Schaefer-100 atlas. In this vein, a IDP vector was generated with a size of 1 x 100. We utilized partial least squares (PLS) regression models for fitting the neurotransmitter distribution (density) to regional IDP differences (z-map for connectiome gradient herein), with 19 x 100 matrix for predictors and with 1 x 100 vector for response (Supplemental Methods 4). PLS components would be captured by the sets of predictors with highest explanation ratio to response (e.g., PLS1, PLS2).

### IDP-transcriptomic signatures

#### Regional gene expression

The AHBA (Allen Human Brain Atlas) dataset provides a powerful and broad-certified benchmark to link the IDP-transcriptional signatures. This dataset encompasses an atlas for regional gene expression from six postmortem brains ( https://human.brainmap.org/static/download). By pre-processing benchmark^23^, the gene-atlas matrix (10,027 genes x 280 regions) was generated for IDP transcriptional analysis (Supplemental Method 5). To reveal IDP transcriptional signatures, the PLS model was used to fit gene expression level (10,027 x 280) to regional z-map vector that had been aligned into HCP_1.0 atlas (1 x 280). The bootstrapping processes were used for statistical inferences. The gene sets were divided into PLS+ and PLS-for multiple comparison correction (Bonferroni-Hochberg correction at p < .001) (Supplemental Method 5).

#### Transcriptomic-IDP decoding and neuropsychiatric/neurological decoding at GAMBA

GAMBA (Gene Annotation by Macroscale Brain-imaging Association) is a pathophysiological database for online meta-analytic gene-list annotation^24^. The genes that survived after Bonferroni-Hochberg correction in the PLS set were assigned into two subset, with one for positive weights (PLS+) and another one for negative weights (PLS-). We used general linear regression model to fit the PLS+ and PLS-for the neuroimaging-derived regional/global phenotype, respectively, which included resting-state brain network, cognitive brain components, cortical metabolism and evolutionary cortical expansion. In addition, by using activation-likelihood estimate (ALE) analysis, the neuropsychiatric/neurological association of these IDP-transcriptomic gene sets was examined at GAMBA as well. Beta coefficient (β_1_) for each model was standardized, while the corresponding p value was corrected by Bonferroni-Hochberg method at alpha = 0.05 (Supplemental Method 6).

#### Enrichment analysis

To reveal the biological process and potential pathways, we utilized an online meta-analytic tool entitled “Metascape” for enrichment analysis of these transcriptomic-IDP gene sets (https://metascape.org/gp/index.html)25. We sorted gene sets into PLS+ and PLS-component, respectively, which contained genes that survived from Bonferroni-Hochberg correction at p < .001. Then, both components were subjected into Metascape for examining which pathways were enriched for this gene list (false-discovery error (FDR) correction at p < .05). In addition, to understand whether the IDP-transcriptomic gene lists showed pathways for known polygenic risky genes, we reviewed recent genome wide association study (GWAS) and extracted common genes that reached significance level from recent didactic reviews^26–28^. Further, we inputted these risky gens as comparable list for additional multi-gene-list enrichment analysis^25^ to uncover the shared- and specific-pathway or gene ontology (GO) clusters (FDR correction at p < .05).

#### Cell type-specific and chromosome-IDP-transcriptomic signatures

To probe into the cell type-specific genes, we compiled cell type data from single-cell studies that used postmortem cortical samples and a following didactic pipeline^29, 30^. Genes that survived after correction in PLS+ loading were assigned into cell-type atlas, and were counted for the overlapping from difference cell types as “true effect”. Further, the pseudo-gene-list was generated by randomly selecting genes from the whole gene pool (10,027 gene) with the same number of this gene set (i.e., these survived genes in PLS+). Then, the overlapping counts in each cell type were recorded as “null effect”. Likewise, the median rank-method^30^ was used to examine how this gene set that survived after Bonferroni-Hochberg correction in PLS+ component enriched for chromosome profiles, with actual median rank for “true effect”. This permutation process has been iterated for 5,000 to generate the null distribution for each cell type or for each chromosome. Finally, the statistical inference could be made by inspecting whether the point estimate of “true effect” fall outside of the null distribution at 5^th^ or 95^th^ centile (i.e., p value for permutation < .05 at n = 5,000).

#### Replication analysis

We replicated the main analysis in an independent small dataset to validate the reproducibility of image-derived phenotype (IDP) (n = 22). The Jaccard index was used to quantitatively estimate the consistency of all the results in validation dataset by referring from main findings. Details have been documented in the Supplemental Result 17, Figure S13 and Table S30-35.

## RESULTS

### Connectome gradient patterns of ADHD

#### Global topology of gradients

We capitalized on the diffusion map embedding method to capture connectome gradient patterns in both groups (Supplemental Result 1 and **Figure S3**). Principal gradients 1-3 showed the prominent posterior-unimodal-anterior-multimodal, association-cortex and unimodal hierarchy in both cohorts, respectively (**Fig. 2A-B**, Supplemental Result 1 and **Figure S4-5**). However, no statistical differences for global topology were found.

**Fig. 2.**
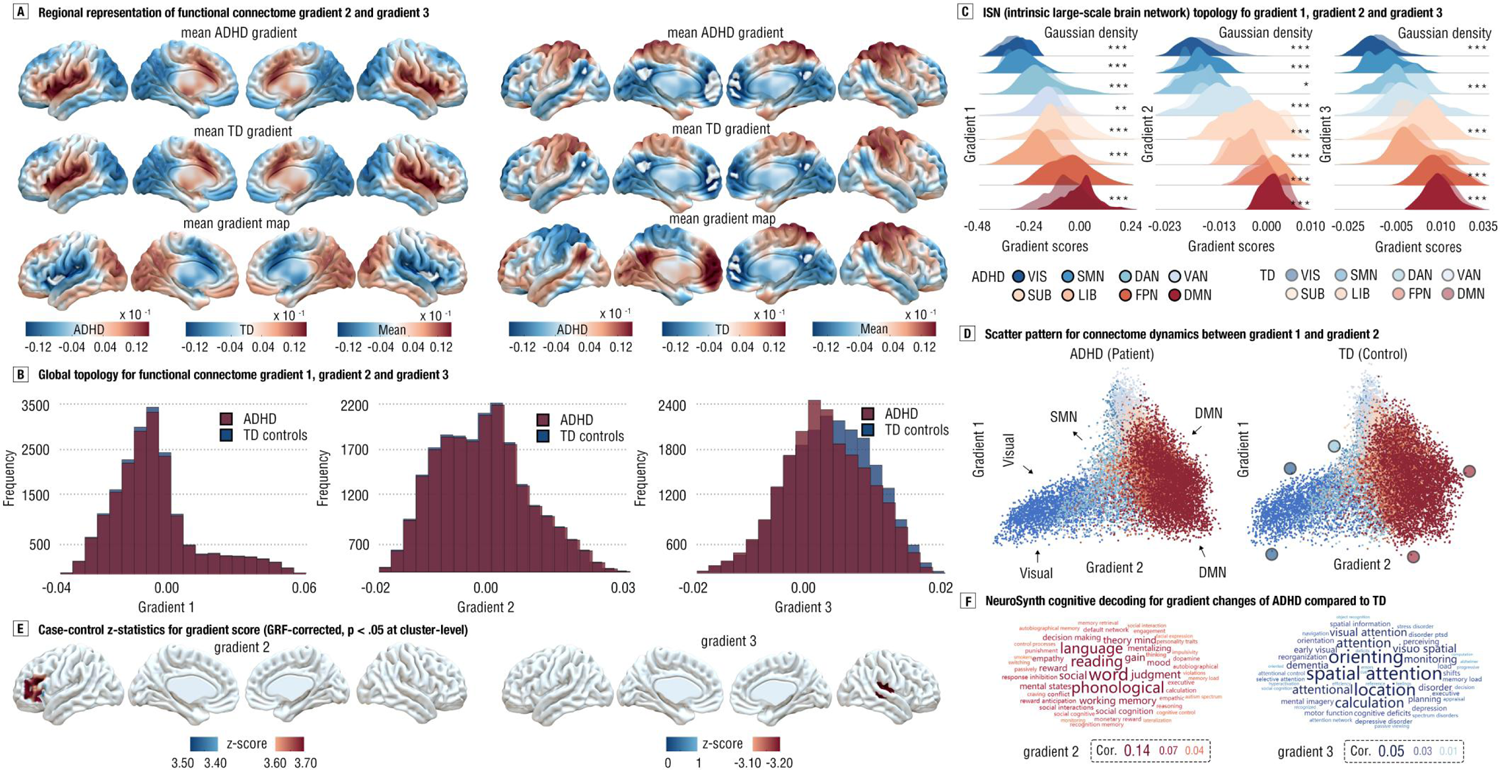
Gradient differences and their cognitive terms from ADHD to TD controls. **(a)** Averaged-group gradient maps for second (gradient 2, left panel) and third gradient (gradient 3, right panel) in ADHD, TD and all the participants. Colorbars represented gradient scores. **(b)** Global topology for functional connectome for principal gradient (1) and gradient 2-3. Dark red means ADHD, while dark blue means TD controls. Axis-X indicated gradient scores. **(c)** ISN (intrinsic large-scale brain network) topology for principal gradient and gradient 2-3. All the panels showed the density plots for corresponding gradient scores from eight ISNs, with Gaussian kernel for estimating density for each gradient scores distribution, including visual network (VIS), sensory/motor (SMN), dorsal attention network (DAN), ventral attention network (VAN), subcortical network (SUB), limbic network (LIB), frontoparietal network (FPN) and default mode network (DMN). K-S two sample-test was used to examine the distribution differences between ADHD and TD, with *** for p < .001, with ** for p < .01. (d) Scatter pattern for connectome dynamics between gradient 1 and 2. Colors for each dot were labeled as section C, and the circles for the TD scatter plot indicated the largest differential voxels for gradient scores compared to ADHD in specific brain subsystems. (e) Case-control z-map for gradient score (GRF-corrected, p < .05 at cluster-level). **(d)** NeuroSynth cognitive decoding for gradient changes of ADHD compared to TD, with large font size for high correlation strength.

#### System-based gradient perturbation

In the principal gradient, we found less gradient scores in dorsal attention network (DAN, Cohen d = −0.11, p < .0001), ventral attention network (VAN, Cohen d = −0.08, p < .001), limbic network (LIB, Cohen d = −0.41, p < .0001), default mode network (DMN, Cohen d = −0.11, p < .0001) and subcortical network (SUB, Cohen d = −0.10, p < .0001) in ADHD, yet higher gradient scores in visual network (VIS, Cohen d = 0.21, p < .0001) and sensory/motor network (SMN, Cohen d = 0.16, p < .0001) in ADHD (**Fig. 2C-D**, Supplemental Result 2 and **Table S1**). Instead of a global gradient shift, these results may reflect dysregulations in hierarchical architectures of the brain’s attention systems and unimodal-multimodal switching system in ADHD (Supplemental Result 2 and **Table S2-3**).

#### Vertex-level gradient perturbation

The vertex-based analysis showed higher gradient scores in the left lateral orbitofrontal cortex (LOFC) in ADHD (p_(cluster-wise)_ < .05, GRF-corrected). In the third gradient, the lower gradient scores were observed for ADHD patients in a cluster spanning superior temporal areas (p_(cluster-wise)_ < .05, GRF-corrected) (**Fig. 2E**). In sum, ADHD patients may present regional similarity perturbations in prefrontal and temporal connectome profiles.

#### Cognitive decoding for ADHD gradient perturbations

By decoding the regional gradient perturbation maps in NeuroSynth that implementing the online meta-analytic activation likelihood estimate (ALE) to behavioral-brain association in the database, we found changes in the principal gradient 1 for ADHD patients were involved into episodic memory and retrieval process (Supplemental Result 3 and **Figure S7**). Further, cortical perturbations in the second-third gradient were mainly correlated with functions related to language or spatial attention, respectively (**Fig. 2F**).

#### Neurotransmitomic signature for gradient-derived phenotype of ADHD

We complied a neurotransmitomic atlas to map gradient changes of ADHD by a standard pipeline (Hansen et al., 2022) (Supplemental Result 4 and **Table S4**). By using PLS regression model, the PLS1 neurotransmitter-distribution map presented the prominent posterior-anterior hierarchy (**Fig. 3A**). Further, to probe into the roles of specific neurotransmitomic receptors on connectome dysfunctions in ADHD, the GABA_A/BZ_ and 5-HT_2A_ receptors that significantly explained model variances were selected from PLS1 component (p < .001; Z > 4.2 or Z < −4.2, BH-corrected; **Fig. 3B**). It was found that the spatial distributions of them were significantly associated with changes of gradients for ADHD (r_GABAA_ = .27, p < .01; r_5-HT2A_ = .25, p < .01, spatially-corrected; **Fig. 3C**; Supplemental Result 5, **Figure S8-9** and **Table S5-8**). Thus, we may infer that the GABA_A/BZ_ and 5-HT_2A_ were over-distributed as changes of principal gradient architecture in ADHD patients.

**Fig. 3.**
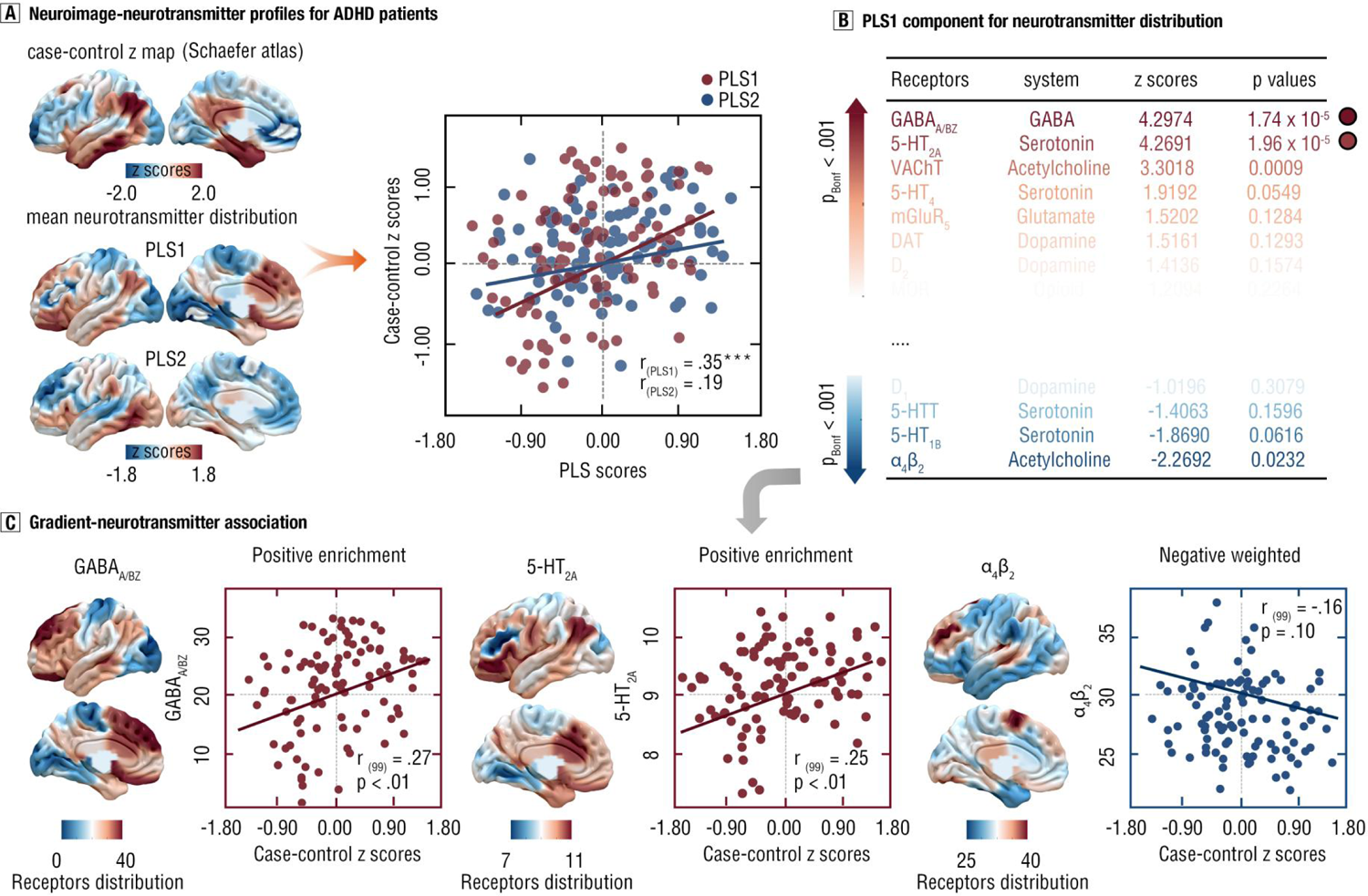
Neurotransmitomic signatures of gradient-derived differences for ADHD. **(a)** Neuroimage-neurotransmitter profiles for ADHD patients. By using Schaefer-100 atlas, the whole-brain case-control z map for principal gradient has been aligned into 100 cortical regions. Further, By using the PLS regression model, we revealed the PLS1 and PLS2 components (p <.01) to predict case-control gradient differences (z-values). Thus, brain maps for PLS1 and PLS2 were presented z-transformed PLS scores. Right panel showed the scatter plot for the correlations between case-control differences and PLS scores. **(b)** PLS1 component for neurotransmitter distribution. In the PLS1 component, we used Bootstrapping methods to estimate the weights (z scores) for each neurotransmitter (a total of 19). By correcting p values for each one with Bonferroni-Hochberg correction at < .001, the GABA_A/BA_ and 5-HT_2A_ receptors were survived. **(c)** Gradient-neurotransmitter association. The receptor distribution for GABA_A/BA_ and 5-HT_2A_ were shown by Schaefer atlas, and corresponding scatter plots were provided. Scatter plot for most negative weighted receptor has been provided as well.

### Transcriptomic signature for gradient-derived phenotype of ADHD

#### Regional gene expression pattern of gradient perturbation

By utilizing AHBA atlas and PLS models (see Supplemental Method 5), we found that top two PLS components (i.e., PLS1 and PLS2) explained 17.84 % variance of ADHD-specific gradient changes for principal gradient (p < .001, spatially-corrected). PLS1 map presented a prominent top-down hierarchy, indicating a downstream transcriptomic trend for ADHD-specific perturbation in principal gradient (**Fig 4A**). PLS2 map illustrated a dominant anterior-posterior hierarchy, with low gene expressions in prefrontal areas and with high gene expression in posterior sensory and visual system (**Fig. 4A**). Supporting that, we also found the statistically correlations of ADHD-specific principal gradient changes to both PLS1 and PLS2 scores (r _PLS1_ = .372, p < .001, r _PLS2_ = .352, p < .001, spatially-corrected). Thus, the principal gradient changes of ADHD may be also the function of transcriptomic dynamics, especially in the top-down and anterior-posterior gene expression patterns.

**Fig. 4.**
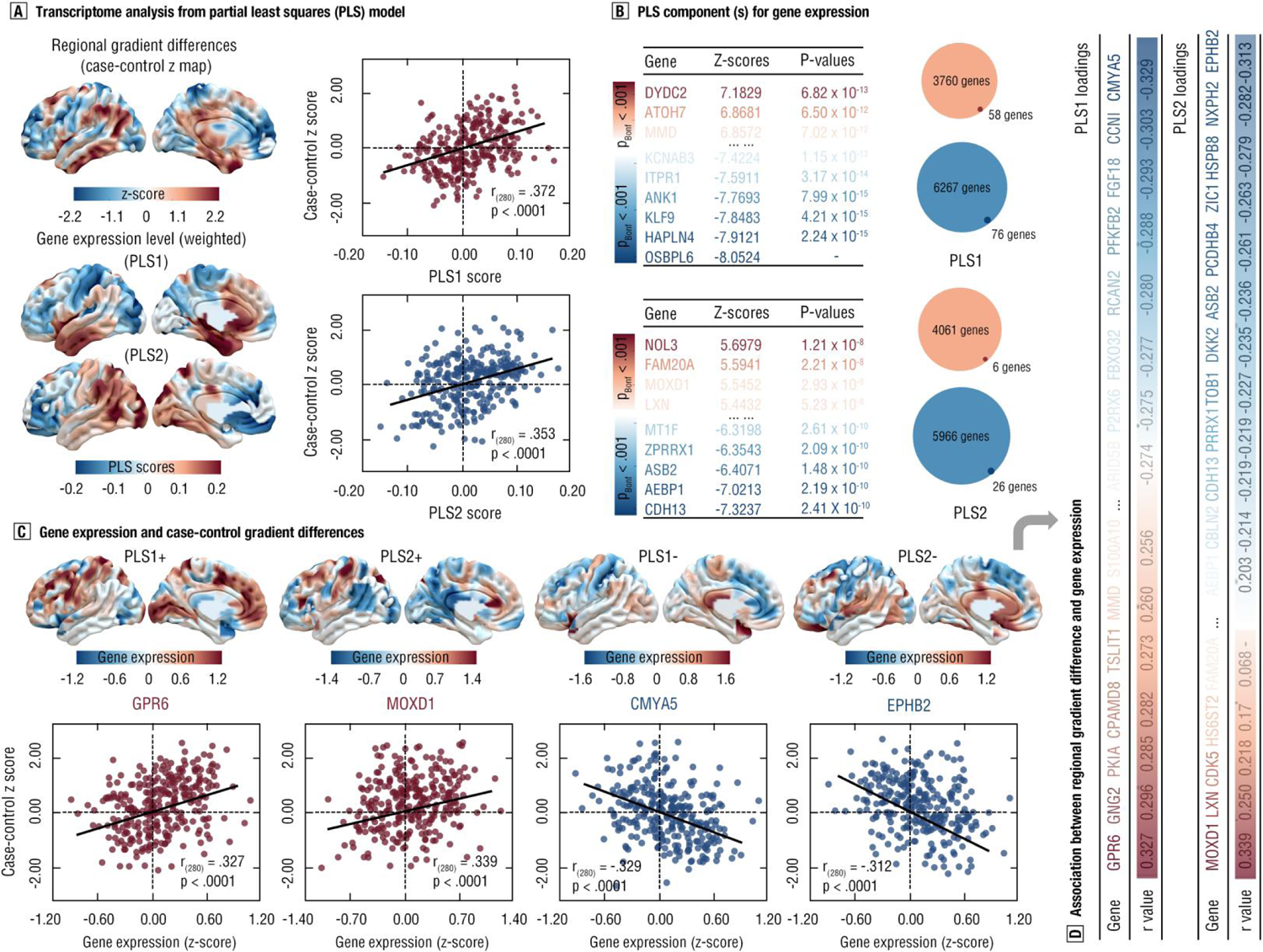
Transcriptomic profiles of gradient-derived differences for ADHD. **(a)** Transcriptome analysis from partial least squares (PLS) regression model. Regional gradient differences were aligned into HCP_MMP_1.0 atlas. Further, PLS1 and PLS2 scores (i.e.,g weighted gene expression level) were presented into HCP_MMP_1.0 atlas here. Corresponding scatter plots were provided in this right panel. **(b)** PLS components for gene expression. we used Bootstrapping methods to estimate the weights (z scores) for each gene in PLS components (PLS1 and PLS2). Then the weights were partitioned into positive and negative sets, i.e., PLS1+, PLS1-, PLS2+, PLS2-. After multiple comparison correction (Bonferroni-Hochberg correction at < .001), 58 genes (76 genes) survived from 3760 gene (6267 gene) sets in PLS1; 6 genes (26 genes) survived from 4061 genes (5966 genes) in PLS2. In the right panel, the pie chart showed relative ratio of selected genes from total candidates in corresponding PLS components, with red for positive weight component and with blue for negative weight component. **(c)** Gene expression and case-control gradient differences. We extracted gene expression level for these selected genes from PLS components, and illustrated scatter plots for each PLS component that showing the largest correlation strengths between this given gene and case-control difference. **(d)** Association between regional gradient difference and gene expression. Correlations for expression levels of all the genes and case-control differences were calculated, and were presented in this chart with descending order.

Further, by using Bootstrapping method, 58 genes (e.g., DYDC2, ATOH7) were revealed to over-expressed with principal gradient changes in the PLS1+, whilst 76 genes (e.g., OSBPL6, HAPLN4) were found to be under-expressed as well (PLS1-; **Fig. 4B**, Supplemental Result 6 and **Table S9-10**). Results for PLS2 can be found at the Fig. 4B, Supplemental Result 6 and Table 11-12. Also, we found the statistically significant positive correlation between principal gradient changes of ADHD and regional gene expressions of DYDC2 (top-expressed single gene in PLS1+; r = .327, p < .001, spatially-corrected) and NOL3 (top-expressed single gene in PLS2+; r = .339, p < .001, spatially-corrected; **Fig. 4C-D**, Supplemental Result 8 and **Table S13-14**). Results for PLS2 could be found in Table S15-16. To examine the specificity of these risky genes, we further correlated In Situ Hybridization (ISH) genes for other neurodevelopmental disorders to such gradient changes in ADHD. No statistically significant associations between them were found (Supplemental Result 8 and **Table S17**). In sum, these findings may reveal the specific transcriptomic signatures of principal gradient changes in ADHD.

#### Transcriptomic decoding at Gene Annotation by Macroscale Brain-imaging Association (GAMBA)

To decode ADHD-specific PLS components, we performed online meta-analytic image-related decoding for these PLS-gene lists by using GAMBA. We found that the visual networks were negatively (positively) predicted by gene list in the PLS1+ (β = −.50) and PLS1-(β = −.49, all p < .05 with Bonferroni-Holm FDR correction) (**Fig. 5A**). In addition, the limbic network was revealed to be positively (negatively) predicted by gene lists in the PLS1+ (PLS1-) (β = .58, β = −.55, all p < .05 with Bonferroni-Holm FDR correction) (**Fig. 5A**, Supplemental Result 9, and Table S18). Full results for PLS2 component have been sorted into the **Table S19**.

**Fig. 5.**
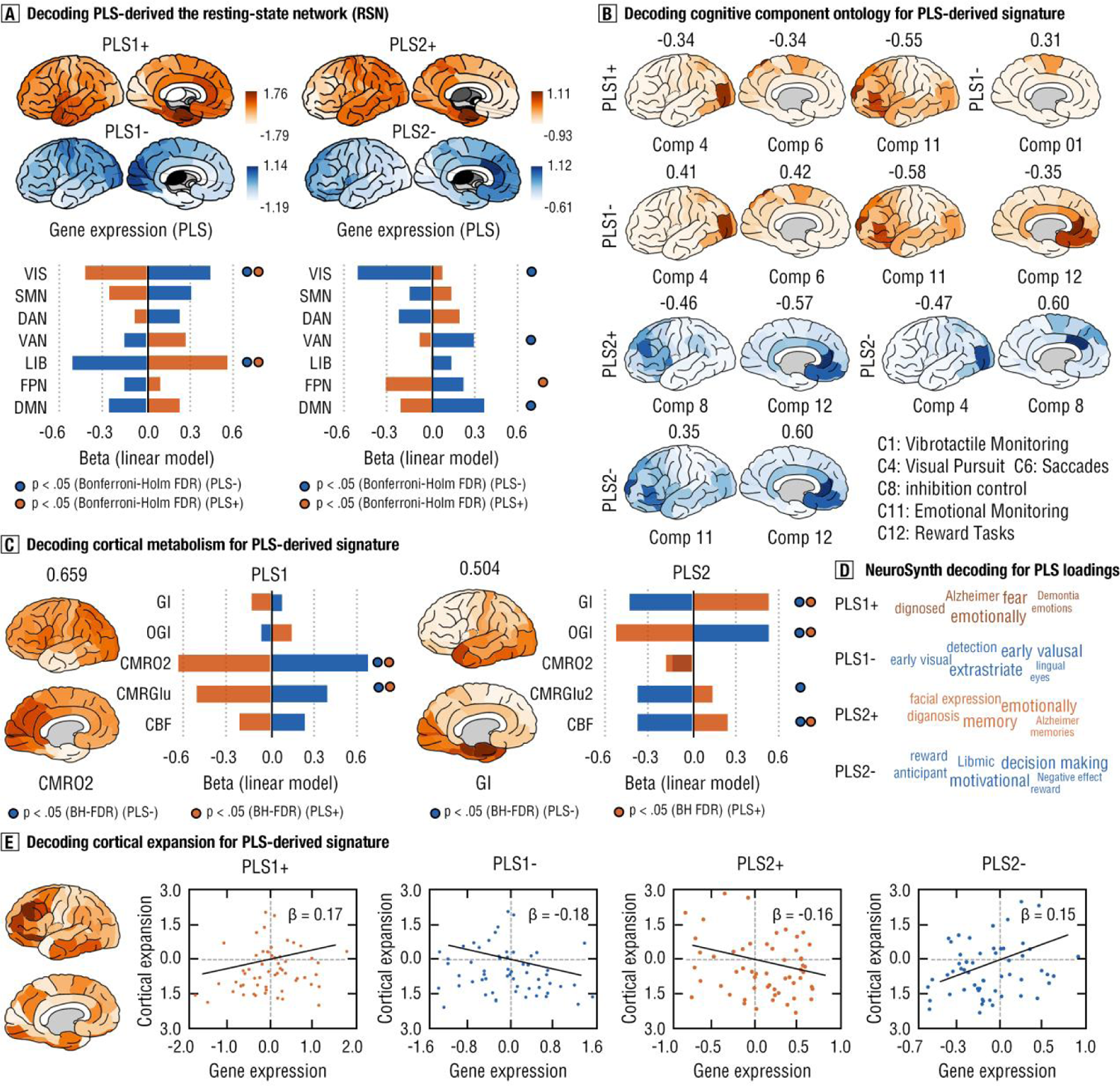
Transcriptomic decoding at Gene Annotation by Macroscale Brain-imaging Association (GAMBA). **(a)** Decoding PLS-derived the resting-state network (RSN). We decomposed PLS components that detected from transcriptomic analysis into PLS1+, PLS1-, PLS2+ and PLS2-. By using the GAMBA decoding, the gene expression plots were provided by using D-K atlas. The linear regression model was used to fit the expression levels to the network property of RSNs, and the standardized beta coefficient was presented by bar plots. Dark orange (PLS+) and blue (PLS-) indicated the p value for this beta coefficient reached statistical significance level (p < .05 at Bonferronni-Holm FDR correction). **(b)** Decoding cognitive ontology for PLS-derived signatures. Statistics are comparable with section (a). Brain maps were illustrated by the beta coefficients from linear models. The beta coefficient for corresponding cognitive component (ontology) have been labeled at the head of brain map. The beta coefficients were presented only if their p values reached statistical levels. Specific functions for cognitive components (ontology) have been embedied in this chart. **(c)** Decoding cortical metabolism for PLS-derived signatures. Statistics, brain maps and labels are comparable with section (b). Glycolytic index = GI; oxygen-glucose index = OGI; cerebral metabolic rate of oxygen/glucose = CMRO2/GMRGlu; cerebral blood flow = CBF. **(d)** NeuroSynth decoding for PLS loadings. We decoded the cognitive terms of PLS components at NeuroSynth, respectively, with large font size for high correlation strengths. **(e)** Decoding cortical expansion for PLS-derived signatures. Brain map showed the beta values from the regression model of gene expressions to cortical expansion. Due to the lack of regional cortical expansion data in right hemisphere, only left one with 57 regions in D-K atlas was used for regression model.

Furthermore, gene list in the PLS1+ could negatively predict brain cognitive component 4, 6, and 11, which indicated the over-expressed gene set may disrupt brain cognitive functions of visual and emotional monitoring (**Fig. 5B**, Supplemental Result 10, and **Table S20-21**). Gene list in the PLS1+ (PLS1-) components were demonstrated to positively (negatively) predict both cortical metabolic rate for oxygen (CMRO^2^) and for glucose (CMRGlu)(**Fig. 5C**, Supplemental Result 11, and **Table S22-23**). In this vein, it may imply a potential moderator of gradient-derived gene expression on cortical metabolisms for ADHD patients. In addition, by using NeuroSynth database, we found that the PLS1+ gene set was positively associated with cognitive terms of emotions, Alzheimer and fear (**Fig. 5D**, Supplemental Result 12, and **Figure S10**). No statistical correlational associations were found in decoding cortical expansions (**Fig. 5E**). In total, the transcriptomic profiles of gradient-derived phenotype for ADHD may be associated with the perturbations in the brain visual/limbic networks, visual/emotional monitoring functions and cortical metabolisms.

#### Transcriptomic decoding neuropsychiatric or neurological diseases at BrainMap dataset

We further decoded these PLS gene sets for associating neuropsychiatric or neurological diseases at online BrainMap dataset (**Fig. 6A**). In the voxel-based morphology (VBM) dataset, the PLS2 component robustly predicted cortical structural deficits with obsessive-compulsive disorder (OCD), bipolar disorder (BP), schizophrenia, depression and anxiety (**Fig. 6B**, Supplemental Result 13, and **Table S24**). In the functional MRI dataset, PLS1 robustly predicted cortical functional dysfunctions in Asperger, autism spectrum disorder (ASD), Dyslexia, Parkinson and stroke (**Fig. 6B** and **Table S24**), whilst PLS2 could robustly predict post-traumatic stress disorder (PTSD), depression, ADHD, anxiety, mild cognitive impairment (MCI) and bipolar disorder (BP) (**Fig. 6B** and **Table S25**). Taken together, in addition to ADHD, this gradient-derived transcriptomic profiles of ADHD may be associated with brain structural or functional disruptions in broad psychiatric and neurological diseases.

**Fig. 6.**
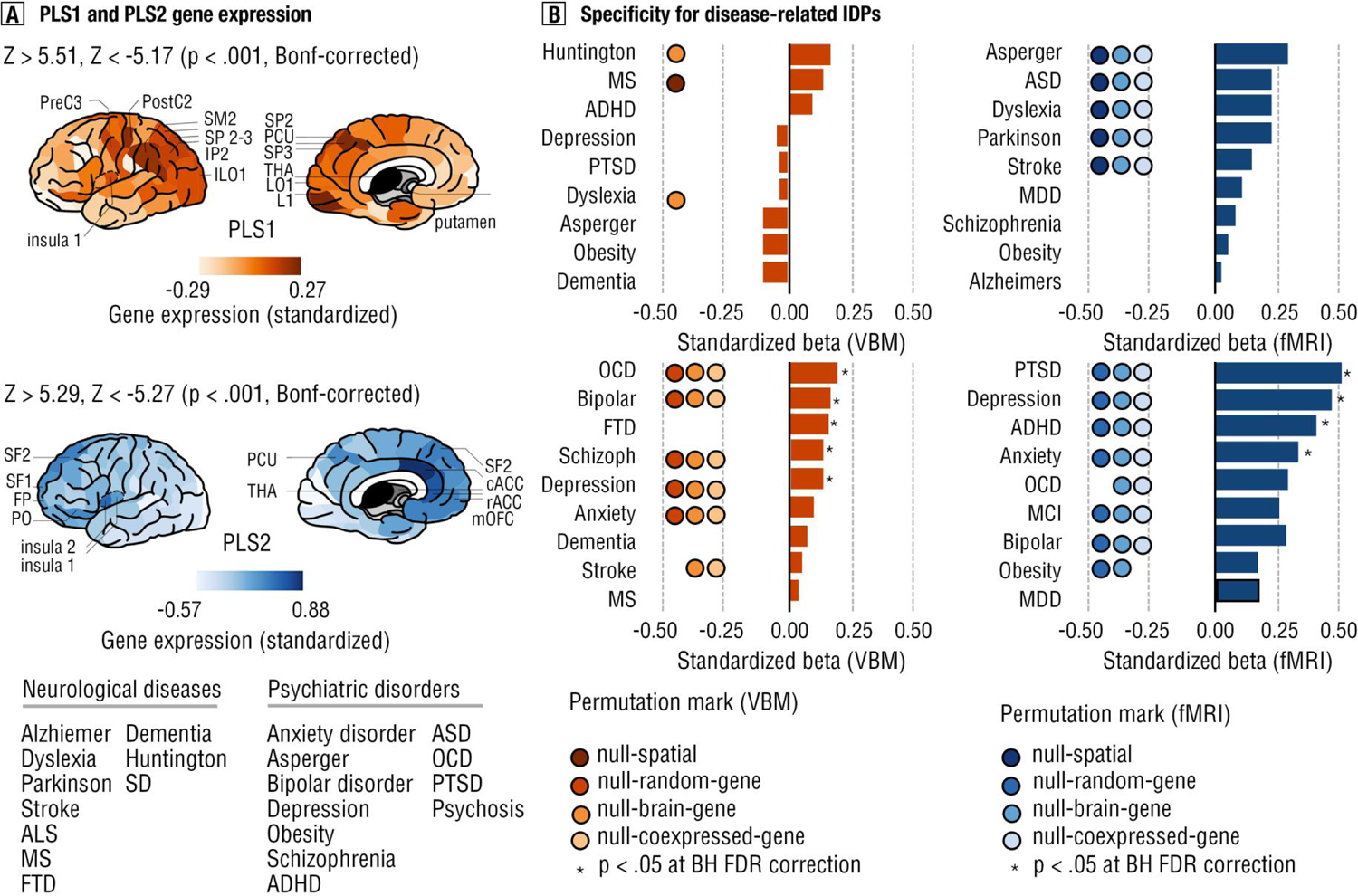
Psychiatric and neurological conditions decoding. **(a)** PLS1 and PLS2 gene expression. We generated PLS gene sets that reached statistical significance level (Bonferroni-Hochberg correction at < .001) into PLS1 and PLS2 components, and decoded them into BrainMap dataset. Brain map showed the standardized expression level pattern for PLS1 and PLS2 component from D-K atlas, respectively. Top 10 regions that show the highest expression levels were labeled. In the bottom chart, neurological disease and psychiatric disorders that the BrainMap dataset allowed have been given. **(b)** Specificity for disease-related image-derived phenotype (IDPs, i.e., gradient differences for ADHD patients herein). The linear regression models were built to fit gene expression of these gene sets (PLS1 and PLS2) to brain dysfunctions of these disease or disorders. P values were corrected by Bonferroni-Holm FDR corrections. To examine specificity for these results, the multiple permutation methods with 5, 000 iterations were used.

#### Enrichment clusters and pathways of transcriptomic signatures for ADHD

By using Metascape tool (https://metascape.org/gp/index.html#/main/step1), we found significant gene enrichment in GO biological processes, such as “regulation of secretion”, “regulation of chemotaxis”, “synaptic vesicle cycle”, and “negative regulation of locomotion”(**Fig. 7A**, Supplemental Result 14, and Table S26). Further, the KEGG pathway of alcoholism was revealed to be enriched as well. In the protein-protein interaction (PPI) network, the “muscle contraction” and “calcium signaling pathway” components were uncovered (**Fig. 7B**). Full results for enrichment analysis in PLS1 have been documented at Supplemental Result 14, **Table S27**, and **Figure S11**.

**Fig. 7.**
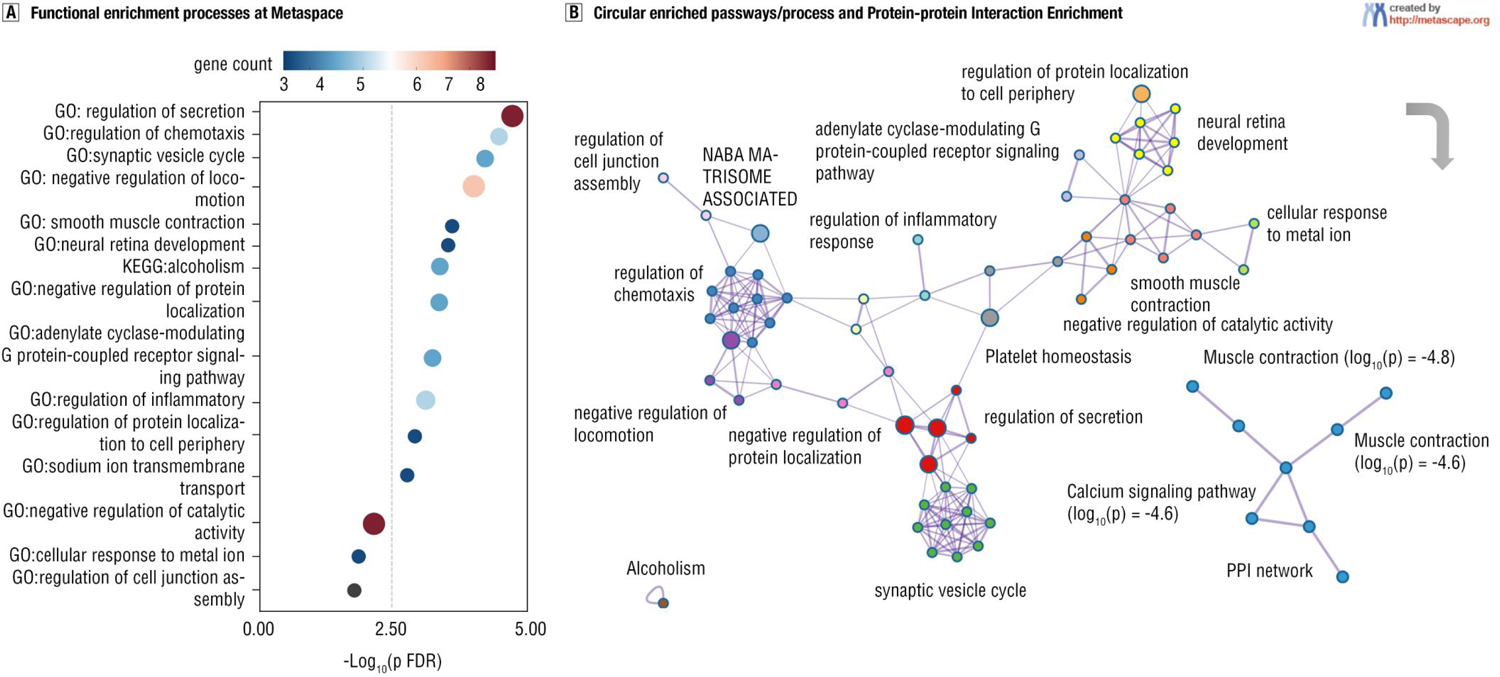
Enrichment analysis and protein-protein interaction. **(a)** Functional enrichment process at Metascape. All the gene ontology (GO) and Kyoto Encylopedia of Gene, Genomes (KEGG) pathways were reported here only if they survived from FDR correction at p < .05. **(b)** Circular enriched pathways/process and protein-protein interaction (PPI) enrichment. GO or KEEG pathways have been labeled at corresponding locations, and the PPI network has been embedied into the right-bottom corner. The network is visualized using Cytoscape from the Metascape.

To further examine the association of this PLS1+ gene set to canonical polygenic risks of ADHD^28, 31^ that tested from genome-wide meta-analysis studies (GWAS), we performed multi-gene-list enrichment at Metascape as well. It was found that they shared only six gene ontology terms in enrichment pathways (**Fig. 8A**). Likewise, for PLS1 component, we found that only three genes shared the same ontology terms between multi-gene-lists. Full results have been reported in the Supplemental Result 15, Table S28 and Figure S12. Thus, we may provide additional gene-related transcriptional signatures for ADHD.

**Fig. 8.**
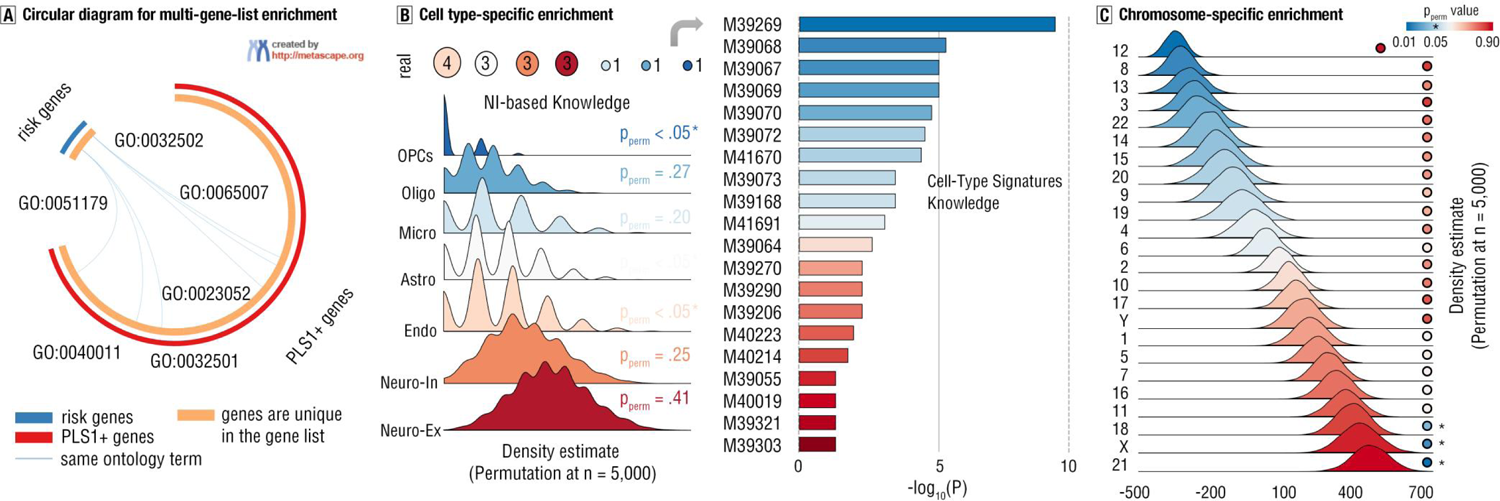
Multi-gene-list, cell type-specific and chromosome-specific enrichment. **(a)** Circular diagram for multi-gene-list enrichment. Blue blocks means the risky genes that we extracted from canonical GWAS studies, while the red outside circle indicate the PLS1+ gene sets. Orange circles mean the genes that are unique in their gene lists. Six pairs of genes were found to be categorized into the same ontology terms, and were connected by light blue lines. IDs of nntology terms have been given. **(b)** Cell type-specific enrichment. The left panel illustrated the density maps based on Gaussian kernel for canonical seven cell types from neuroimaging-based knowledge by using permutation method at n = 5, 000. The real numbers of overlapping of PLS1+ gene list to corresponding cell types were provided at head of these density plots. In the right panel, the Cell-Type Signatures Knowledge dataset was used to examine the cell type-specific enrichment for PLS1+. IDs for these cell types that survived from multiple comparison correction (FDR at p < .05) in this datasets have been provided. **(c)** Chromosome-specific enrichment. We used the median rank method to whether the chromosome-profiles (1:22, X, and Y) were enriched from PLS1 components by using permutation method at n = 5,000. * p < .05.

#### PLS-gene-list enrichment at cell type-specific and chromosome-levels

To delve into the cellular functions of PLS-gene-list enrichment across the brain, we estimated the cell type-specific enrichment for PLS1+ by using both neuroimaging-based knowledge (19) and cell-type signature knowledge^30, 32^. By building null distribution from permutation test (n = 5, 000), the oligodendrocyte precursors (p_perm_ < .05) and endothelial cells (p_perm_ < .05) were enriched from PLS1+ gene list (**Fig. 8B**). Further, by analyzing cell-type signatures of PLS1+ gene set in the Metascape dataset, we found 20 cell-specific signatures to be enriched, such as “HU FETAL RETINA RGC” and “MANNO MIDBRAIN NEUROTYPES HDA1” (**Fig. 8B**, Supplemental Result 16, and **Table S29**).

Given the biological complexity from gene expression, we further probed into whether the PLS1+ gene list enriched at chromosome-level. Here, we capitalized on the median rank-based method to assess enrichment of the neuroimaging-derived PLS1+ component to chromosome-types of 1:22 and XY (19,39)^30^. We found that the PLS1+ genes over-expressed (enriched) at chromosome 18 (p_perm_ < .05), chromosome 19 (p_perm_ < .05) and chromosome X (p_perm_ < .05) (**Fig. 8C**). In this vein, it may extend additional avenue for understanding pathological process of ADHD by perspectives from cell-specific and chromosome-variants.

#### Replication results

In total, main results for the gradient perturbations of ADHD have been majorly validated in replication sample, including globe-based, system-based and vertex-based properties (Supplemental Result 17, **Figure S13**, and **Table S30-35**). Further, we revealed the high Jaccard similarity coefficient by integrating all the replication results (Jaccard = 0.79, 95% CI: 0.75-0.96, p < .001, one-sample ratio Z test), which indicated high reproducibility of main findings in the current study.

## Discussion

Utilizing advanced functional connectome gradient techniques combined with neurotransmitomic and transcriptomic encoding (and corresponding GO, cell type-specific and chromosome-enrichment) approaches, we aimed at facilitating a more holistic and comprehensive understanding of the molecular and transcriptomic signatures of image-derived phenotypes for ADHD. We revealed connectome gradient perturbations in individuals with ADHD, with evidence for dysfunctional gradient architecture in brain systems and networks implicated in basic visual or sensory processing (i.e., MTG, VIS, SMN) and transmodal nodes involved in high-level cognitive and regulatory processes (i.e., lateral OFC, DMN). Changes in the cortical pattern of the principal gradient for ADHD patients displayed positive associations with neurotransmitoic-derived densities of GABA_a/bz_ and 5-HT_2A_, and were robustly associated with the expression profiles of gene sets related to ADHD. Positive and negative associations - PLS1+ or PLS1-, respectively-were observed for: (a) a link with connectome gradient changes, which were decoded to involve brain connections of VIS, LIB and cognitive domains of reward, inhibitory control and emotional monitoring as well as cortical metabolism of CMRO2/CMRGlu2; (b) a decoding of risky transcriptomic signatures for neurological and psychiatric diseases including ASD, Dyslexia and PTSD; (c) an enrichment for the regulation of secretion, chemotaxis and synaptic vesicle cycle, and enrichment of OPCs and Endo, as well into chromosome 18, 21 and X. Accordingly, the current study demonstrated cortical gradient perturbations in ADHD, and further provided insights into associations with gradient-derived phenotypes of ADHD with molecular, cellular and neuroimaging-transcriptomic signatures.

### Cortical gradient perturbations in ADHD

By using connectome gradient analysis which models the intrinsic functional brain architectures in a biologically plausible fashion, we identified dysregulated connectome gradient scores encompassing the LOFC and MTG in ADHD patients^9, 33–35^. Abnormal architecture of the regional connectome gradients in these regions were associated with cognitive domains such as wording and episodic memory processing. Since the MTG and LOFC play a key role in the memory and language domain, respectively ^36, 37^, our findings may reflect dysfunctions in these domains, yet results may be limited by the independent function of single brain regions or networks. Further, the observation of gradient perturbations in the DMN align with a recent overarching neurotheoretical model of ADHD proposing a dysregulated connectome-developments of DMN as key pathological candidate mechanism^38–43^.

### Neurotransmitomic markers of gradient-derived phenotype

The current image-derived diagnostic markers for ADHD have a comparably low accuracy, and the molecular underpinnings, particularly the neurotransmitomic receptors mappings may provide further insights with a high highly explainability (r^2^ > 0.70)^44, 45^. Leveraging a recent cortical neurotransmitter atlas^22^, we found molecular variants dominating ADHD-specific cortical differences in specific neurotransmitter systems, particularly GABA and serotonin receptors^22^. The decreased GABA densities across cortical areas in ADHD could result in interneuronal deficits disrupting brain structural or functional outcomes^46–48^, abnormal changes of cortical GABA receptors may shape the connectome gradient, triggering ADHD clinical symptoms. Serotonin is involved in several neurodevelopmental and regulatory functions critically associated with ADHD, and this the link between the gradient dysregulations and cortical 5-HT_2A_ receptor may reflect that: 1) several etiologic risk factors (i.e., T102T/C, 1438A>G of HRT2A) for ADHD could directly determine synthesis of 5-HT_2A_ receptor across cortical areas^49–53^, 2) variations of signaling pathways of 5-HT_2A_ receptor moderate executive functions^54^ critically impaired in ADHD, 3) variations in 5-HT signaling mediate the impact of early life stress on emotional regulation and brain architecture^55, 56^. As such, enriched distributions of 5-HT_2A_ receptor along with principle gradient changes may reflect molecular-neural dysfunctions for ADHD.

### Transcriptomic genetic vulnerability for gradient-derived changes

Despite molecular evidence on the neurotransmitter level, ADHD relevant gradient perturbations could be ascribed to systematical alterations from gene expression, molecular processing or cell pathways. The dysfunctions of gradient-derived phenotype could be moderated by transcriptomic risk signatures for neuropsychiatric disorder^20, 57^. In line with this perspective, we further determined specific gene expression patterns that co-vary with gradient changes of ADHD.

Expression of both PLS1 and PLS2 gene lists showed significant associations with aberrant connectome gradient changes in ADHD, especially in the DYDC2, OSBPL6 and CDH13. The DYDC2 was found to erroneously modify or be direct involved in neurodevelopmental processes and has been proposed as a etiological genetic risk factor for disorders characterized by emotion dysregulations such as bipolar disorder^58, 59^. It is conceivable that over-expression of DYDC3 in ADHD patients may disrupt cortical protein coding and which in turn support dysfunctional neuronal hierarchy. Moreover, GPR6 expression could directly induce ADHD-like behaviors in rodents via mediating dopamine D1 receptors and receptor-toxin interaction in prefrontal areas^60, 61^. While there is currently no direct evidence for an association between CMYA5 and ADHD, the CMYA5 has been observed as a genetic risk maker for a range of neuropsychiatric disorders exhibiting gradient perturbations^62, 63^. Moreover, under-expressions of CHD13 has been observed in ADHD as the results of abnormal subtle coding in monoamine-glutamate circuit associated with monoaminergic signaling, particularly in DA and 5-HT^64, 65^. Molecular dysfunctions coded by CDH13 expression may confer ADHD susceptibility by subtle changes in neuronal connections and synapse formation^66, 67^ that develop into macroscale gradient dysregulations. Our findings hence revealed risk gene sets correlating to macroscale brain functional changes, extending the understanding of neuroimaging-transcriptomic mechanism.

### Gene-set decoding of PLS components

To facilitate biological plausible interpretations we further decoded associations of these PLS gene lists with neurophysiologocal components. We found that the PLS1 gene sets could robustly predict connectome architecture in both visual and limbic networks, which provide a molecular explanation of evidence that ADHD patients show dysfunctions in visual attention and reward processing^35, 68–72^. This was also paralleled by the finding that such gene expressions were closely related to cognitive functions, such as inhibitory control and reward, at the core of symptomatic dysfunctions in ADHD. Intact inhibitory control and adaptive reward processing are intimately linked to (sub)cortical-cortical communication^73, 74^ and circuit level dysregulations in the underlying dysfunctionsDLPFC and basal ganglia pathways and their genetic modification have been reported in ADHD^75, 76^. In addition, we also revealed metabolic dysfunctions of ADHD in oxygen (CMRO_2_) and glucose (CMRGlu). These metabolic associations in ADHD may reflect interaction of hormonal and neurotransmitomic regulation in the serotonin and dopamine systems^77–80^.

We additionally explored disease association of gradient-derived gene-set expressions to determine relevant psychiatric and neurological risks. The PLS1 gene was found to robustly predict a range of psychiatric disorders associated with emotional dysregulations including OCD, BP and depression via effects on brain structural and functional deficits, indicating a common mediating impact on clinical outcome from cortical phenotype^81, 82^. Results may reflect that interactions between gene expressions may lead to a range of psychiatric outcomes and may explain potential comorbidity^83, 84^, instead of an ADHD-specific genetic risk factor. The identified signatures may thus reflect that ADHD may be involved in a complex transdiagnostic genetic moderation^28, 85^.

### Functional enrichment for gradient-derived gene sets

To further identify biophysiolocial processes that are underpinned by these gene sets, the neurobiological associations with the PLS1+ components were examined and revealed a dense, clustered, tangle-interacted enrichment network for GO biological process and KEGG pathways, alongside with three-component PPI architecture. The GO biological process of “regulation of secretion” has been proposed as the fundamental and crucial biological function for human beings, such as 5-HT, dopamine and brain-derived neurotrophic factor (BDNF)^86–88^. Several previous studies have demonstrated that ADHD was caused by neurotransmitomic failures including an erroneous regulation of secretion of serotonin and dopamine systems^52, 89^.

The PPI network showed a prominent tangle architecture in calcium signaling pathway. Disruption of the calcium signaling pathway was susceptible from the transcriptomic process and could cause maladaptive regulation for high-order cognitive functions (e.g., executive functions) related to ADHD symptoms^90^. This enriches knowledge for the pathological mechanism of ADHD onto cellular signaling pathways. Indeed, we provided additional gene-neuroimaging candidate to delve into the pathophysiological understructure for ADHD.

### Cellular and Chromosome-architecture for gradient-derived gene sets

We further translated cortical gene expression patterns into cell type specific maps and captured both OPCs and Endo cells. OPCs served as the supplemental cells for inflammation-injury remyelination for Oligo engaged in modulating memory, learning and cognitive processing^91^. Dysfunctional distribution of OPCs dysregulate maladaptive myelination/neuronal processes and cognitive performances as a predisposition for ADHD^92^. Endo cells were also implicated in inflammatory responses in neuropsychiatric disorders^93–95^, and may play a potential role in the pathogenesis of ADHD. We therefore offer cellular evidence for understanding ADHD pathogenesis from an immune response perspective.

Gradient-derived phenotypes in ADHD were also enriched for specific gene expression pattern of chromosome such as 18, 21 and X. This finding is consistent with prior work identifying variants of chromosome 18 (e.g., 18q11.2-12.3 and 18q21-22) in ADHD patients^96, 97^. As the reliable risk factor for the Down syndrome, the chromosome 21 was also linked with ADHD^98, 99^, and its duplication could increase risk for neurodevelopmental disorders including ADHD^100^. Furthermore, existing studies have uncovered the chromosomal X variant in ADHD, such as Xp11.4-Xp21 and Xq23^101, 102^. This could support that ADHD is the X-linked haploinsufficiency diseases and partially explained why male-female skew in the ADHD patients ^103^.

### Limitation and future direction

Findings have to be interpreted in the context of potential limitations. Gene sets in our study may not represent a specific diagnostic biomarker for the polygenic characteristics of ADHD per se. Moreover, the group-averaged analysis could not provide a powerfully genetic signature for individual prediction or precision medicine. Future studies could employ within-subject design or use more advanced precise methods to improve both group- and individual-level prediction power. We adopted gradient-derived neuroimaging phenotypes for ADHD and captured relevant cellular and chromosomal variants. However, brain functional changes may be not be accurately aligned to cellular or chromosomal activities compared with brain structural co-variance. Finally, we decoded gene sets by using online meta-analytic models which are continuously updated, yet finding can be extended based on more comprehensive future data.

## Conclusions

This study revealed the cortical gradient perturbations for ADHD, and further mapped associations of this gradient-derived phenotype with specific neurotransmitter- and transcriptional expressions. Further, we found the biophysiological underpinnings of these gradient-derived gene sets by determining enriched underlying factors including: 1) associations with specific brain intrinsic networks, cognition ontology and reward functions, as well as cortical metabolism; 2) implications in neuropsychiatric and neurological diseases; 3) precise characterization on cellular and chromosome-architecture. In sum, these findings mapping the molecular, cellular and transcriptomic mechanism of gradient-derived phenotypes in ADHD facilitate a comprehensive knowledge for ADHD pathophysiological underpinnings.

## ACKNOWLEDGEMENT

This work was supported by the PLA Key Research Foundation (CWS20J007), PLA Talent Program Foundation (2022160258) and National Key Research and Development Program of China (2022YFC2705201).

## AUTHOR CONTRIBUTION

Z.C and T.X contributed to Conceptualization, Methodology, Software, Visualization, and Writing - Original Draft; X.L, K.M, B.X and Z.G contributed to Data curation, Methodology and Validation; B.B, R.Z, Z.H, J.C and Y.T contributed to Writing - Review & Editing and Methodology; Z.X contributed to Validation, Data curation and Methodology; T.F and Z.F contributed to Project administration and Funding acquisition.

## AVAILABILITY OF DATA AND MATERIALS

Neuroimaging data can be found in ADHD-200 Sample initiative (http://fcon_1000.projects.nitrc.org/indi/adhd200/index.html); Pre-processing data for these neuroimages can be found in the Supplementary Data 1; Gene expression data that was used for transcriptional analysis can be found in the ABHA database (https://human.brainmap.org/static/download); Data and codes for neurotransmitters can be found at Hansen’s work https://github.com/netneurolab/hansen_receptors; Cell-specific gene enrichment data was extracted from Seidlitz’s dataset (https://staticcontent.springer.com/esm/art%3A10.1038%2Fs41467-020-17051-5/MediaObjects/41467_2020_17051_MOESM8_ESM.xlsx); Codes for estimating gradients and PLS components can be found at Xia’s work (https://github.com/mingruixia/MDD_ConnectomeGradient); Brain maps and plots were built by Surf Ice (https://www.nitrc.org/projects/surfice) and R packages.

