## Supplemental Information for "Cortical gradient perturbation in attention deficit hyperactivity disorder correlates with neurotransmitter-, cell type-specific and chromosome- transcriptomic signatures"

Running title: Transcriptional and molecular markers for ADHD

**Authors:** Zhiyi Chen^1,2†^**✉**, Ting Xu^3,4†^, Xuerong Liu^1†^, Benjamin Becker^3,4^, Wei Li^1^ Kuan Miao^1^, Zheng Gong^1^, Rong Zhang^2^, ZhenZhen Huo^2^, Bowen Hu^2^, Yancheng Tang^5^, Zhibin Xiao^6^, Zhengzhi Feng^1,2^**✉**, Ji Chen^7,8^, Tingyong Feng^2^**✉**

**Affiliation:**

^1^ Experimental Research Center for Medical and Psychological Science (ERC-MPS), School of Psychology, Third Military Medical University, Chongqing, China

^2^ Faculty of Psychology, Southwest University, Chongqing, China

^3^ The Center of Psychosomatic Medicine, Sichuan Provincial Center for Mental Health, Sichuan Provincial People's Hospital, Chengdu, China

^4^ The Clinical Hospital of Chengdu Brain Science Institute, MOE Key Laboratory for Neuroinformation, University of Electronic Science and Technology of China, Chengdu, China

^5^ School of Business and Management, Shanghai International Studies University, Shanghai, China

^6^ State Key Laboratory of Cognitive Neuroscience and Learning, Beijing Normal University, Beijing, China

^7^ Department of Psychology and Behavioral Sciences, Zhejiang University, Hangzhou, China

^8^ Department of Psychiatry, The Fourth Affiliated Hospital, Zhejiang University School of Medicine, Yiwu, Zhejiang, China

**✉** Corresponding at:

Zhiyi Chen) or Zhengzhi Feng), Experimental Research Center for Medical and Psychological Science (ERC-MPS), School of Psychology, Third Military Medical University, Chongqing, 400038 P.R. China; or Tingyong Feng), Faculty of Psychology, Southwest University, Chongqing, 400415, P.R. China.

**Summary:**

1 of 1 Supplementary Data

6 of 6 Supplementary Methods

17 of 17 Supplementary Results

13 of 13 Supplementary Color Figures

35 of 35 Supplementary Tables

**SUPPLEMENTAL METHODS**

1. **Participants**

**Diagnostic benchmarks**

The ADHD cohort obtained from ADHD-200 Sample initiative by recruiting data in eight image sites. To control symptom heterogeneity, we included ADHD patients who are combined type for ADHD diagnosis and medicine-naive. Diagnostic criterion for combined symptoms for ADHD are slightly different across these image sites. Thus, we provided diagnostic information following ADHD-200 Sample dataset fully. More details can be found in project website as below: <http://fcon_1000.projects.nitrc.org/indi/adhd200/index.html>.

**Site 1: Bradley Hospital/Brown University (BHBU)**

Psychiatric diagnoses were based on evaluation by the same board-certified child/adolescent psychiatrist (DPD) for all participants, using the Child Schedule for Affective Disorders Present and Lifetime version (KSADS-PL) administered to parents and children separately, and the Conners’ Parent Rating Scale-Revised, Long version (CPRS-LV). All participants completed the Wechsler Abbreviated Scale of Intelligence (WASI) as an overall measure of cognitive ability. Children in the ADHD group had to meet Diagnostic and Statistical Manual 4th Edition Text Revision (DSM-IV-TR) criteria for ADHD, as determined by parent and child answers to the KSADS-PL and were required to have ongoing psychiatric treatment. Exclusion criteria were comorbid mood or anxiety disorders, autistic or Asperger’s disorder, medical illness that was unstable or could cause psychiatric symptoms, or substance abuse within 2 months of participation. All but 1 ADHD participant taking psychostimulant medications (i.e., derivatives of methylphenidate or dextroamphetamine) were scanned when medication-free for at least 4 days drug half-lives. TDC participant inclusion criteria were a negative history of psychiatric illness in the participant and their first-degree relatives. Exclusion criteria were pregnancy, ongoing medical or neurological illness or past/present psychiatric or substance disorder. The All participants were aged between 7 and 17 years and had an IQ greater than 70.

**Site 2: Kennedy Krieger Institute (KKI)**

Psychiatric diagnoses were based on evaluations with the Diagnostic Interview for Children and Adolescents, Fourth Edition (DICA-IV, 1997), a structured parent interview based on DSM-IV criteria; the Conners’ Parent Rating Scale-Revised, Long Form (CPRS-R), and the DuPaul ADHD Rating Scale-IV(Reid, 1998). Intelligence was evaluated with the Wechsler Intelligence Scale for Children-Fourth Edition (WISC-IV) and academic achievement was assessed with the Wechsler Individual Achievement Test-II [Wechsler, 2002].

All study participants were between 8.0 and 11.0 years, and had a Full Scale IQ of 80 or higher. They had no history of language disorder or a Reading Disability (RD) either screened out before a visit or based on school assessment completed within 1 year of participation. RD was based on a statistically significant discrepancy between a child’s FSIQ score and his/her Word Reading subtest score from the Wechsler Individual Achievement Test-II [Wechsler, 2002], or a standard score below 85 on the Word Reading subtest, regardless of IQ score. Participants with visual or hearing impairment, or history of other neurological or psychiatric disorder were excluded.

Children assigned to the ADHD group met criteria for ADHD on the DICA-IV and either had a T-score of 65 or greater on the CPRS-R Long Form (DSM-IV Inattentive) and/or M (DSM-IV Hyperactive/Impulsive) or met criteria on the DuPaul ADHD Rating Scale IV (six out of nine items scored 2 or 3 from Inattention items and/or six out of nine scored 2 or 3 from the Hyperactivity/Impulsivity items). Children with DSM-IV diagnoses other than Oppositional Defiant Disorder or Specific Phobias were excluded. DSM-IV criteria and the aforementioned rating scales were also used to evaluate the three ADHD subtypes (Inattentive: ADHD-I; Hyperactive/Impulsive: ADHD-HI; Combined: ADHD-C). Children with ADHD were assigned to the ADHD-I group if they met criteria for inattentiveness but not hyperactivity/impulsivity on the DICA-IV, and had a T-score of 65 or greater on the CPRS Scale L, and a T-score of 60 or less on the CPRS Scale, or had a rating of 2 or 3 on six out of nine Inattention items on the ADHD Rating Scale IV and a rating of 2 or 3 on four or fewer items on the Hyperactivity/Impulsivity scale. Children were assigned to the ADHD-HI if they met criteria for hyperactivity/impulsivity but not inattention on the DICA-IV, and a T-score of 65 or greater on the CPRS Scale M and a T-score of 60 or less on the CPRS Scale L, or had a rating of 2 or 3 on six out of nine Hyperactivity/Impulsivity items on the ADHD Rating Scale IV and a rating of 2 or 3 on four or fewer items on the Inattention scale. All other children who met criteria for ADHD were assigned to the ADHD-C (Combined subtype) group. Children with ADHD taking psychoactive medications other than stimulants were excluded. Children who were taking stimulant medication were removed from these medications the day before and the day of testing.

TDC participants were required to have T-scores of 60 or below on the DSM-IV Inattention (L) and DSM-IV Hyperactivity (M) subscales of CPRS-R and no history of behavioral, emotional, or serious medical problems. Additionally, TDC individuals were not included if there was a history of school-based intervention services as established by parent interview, or if they met DSM-IV psychiatric disorder except specific phobia as reported on the DICA-IV.

**Site 3 NeuroIMAGE (NI)**

Psychiatric diagnoses were based on evaluations with the Schedule of Affective Disorders and Schizophrenia for Children—Present and Lifetime Version (KSADS-PL) administered to parents and children and the Conners’ Parent Rating Scale-Revised, Long version (CPRS-LV). Intelligence was evaluated with the Wechsler Abbreviated Scale of Intelligence (WASI). Inclusion in the ADHD group required a diagnosis of ADHD based on parent and child responses to the KSADS-PL. Psychostimulant drugs were withheld at least 24 hours before scanning, and other psychotropic medications were withheld at least 3 days before scanning. Inclusion criteria for TDC required absence of any Axis-I psychiatric diagnoses per parent and child KSADS-PL interview, as well as T-scores below 60 for all the CPRS-R: LV ADHD summary scales. Estimates of FSIQ above 80, and absence of other chronic medical conditions were required for all children.

**Site 4 New York University Child Study Center (NYU)**

Psychiatric diagnoses were based on evaluations with the Schedule of Affective Disorders and Schizophrenia for Children—Present and Lifetime Version (KSADS-PL) administered to parents and children and the Conners’ Parent Rating Scale-Revised, Long version (CPRS-LV). Intelligence was evaluated with the Wechsler Abbreviated Scale of Intelligence (WASI). Inclusion in the ADHD group required a diagnosis of ADHD based on parent and child responses to the KSADS-PL as well as on a T-score greater than or equal to 65 on at least one ADHD related index of the CPRS-R: LV. Psychostimulant drugs were withheld at least 24 hours before scanning. Inclusion criteria for TDC required absence of any Axis-I psychiatric diagnoses per parent and child KSADS-PL interview, as well as T-scores below 60 for all the CPRS-R: LV ADHD summary scales. Estimates of FSIQ above 80, right-handedness and absence of other chronic medical conditions were required for all children.

**Site 5 Oregon Health & Science University (OHSO)**

Psychiatric diagnoses were based on evaluations with the Kiddie Schedule for Affective Disorders and Schizophrenia (KSADS-I) administered to a parent; parent and teacher Connors’ Rating Scale-3rd Edition; and a clinical review by a child psychiatrist and neuropsychologist who had to agree on the diagnosis. Intelligence was evaluated with a three-subtest short form (Block Design, Vocabulary, and Information) of the Wechsler Intelligence Scale for Children, Fourth Edition.

Children were excluded if they did not meet criteria for ADHD or non-ADHD groups (i.e. children deemed sub-threshold by the clinicians were excluded). Children were also excluded if a history of neurological illness, chronic medical problems, sensorimotor handicap, autistic disorder, mental retardation, or significant head trauma (with loss of consciousness) was identified by parent report, or if they had evidence of psychotic disorder or bipolar disorder on the structured parent psychiatric interview. Children prescribed short-acting stimulant medications were scanned after a minimum washout of five half-lives (i.e., 24-48 hours depending on the preparation).

Typically developing control children (TDC) were excluded for presence of conduct disorder, major depressive disorder, or history of psychotic disorder, as well as for presence of ADHD.

**Site 6 Peking University (Peking)**

Study participants with the diagnosis of ADHD were initially identified using the Computerized Diagnostic Interview Schedule IV (C-DIS-IV). Upon referral for participation to the study participation, all participants (ADHD and TDC) were evaluated with the Schedule of Affective Disorders and Schizophrenia for Children—Present and Lifetime Version (KSADS-PL) with one parent for the establishment of the diagnosis for study inclusion. The ADHD Rating Scale (ADHD-RS) IV was employed to provide dimensional measures of ADHD symptoms. Additional inclusion criteria included: (i) right-handedness, (ii) no lifetime history of head trauma with loss of consciousness, (iii) no history of neurological disease and no diagnosis of either schizophrenia, affective disorder, pervasive development disorder, or substance abuse and (iv) full scale Wechsler Intelligence Scale for Chinese Children-Revised (WISCC-R) score of greater than 80. Psychostimulant medications were withheld at least 48 hours prior to scanning. All research was approved by the Research Ethics Review Board of Institute of Mental Health, Peking University. Informed consent was also obtained from the parent of each subject and all of the children agreed to participate in the study.

**Site 7 University of Pittsburgh (UP)**

Participants were drawn from a community sample and denied any significant history of psychiatric, neurological or medical illness. The abbreviated WASI was employed to assess IQ. Medication status provided is based on time of scanning, not lifetime history.

**Site 8 Washington University School of Medicine/St.Louis Children Hospital (WUSM)**

Participants were drawn from a community sample and denied any significant history of psychiatric, neurological or medical illness. The abbreviated WASI was employed to assess IQ.

**Propensity score matching (PSM)**

Given the unbalance case-control ratio, we estimated the matching propensity scores for each participant by taking all the covariate of no interests into accounts to between-group heterogeneity, including image site, gender, age, handedness verbal intelligence and intelligent performance. Specifically, the logistic function was used to fitting propensity scores by these covariates. Then, we used nearest neighbor matching with caliper to mark “match” cases. Further, the average treatment effect for the treated group (ATT) was estimated as statistics. Finally, the matched labels in TD groups have been outputted if they are balanced. By estimating propensity scores, 93 TD individuals were marked as matching cases. Standardized mean differential variances (MDV) between ADHD and matched TD group was found to be around 0.00, whilst variance ratio (VR) was found to be around 1.0. Both indices reflected balanced covariance for matched groups (see Figure S1-2).

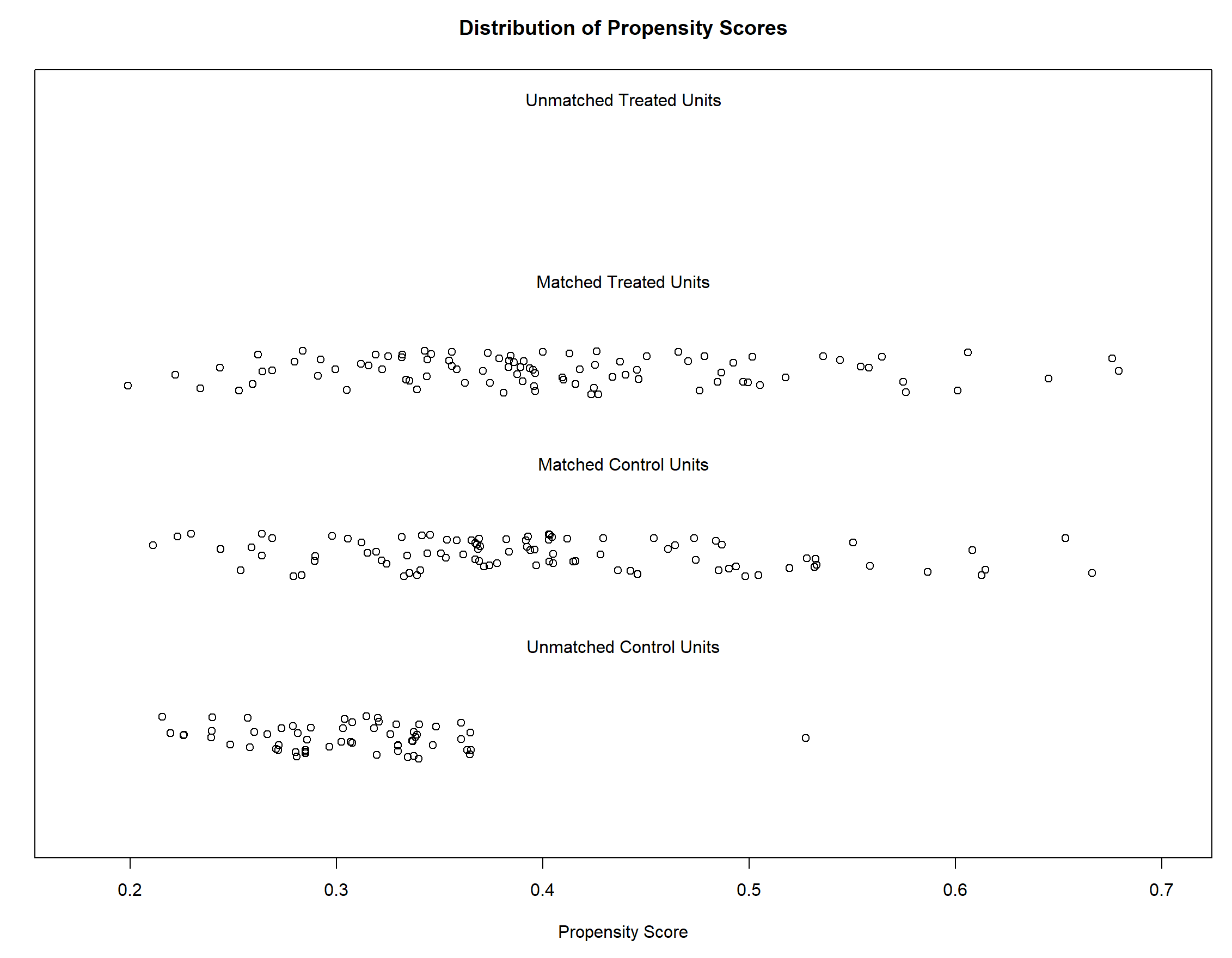

**Figure S1** Propensity density plot for both groups (treated units for ADHD patients and control units for TD individuals).

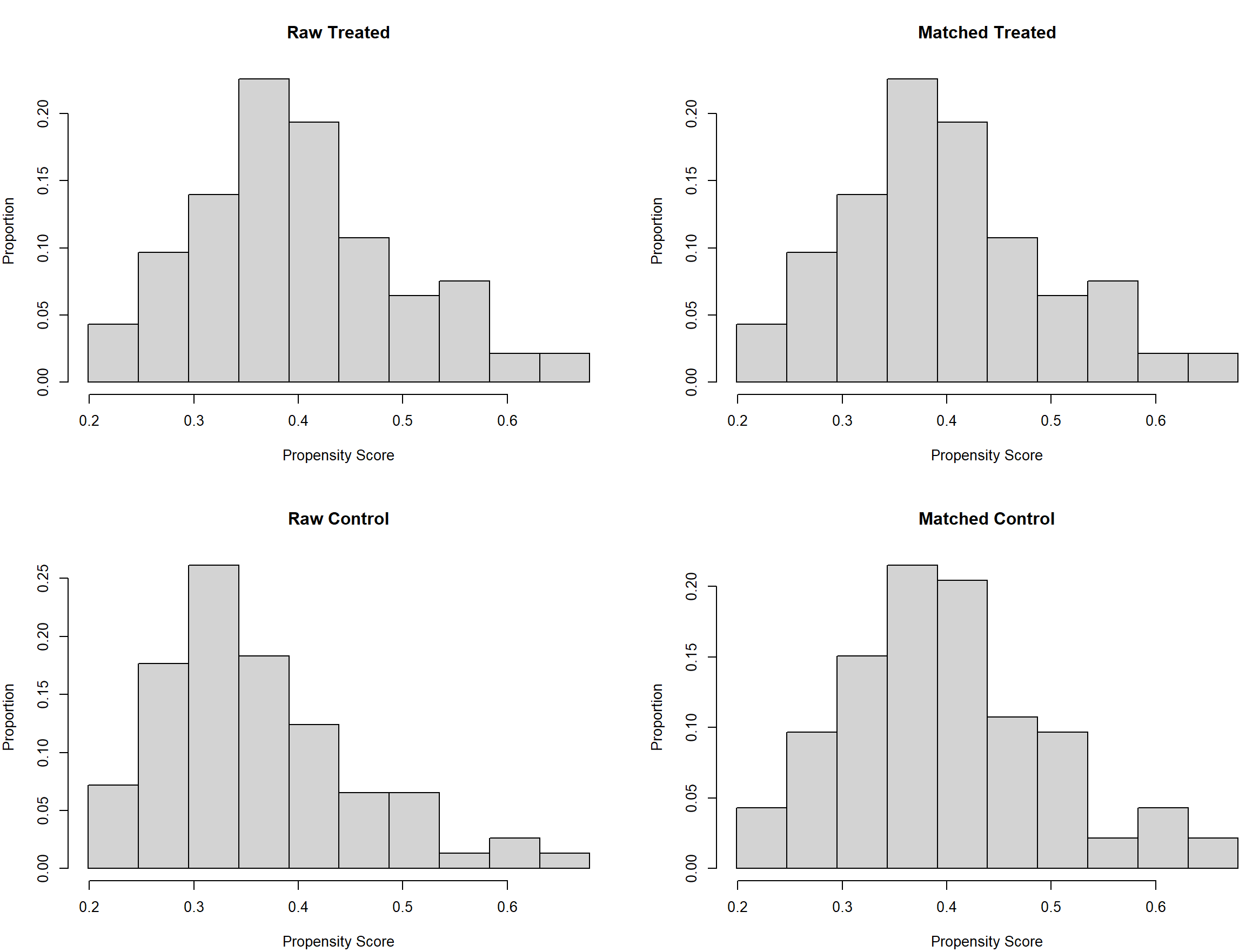
**Figure S2** Comparison for frequency of both groups before and after PSM (treated units for ADHD patients and control units for TD individuals).

2. **Preprocessing and quality control**

Image preprocessing was fully followed by NeuroBureau NIAKPipeline. Preprocessing pipeline can be found at NITRC Neurobureau website (https://www.nitrc.org/frs/?group_id=383). It should be mentioned that the noise detection was performed by CORSICA correction method, rather conventional regression models. Here, we cited report of Neurobureau NIAKPipeline to detail main steps for the NIAK pipelines that were used in the current study:

**T2 functional images:**

(1) Slice timing correction (piecewise cubic spline temporal interpolation);
(2) Motion correction (rigid-body, to the median volume of the resting-state of each run, and then between-runs/sessions);
(3) Quality control for motion correction;
(4) Linear and non-linear spatial normalization of the anatomical image (and many more anatomical stuff such as brain masking and CSF/GM/WM classification, see below);
(5) Coregistration of the anatomical volume with the mean functional volume;
(6) Concatenation of the T2-to-T1 and T1-to-stereotaxic-nl transformations;
(7) Extraction of mean/std/mask for functional images, in various spaces (Linear and non-linear stereotaxic spaces);
(8) Quality control for 4 and 5;
(9) Correction of slow time drifts (high-pass filtering with discrete cosines, cut-off frequency 0.01Hz);
(10) Correction of physiological noise (20 components, selection threshold 0.15);
(11) Resampling of the functional data in the stereotaxic space (tricubic spatial interpolation, 3mm isotropic resolution);
(12) Spatial smoothing (Gaussian kernel, 6mm isotropic).

**T1 anatomical images:**

(1) Non-uniformity correction in native space (without mask);
(2) Brain extraction in native space;
(3) Linear coregistration in stereotaxic space (with mask from 2);
(4) Non-uniformity correction based on the template mask;
(5) Brain extraction, combined with the template mask;
(6) Intensity normalization;
(7) Non-linear coregistration in template space (with mask from 5);
(8) Generation of the brain mask in the non-linear stereotaxic space by intersection of the template mask with a head mask;

Generation of the mask in the stereotaxic linear space by application of the inverse non-linear transform from 7 and the brain mask from 8.

**Quality control**

**T1-fMRI Co-registration:**

KKI: The T1-fMRI failed for four subjects (X_1577042, X_2344857, X_8628223, X_3699991). The pipeline was restarted on these subjects only, after selecting an initial realignement of the center of mass of the brain masks in T1 and fMRI (opt.anat2func.init = 'center'). This solved the problem. The results of these four subjects have been updated for the release, except for the QC results which correspond to the first pass;

OHUS: The T1-fMRI coregistration failed for four subjects (X_3358877, X_3812101, X_4072305, X_4529116). The pipeline was restarted on these subjects only, after selecting an initial realignement of the center of mass of the brain masks in T1 and fMRI (opt.anat2func.init = 'center'). The new registration was satisfactory and the results of the four subjects have been updated for the release;

Peking: Three subjects (X_2919220, X_2268253, X_2950754) were missing dorsal slices and should be excluded from analysis. This was found manually on a subset of subjects and more might have the same problem. The T1-fMRI coregistration failed for two subjects (X_1805037, X_4053836). The pipeline was restarted on these two subjects only, after selecting an initial realignement of the center of mass of the brain masks in T1 and fMRI (opt.anat2func.init = 'center'). The new registration was satisfactory and the results of the two subjects have been updated for the release;

UP: The T1-fMRI registration worked well for all subjects. Subject X_0016029 was found to be missing some dorsal slices in the fMRI data and other subjects in the dataset might also share this problem;

NYU: The T1-fMRI coregistration initially failed for thirty-six subjects. The pipeline was restarted on these subjects only, after selecting an initial realignement of the center of mass of the brain masks in T1 and fMRI (opt.anat2func.init = 'center'). This worked for all subjects and the results have been updated for the release (QC results correspond to the first pass);

NeuroIMAGE: The T1-fMRI coregistration initially failed for a majority of subjects and the pipeline was restarted with an initial realignement of the center of mass of the brain masks in T1 and fMRI (opt.anat2func.init = 'center'). This worked for all subjects and the results and QC have been updated for the release;

WU: The T1-fMRI coregistration initially failed for three subjects and the pipeline was restarted with an initial realignement of the center of mass of the brain masks in T1 and fMRI (opt.anat2func.init = 'center'). This worked for all subjects and the results and QC have been updated for the release;

**T1-stereotaxic space coregistration:**

KKI: QC only one subject with a problem in the non-linear coregistration in stereotaxic space: X_2740232. This subject has a very asymmetric head, and the non-linear coregistration algorithm has pushed part of the skull into the occipital cortex. Because the coregistration on the rest of the brain is satisfactory, this subject may still be included in an analysis;

OHSU: Six subjects showed sub-standard quality of the T1 scans that resulted in poor normalization to stereotactic space (X_2292940, X2535204, X_2920716, X_1743472. X_1536593, X_3560456). This is likely attributable to excessive movement and we advise excluding these subjects from the analyses;

Peking: One subject (X_3291029) showed an acquisition artifact and should be excluded from analysis;

UP: The non-linear coregistration in stereotaxic space of the T1 images worked well for all subjects;

NYU: Twelve subjects showed poor normalization to stereotactic space. The pipeline was restarted with 'Nu correct distance = 100' for these subjects. This corrected the problems for five subjects and the remaining seven should be excluded from analyses (X_0010013, X_2297413, X_3679455, X_0010032, X_1435954, X_6206397, X_8415034);

NeuroIMAGE: The normalization to stereotactic space initially failed for a number of subjects and was restarted with the option 'Nu correct distance = 100'. This corrected most problems and the coregistration failed for only one subjects (3304956) which should be excluded from the analyses;

WU: The non-linear coregistration in stereotaxic space of the T1 images worked well for all subjects;

**Head-motion quality**

KKI: Only one subject has signs of severe motion (X_3103809) with over 3 degrees in translation of rotation parameters. Otherwise the following subjects had a slightly high amount of motion (over 1 mm transtition in translation, or 1 degree in transition in rotation) : X_2703289, X_1541812, X_2299519, X_1846346, X_2138826, X_1577042, X_1686265, X_1962503, X_2360428, X_8432725, X_3310328 Those can still be included in an analysis. The remaining subject had negligible motion (less than 1 mm transtition in translation, or 1 degree in transition in rotation);

OHSU: Of the 79 subjects, 44 showed minimal movement. 19 subjects had moderate movement of 1 to 3mm in translation and 1-3 degrees in rotation and can still be considered for inclusion in the analyses (X_4529116, X_3812101, X_3652932, X_2426523, X_2054310, X_2124248, X_8064456, X_3466651, X_4072305, X_2790141, X_4219416, X_3869075, X_2929195, X_3470141, X_4103874, X_2561174, X_7333005, X_2409220, X_3206978). 16 subjects showed movement exceeding 3mm translation or 3 degrees rotation and should be excluded from analysis (X_3560456, X_2455205, X_1386056, X_2288903, X_3286474, X_1696588, X_2620872, X_3684229, X_8720244, X_2920716, X_2559559, X_2054998, X_1536593, X_2845989, X_2292940, X_2535204);

Peking: Of the 194 subjects, 175 showed minimal movement. 14 subjects had moderate movement of 1 to 3mm in translation and 1-3 degrees in rotation and can still be considered for inclusion in the analyses (X_2529026, X_3205761, X_4334113, X_2107404, X_2174595, X_2228148, X_1805037, X_1494102, X_9002207, X_3593327, X_3561920, X_4073815, X_1947991, X_3993793, ). 5 subjects showed movement exceeding 3mm  translation or 3 degrees rotation and should be excluded from analysis (X_1791543, X_4225073, X_2296326, X_1860323, X_2367157);

UP: Of the 89 subjects 65 showed minimal movement. 14 had movement ranging from 1 to 3 mm (X_0016069, X_0016009, X_0016028, X_0016005, X_0016083, X_0016034, X_0016048, X_0016044, X_0016013, X_0016054, X_0016041, X_0016047, X_0016053, X_0016003). 10 subjects had movement exceeding 3mm and should excluded from analysis (X_0016072, X_0016040, X_0016024, X_0016023, X_0016025, X_0016015, X_0016026, X_0016017, X_0016007, X_0016079);

NYU: Of the 216 subjects 136 showed minimal movement. 55 had movement ranging from 1 to 3 mm (X_0010032, X_4562206, X_0010074, X_0010086, X_4079254, X_3542588, X_0010012, X_1992284, X_9578663, X_1700637, X_2741068, X_2030383, X_3433846, X_2107638, X_4084645, X_0010018, X_1780174, X_0010031, X_0010011, X_0010102, X_3619797, X_0010060, X_0010050, X_1497055, X_3235580, X_3243657, X_0010041, X_1000804, X_1854959, X_2230510, X_1918630, X_0010090, X_0010044, X_0010028, X_2054438, X_0010049, X_3845761, X_0010096, X_0010047, X_0010019, X_2136051, X_2821683, X_0010022, X_0010033, X_3163200, X_0010062, X_2735617, X_0010092, X_2907383, X_3349205, X_2950672, X_5971050, X_0010106, X_3653737, X_0010085).25 subjects had movement exceeding 3mm and should excluded from analysis (X_8915162, X_0010061, X_0010025, X_0010117, X_6568351, X_3518345, X_3601861, X_4187857, X_0010015, X_0010014, X_0010078, X_0010030, X_1187766, X_0010046, X_0010095, X_0010077, X_0010005, X_3662296, X_0010119, X_0010103, X_0010003, X_0010066, X_0010111, X_0010067, X_1023964);

NeuoIMAGE: Of the 48 subjects 37 showed minimal movement. 6 had movement ranging from 1 to 3 mm (1580708, 1312097, 3108222, 4239636, 1438162, 5045355). Five subjects had movement exceeding 3mm and should excluded from analysis (2029723, 3048588, 3082137, 4919979, 3808273);

WU: Of the 49 subjects 23 showed minimal movement. 17 had movement ranging from 1 to 3 mm (X_0015012, X_0015021, X_0015025, X_0015029, X_0015035, X_0015055, X_0015050, X_0015022, X_0015002, X_0015044, X_0015049, X_0015037, X_0015047, X_0015008, X_0015003, X_0015051, X_0015056). Nine subjects had movement exceeding 3mm and should excluded from analysis (X_0015034, X_0015015, X_0015020, X_0015010, X_0015060, X_0015059, X_0015038, X_0015046, X_0015023).

3. **Gradient estimate and data harmonization**

To obviate the bias of randomly selecting discrete parcellation atlas, we build up the functional connectome by using voxel-wise scheme, with each voxel for one node. Given the computational burden, we downsampled normalized images from 3-mm size into 4-mm isotropic resolution, and obtain 18,933 x 18,933 connectivity matrix. Furthermore, to mirror the main representation of whole-brain connectome, we selected top 10 % connections with most connectivity strength (i.e., Pearson correlation coefficient) from the voxel-wise connectivity matrix for each participant. To approach the organizational nature of brain, we used cosine similarity for estimating sparse connectivity pattern for these selected voxels (nodes). As the unclear biological or neural functions for negative functional connectivity, we scaled and normalized these resultant cosine similarity matrix into non-negative angle matrix^1, 2^. As one of the ideally suitable non-linear dimension reduction method, the diffusion map embedding has been used to detect principal gradient component (s), which translated high-dimensional cosine similarity matrix into low-dimensional space. Diffusion map embedding method aimed to translate connectivity pattern of each node into distances of high-dimensional space, with long distance for dissimilarity of connectivity pattern and vice versa. In this vein, high gradient scores reflected the connectivity similarity for a given node to other nodes. Given no a prior knowledge for ADHD gradient, we initially set learning ratio to be 0.5 for diffusion model, and used iterative Procrustes rotation to co-register individual gradient maps^1, 3, 4^.

We estimated three indices for quantifying the global gradient difference for connectome, including gradient explanation ratio, gradient range and gradient variation. Gradient explanation ratio calculated the extent to which a given gradient explain the variance for connectome (%), with high ratio for a robust hierarchical organization in the embedding axis. In addition, the gradient range reflected the gradient-score difference between the maximum and minimum value, which quantify the differentiation for a given connectome. Lastly, the gradient variation was calculated as the standard deviation of these gradient scores in the connectome. This index reflect how much the heterogeneity for connectivity pattern is in the connectome. Gradient estimates were implemented by BrainSpace toolbox^5^.

Given the potential multi-site heterogeneity, we used ComBat suit to correct these across-site noises. ComBat was a Bayesian multivariate linear regression algorithm to model the “true effect” for variate of interest as as a linear combination of multi-site differences and variates of no interests^6, 7^. In addition, the model error was assumed to be moderated by site-specific variances. Thus, by using ComBat algorithm, the indices for gradients have been corrected for both multi-site variants and potential confounding factors. More details for ComBat toolbox can be found elsewhere (13,14).

4. **Partial least squares (PLS) model**

Given the co-linearity structure for neurotransmitter systems, we build upon the partial least square (PLS) to fitting the 19 neurotransmitter receptors/translator distribution density to z score for regional gradient differences. In this model, the first component could be captured by a linear combination of standardized predictors and responses, which was required to extract maximum variances (u_1_, v_1_). Thus, the correlation between u_1_ and v_1_ would be the maximum pair. Then, the observation matrix for each variate (***A, B***) could used to estimate the vector scores (u^^^_1_, v^^^_1_) for the first component. In this vein, the vector scores could be calculated by a Lagrange multiplier method.

$$max (u_{1}^{^}\cdot v_{1}^{^}) =\rho^{(1)T}A^{T}B^{\gamma(1)};\gamma^{(1)}=\frac{1}{\delta1}B^{T}{A\rho}^{(1)}$$

Following that, the regression model can be used for fitting response to u_1_, one-by-one and verse vice. Then, the residual matrix can be obtained as ***A1*** and ***B1****.* If the elements in both residual matrix approached zero, this component would be assumed to explained this model. If it is not this case, both residual matrix should be used to replace original correlation matrix for recalculating these statistics until the condition was fulfilled. In the resultant step, if the rank of observation matrix (n x m) was smaller than minimum value at position of (n-1, m), all the qualified components would be considered to be extracted completely. In this case, the PLS can be used to fit u~x to y~u.

5. **AHBA dataset and preprocessing**

AHBA dataset was built by AIBS (Allen Institute for Brain Science) (http://human.brain-map.org, RRID: SCR_007416) for providing whole-brain spatially heterogeneous gene expression level from six donors^8, 9^. These adult donors provided their postmortem brains (age = 42.50 ± 13.38, 1 females, two whole-brain samples and five left-hemisphere brain samples). All the donors are required to be no history of neuropsychiatric disorders or neurological conditions, such as brain injury, epilepsy and any substance abuses. These postmortem brains have been sampled within 30 hours from death (https://help.brain-map.org/download/attachments/2818165/). Each postmortem brain was firstly sliced into ~ 500 anatomical samples in each hemisphere. Then, the DNA microarray high-pass sequences technique was used to test the gene expression level for each sample. Afterwards, gene information for each sample was mapped into 3D MRI coordinate space (standard MNI space) by using precedent high-resolution T1 images. The whole dataset contained 3072 samples, and obtained qualified gene expression data from 58,692 probes. To obtain the group-averaged gene expression atlas, the normalization processes were implemented across samples and across donors. To increase regional resolution for IDP-transcriptional analysis, we initially selected HCP_1.0 MMP atlas with 360 regions for assignments. In addition, image results indicated the gradient scores in cortical areas, which suited to parcel whole-brain into this HCP atlas. Further, the connectome gradient was revealed from HCP dataset, and was thus suitable for using HCP_1.0 MMP atlas to reduce parcelliation heterogeneity. As recommended pipeline (Alchikle et al., 2019), we preprocessed whole-brain transcriptional dataset herein on 10, October, 2022 by following this workflow (https://github.com/BMHLab/AHBAprocessing). Main steps for preparing this dataset included six section. (1) Gene information re-annotation. To obviate outdated gene annotation, the probe-to-gene mapping was implemented by using re-annotator toolbox for updating annotation information (probe n = 45,821 corresponding to 20,232 genes) (https://earray.chem.agilent.com/earray) in the hg38 sequencing database; (2) To remove background noise, the IBF (intensity-based filter) was used to filter probes that exceeded background (gene n = 10,190); (3) to tackle with multiple-probes for one gene, the gene annotation was determined by the highest correlation of RNA-seq to a given gene (probe selection); (4) for IDP-transcriptional analysis, each sample tissue was aligned into HCP_MMI 1.0 atlas with 300 regions in the MNI space, and the samples would be removed once the centroid Euclidean distance of sample to region in this atlas exceeded 2 mm; by using such criterion, 820 samples were retained to cover 280 regions in the atlas, with each sample for 10,027genes; (5) to correct the inter-sample and inter-donor heterogeneity, scaled robust Sigmoid normalization has been used; for inter-sample variant, the first-step normalization was implemented across all the probes within sample; for each brain, the probes were normalized across all the sample; (6) for each region, gene expression levels are averaged across all the samples from six donors (gene selection). Following that, the atlas-gene matrix was outputted (284 x 10,027). In addition, four regions were further removed because they located outside whole-brain mask that we generated from image analysis. Thus, the final AHBA dataset that we used was the matrix with 280 x 10,027.

6. **GAMBA decoding**

GAMBA platform was used to reveal the association between gene expression level and neuroimaging-derived phenotype by regression model. Main steps for decoding the gene list that we obtained in the current study has been provided in main texts. Here provided original descriptions for statistical details^10^ (http://dutchconnectomelab.nl/GAMBA/):

The association between the cortical gene expression profile and regional properties of different neuroimaging phenotypes is assessed using linear regression.

$$y_{i} =\beta_{i} \beta1Xj+\varepsilon$$

where Y_i_ indicates the normalized gene expression profile of gene i or the averaged profile of a gene set i, X^j^ the normalized cortical profile of neuroimaging phenotype j, and cov the normalized covariate. Normalization is performed by substracting each value by the mean, followed by diving values by one standard deviation. The standardized regression coefficient β_1_ and the corresponding p-value are obtained.

Furthermore, GAMBA tests whether the observed association is spatially specific and gene specific. The null-spin, null-random, null-coexpression, and null-brain models are used. Per gene and per brain imaging phenotype, GAMBA performs a z-test to examine whether the observed β_1_ (i.e., the effect size) was higher than the average effect size observed in the null models.

$$z = (\beta_{1}-u)/\sigma$$

where µ, σ indicate the mean and standard deviation of β1 in different null conditions. A two-sided p-value was computed as follows:

$$p = 2\emptyset(-z)$$

where Φ is the standard normal cumulative distribution function. GAMBA implements Bonferroni and FDR correction with adjustable thresholds to correct for multiple comparisons in the analysis of each imaging modality. Results reaching significance are shown in darker colors, otherwise in lighter colors.

**SUPPLEMENTAL RESULTS**

1. **Connectome gradient patterns for ADHD patients**

We used diffusion map embedding method to capture connectome gradients for both groups, and predefined the maximum number of components (gradients) to be 30 as default parameters (Figure S3). It revealed that the gradient 2 explained 7.06 % (± 3.85 %) variance in connectome, with 6.68 ± 3.39 % explanation ratio (ER) for ADHD and with 7.44 ± 3.80 % ER for TD controls. However, the differences between ADHD and TD for explanation ratio did not reached statistical level (p = .17). Further, gradient 3 was reveal to explain 6.06 % (± 3.15 %) variance in connectome, with 6.29 ± 2.97 % ER for ADHD and with 5.83 ± 3.32 % ER for TD. Likewise, no significant differences between cohorts for ER as well (p = .31).

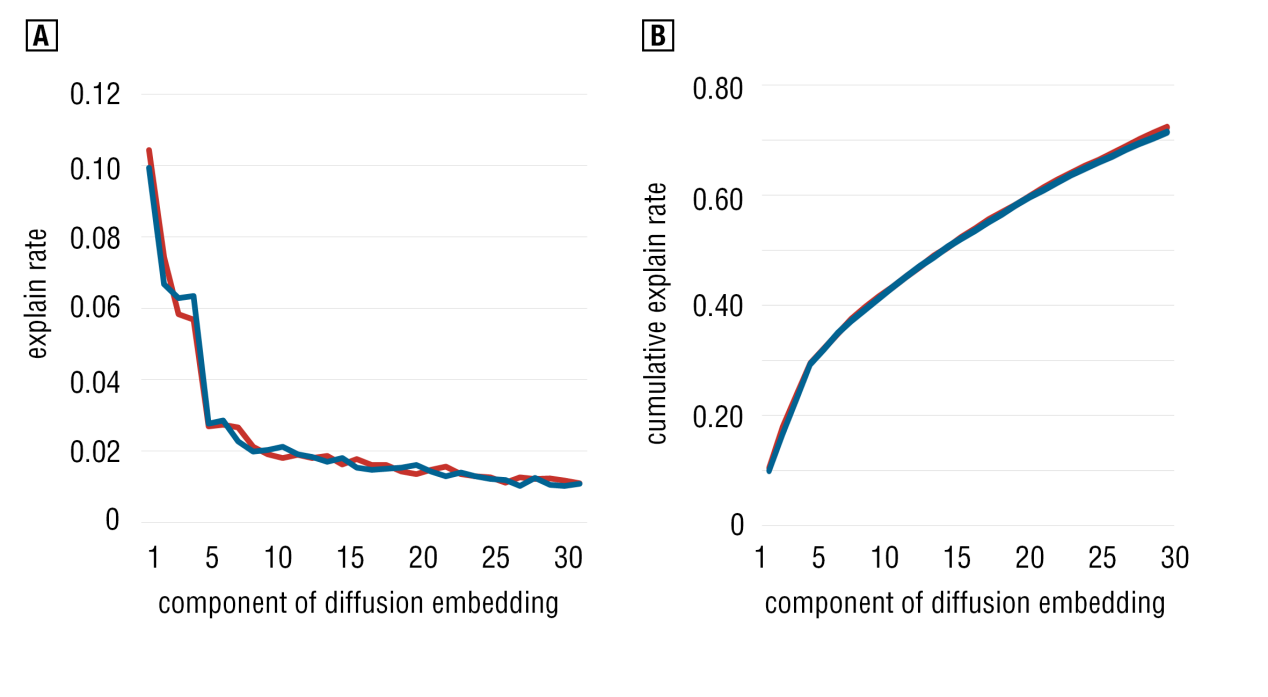
**Figure S3** Scree plot for component of diffusion embedding maps.

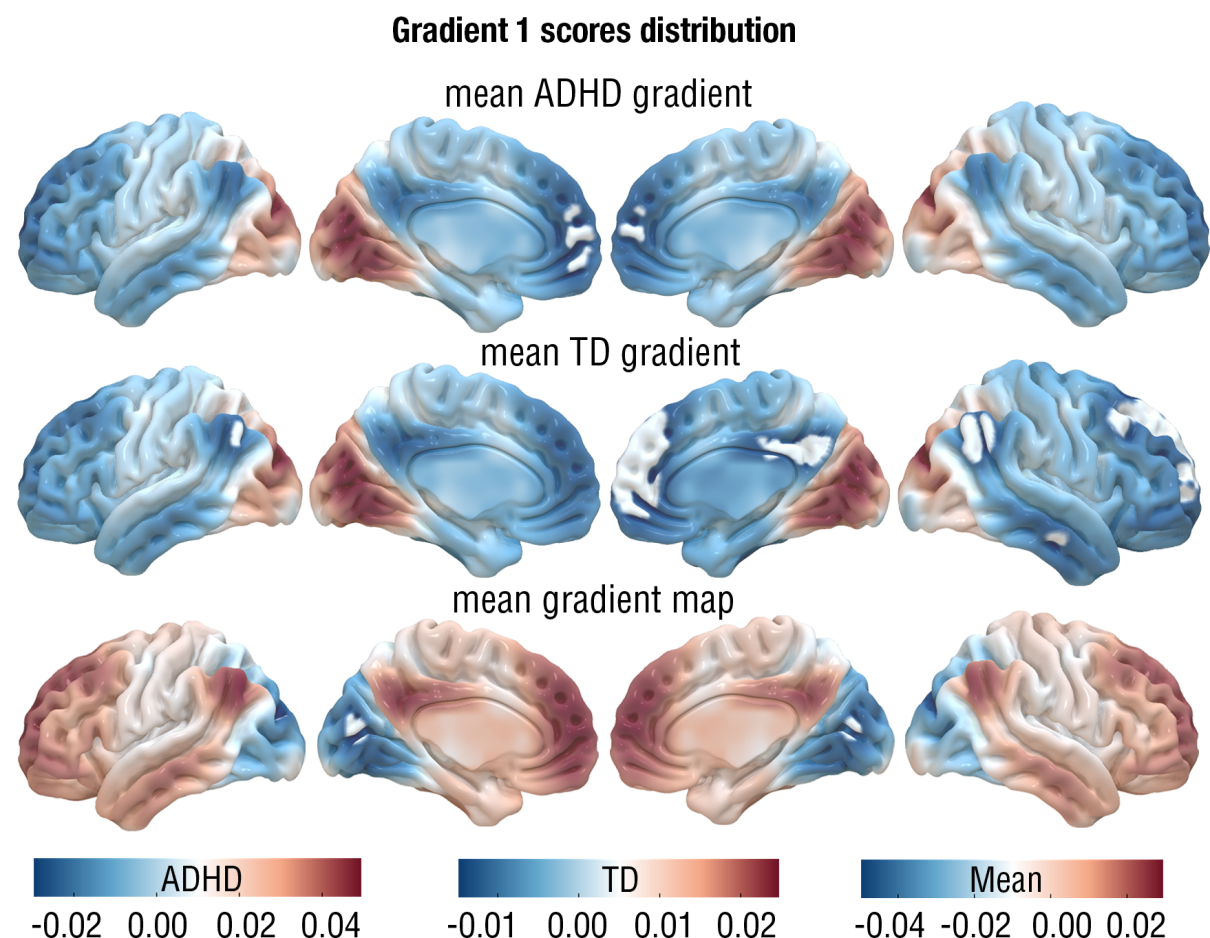

**Figure S4** Principal gradient (1) scores distribution for ADHD, TD and averaged-group one.

In addition, we still found no significant differences for other topological indices in all the gradients, including gradient range and gradient variations.

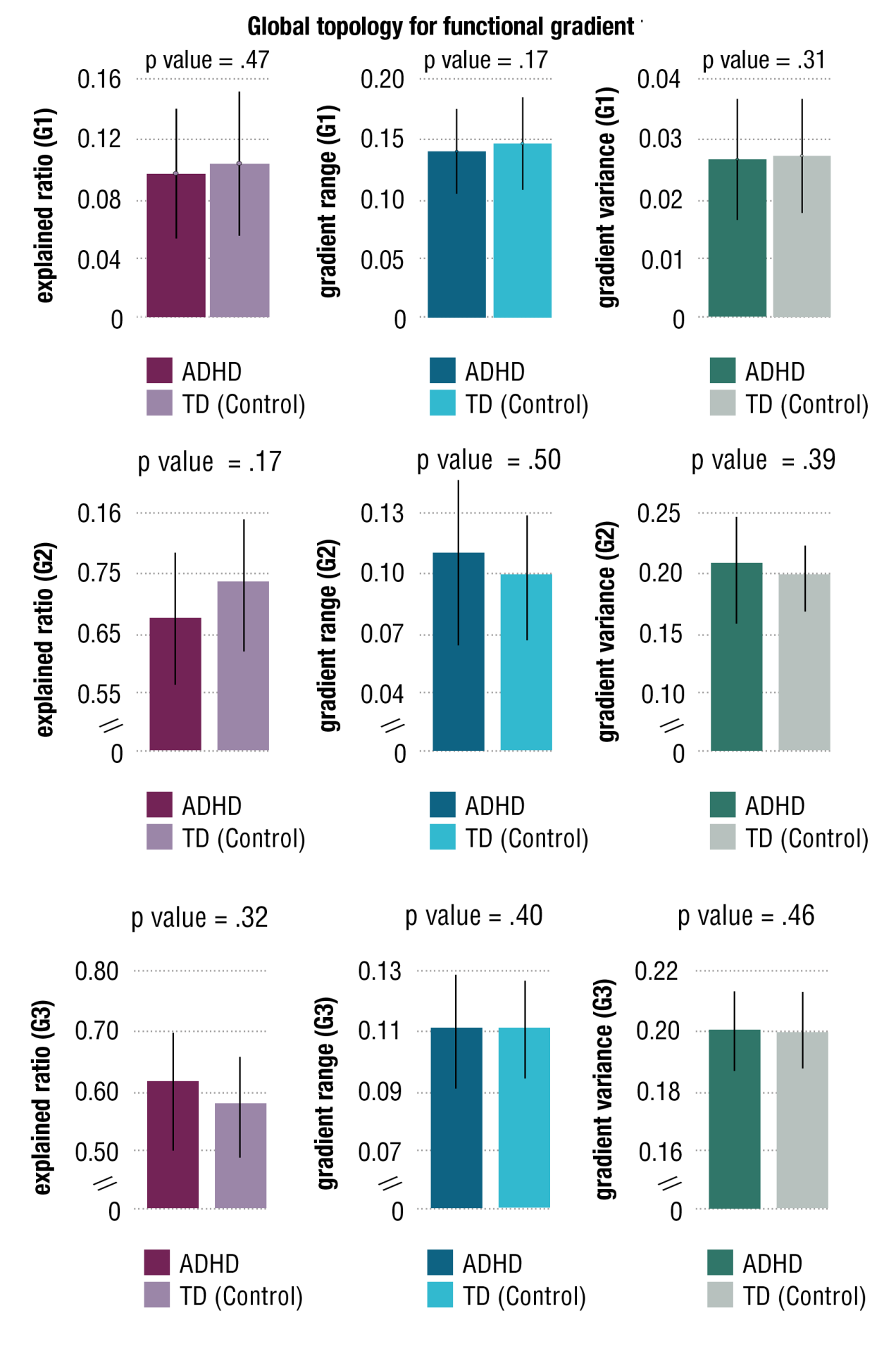

**Figure S5** Statistics for global topology of connectome gradient 1,2,3.

Furthermore, we examined the global spatial similarity between both cohorts for connectome gradients, and found the extremely high similarity for all the gradients (r = .999, p < .0001, permutation tests with spatial autocorrelation corrected).

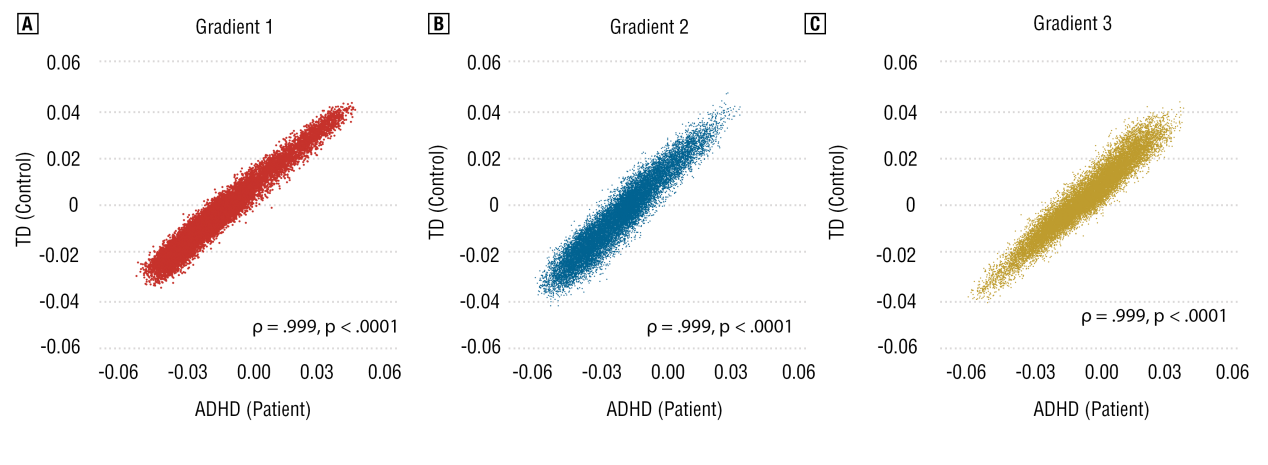
**Figure S6** Scatter plots for the spatial correlations between ADHD and TD controls for gradient scores in principal gradient and gradient 2, 3.

2. **System-based gradient changes for ADHD patients**

According to Yeo network-based parcellation, the whole-brain has been partitioned into eight brain intrinsic networks (ISN). Independent two-sample t-tests were used to examine the average-between differences for gradient scores in principal, second and third gradient. False-discovery error (FDR) correction was implemented to control multiple comparisons. Full results can be found in Table S1-3 as underneath.

|  | VIS | SMN | DAN | VAN | LIM | FPN | DMN | SUB |
| --- | --- | --- | --- | --- | --- | --- | --- | --- |
| q (FDR) | 0.0000 | 0.0000 | 0.0000 | 0.0011 | 0.0000 | 0.5097 | 0.0000 | 0.0000 |
| Cohen d | 0.2154 | 0.1654 | -0.1080 | -0.0798 | -0.4113 | 0.0167 | -0.1148 | -0.1051 |

**Table S1** System-based differences for gradient scores in principal gradient.

|  | VIS | SMN | DAN | VAN | LIM | FPN | DMN | SUB |
| --- | --- | --- | --- | --- | --- | --- | --- | --- |
| q (FDR) | 0.0000 | 0.0000 | 0.0151 | 0.0000 | 0.0000 | 0.0000 | 0.0000 | 0.0000 |
| Cohen d | -0.1563 | 0.4076 | 0.059 | 0.1453 | 0.3408 | -0.2465 | -0.1903 | -0.1577 |

**Table S2** System-based differences for gradient scores in gradient 2.

|  | VIS | SMN | DAN | VAN | LIM | FPN | DMN | SUB |
| --- | --- | --- | --- | --- | --- | --- | --- | --- |
| q (FDR) | 0.0002 | 0.0509 | 0.0000 | 0.0000 | 0.0000 | 0.0000 | 0.0000 | 0.0000 |
| Cohen d | -0.0715 | 0.0390 | -0.3390 | 0.1954 | -0.1228 | -0.1056 | 0.1541 | 0.8262 |

**Table S3** System-based differences for gradient scores in gradient 2.

3. **Cognitive decoding at NeuroSynth**

We used the “decoder” section of NeuroSynth platform to decode automatically cognitive functions that relating to the principal gradient changes of ADHD by online meta-analysis. To do so, the vertex-wise statistical map for the contrast of ADHD to TD controls in principal gradient was transferred into z-value map. Then, this Fisher z-map was provided for “decoder” section of NeuroSynth platform to obtain relevant terms correlating with spatial z values. Finally, we removed non-cognitive terms to generate the list detailing top 50 cognitive terms. Results can be found in Figure S7 as underneath.

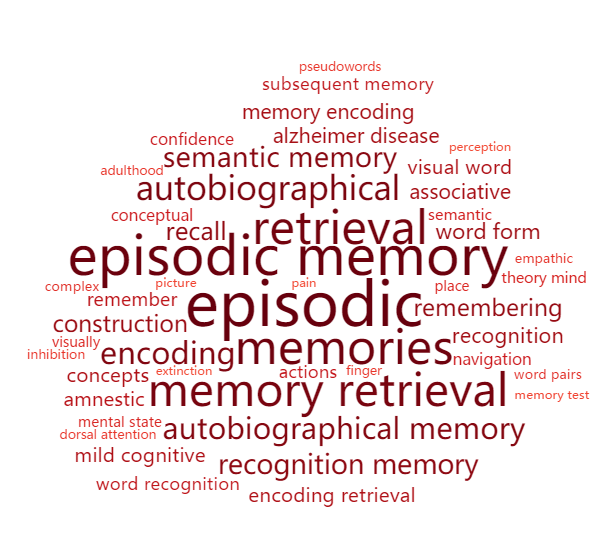

0.15

0.07

0.02

**Figure S7** Word-Cloud plot for the cognitive terms of principal gradient differences for ADHD patients. The size of font reflects corresponding correlation coefficients.

4. **Neurotransmitomic atlas**

Hansen and colleagues (2022) have collated positron emission tomography data from ~ 1,200 healthy participants to build a cortical normative atlas of 19 receptors and transporters across 9 different neurotransmitter systems. More details for these receptors and transporters have been provided into Table S4. Full information for how to gather these data for building neurotransmitter atlas can be found for original work^11^. All the receptor and transporters have been categorized into two types - that is - excitatory or inhibitory system. All the receptor and transporters were estimated by PET scanner, the tracer and measures have been detailed in following table as well.

| **Receptor/transporter** | **Neurotransmitter** | **Tracer** | **Measure** | **Type** |
| --- | --- | --- | --- | --- |
| D_1_ | Dopamine | [^11^C]SCH23390 | BP_ND_ | Excitatory |
| D_2_ | Dopamine | [^11^C]FLB-457 | BP_ND_ | Inhibitory |
| DAT* | Dopamine | [^123^I]-FP-CIT | SUVR | Transporter |
| NET* | Norepinephrine | [^11^C]MRB | BP_ND_ | Transporter |
| 5-HT_1A_ | Serotonin | [^11^C]WAY-100635 | BP_ND_ | Inhibitory |
| 5-HT_1B_ | Serotonin | [^11^C]P943 | BP_ND_ | Inhibitory |
| 5-HT_2A_ | Serotonin | [^11^C]Cimbi-36 | B_max_ | Excitatory |
| 5-HT_4_ | Serotonin | [^11^C]SB207145 | B_max_ | Excitatory |
| 5-HT_6_ | Serotonin | [^11^C]GSK215083 | BP_ND_ | Excitatory |
| 5-HTT* | Serotonin | [^11^C]DASB | B_max_ | Transporter |
| α_4_β_2_ | Acetylcholine | [^18^F]Flubatine | V_T_ | Excitatory |
| M1 | Acetylcholine | [^11^C]LSN3172176 | BP_ND_ | Excitatory |
| VAChT* | Acetylcholine | [^18^F]FEOBV | SUVR | Transporter |
| NMDA | Glutamate | [^18^F]GE-179 | V_T_ | Excitatory |
| mGluR_5_ | Glutamate | [^11^C]ABP688 | BP_ND_ | Excitatory |
| GABA_A/BZ_ | GABA | [^11^C]Flumazenil | B_max_ | Inhibitory |
| H3 | Histamine | [^11^C]GSK189254 | V_T_ | Inhibitory |
| CB1 | Cannabinoid | [^11^C]OMAR | V_T_ | Inhibitory |
| MOR | Opioid | [^11^C]Carfentanil | BP_ND_ | Inhibitory |

**Table S4** Neurotransmitter receptors and transporters included in analyses. BP_ND_, non-displaceable binding potential; VT, tracer distribution volume; B_max_, density (pmol ml^-1^) converted from binding potential (5-HT) or distributional volume (GABA) using autoradiography-derived densities; SUVR, standard uptake value ratio. * means transporters. Types indicated the receptor functions.

5. **Neurotransmitomic signatures of gradient-derived phenotype in ADHD**

To reveal neurotransmitomic association for the gradient-derived phenotype of ADHD in second and third gradient, we capitalized on partial regression model (PLS) to fit the neurotransmitomic matrix (i.e., predictors, regionally cortical neurotransmitomic distribution) to case-control z-value vector (i.e., response, regionally cortical differences for second gradient). PLS1 was found to explain most 25.33% variance in this model (p < .01), whilst PLS2 was found to explain 23.82%. variance (p < .01) here. Further, we estimated the weight z value for each receptor/transporter in PLS1 and PLS2 loading by using Bootstrapping method, respectively. These cortical neurotransmitomic weights were then mapped into Schaefer-100 atlas for PLS1 and PLS2. PLS1 and PLS2 neurotransmitomic weights map was presented in the Figure S8. Supporting that, we observed a statistically significant correlation between the PLS1 weights and case-control z values for second gradient (r = 31, p < .001, spatially-corrected). In addition, the correlation between PLS2 scores and case-control z values was found to reach statistical significance without correction (r = .21, p < .03, uncorrected).

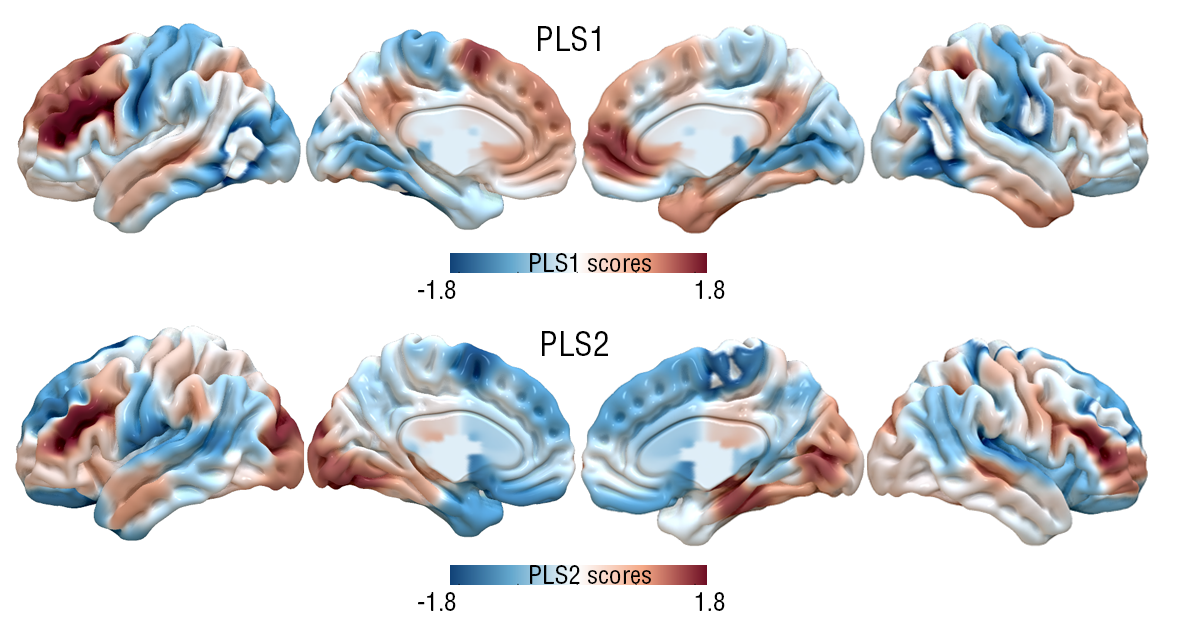
**Figure S8** PLS1 and PLS2 scores maps for PLS models in gradient 2.

For probing into the neurotransmitomic signatures of ADHD-specific gradient perturbation in the gradient 2, we re-ranked weights (z values) of each receptors and transporters with descending order firstly. Then, the p value for each z value was estimated from sampling Z distribution. However, no survived receptors and transporters for Bonferroni-Hochberg correction at p < .001 in the PLS1 and PLS2 component (Z > 4.2 or Z < -4.2) (Table S5-6).

| **Receptor/Transporter** | **ID** | **Z scores** | **P values** | **Type** |
| --- | --- | --- | --- | --- |
| D1 | 1 | 3.088109 | 0.002014346 | Receptor |
| NET | 4 | 2.389311 | 0.016880007 | Transporter |
| GABABZ | 16 | 2.348229 | 0.01886292 | Receptor |
| DAT | 3 | 2.191292 | 0.028430669 | Transporter |
| 5-HT4 | 8 | 2.036602 | 0.041689942 | Receptor |
| 5-HTT | 10 | 1.672973 | 0.094332611 | Transporter |
| 5-HT6 | 9 | 1.59175 | 0.111440889 | Receptor |
| alpha4-beta2 | 11 | 1.423851 | 0.154489609 | Receptor |
| 5-HT1A | 5 | 0.830551 | 0.406227326 | Receptor |
| 5-HT1B | 6 | 0.472397 | 0.636643445 | Receptor |
| mGluR5 | 15 | 0.153857 | 0.877722489 | Receptor |
| MI | 12 | 0.139747 | 0.88885989 | Receptor |
| CB1 | 18 | -0.146136 | 0.883814028 | Receptor |
| VAChT | 13 | -0.390943 | 0.695839369 | Transporter |
| 5-HT2A | 7 | -0.716366 | 0.473765383 | Receptor |
| D2 | 2 | -0.771214 | 0.440580097 | Receptor |
| NMDA | 14 | -0.875202 | 0.381464005 | Receptor |
| MOR | 19 | -1.175887 | 0.239640034 | Receptor |
| H3 | 17 | -3.129513 | 0.001750963 | Receptor |

**Table S5** Weights for receptors and transporters in the PLS1 component. Statistically significant level was set as p < .001 after Bonferroni-Hochberg correction.

| **Receptor/Transporter** | **ID** | **Z scores** | **P values** | **Type** |
| --- | --- | --- | --- | --- |
| DAT | 3 | 1.486329 | 0.13719212 | Transporter |
| D1 | 1 | 0.927505 | 0.353664397 | Receptor |
| GABABZ | 16 | 0.323048 | 0.74665889 | Receptor |
| MI | 12 | 0.117458 | 0.906497125 | Receptor |
| alpha4-beta2 | 11 | 0.032146 | 0.97435562 | Receptor |
| 5-HT4 | 8 | -0.135689 | 0.892067144 | Receptor |
| NET | 4 | -0.179791 | 0.857316649 | Transporter |
| 5-HT1B | 6 | -0.188232 | 0.850694788 | Receptor |
| 5-HT1A | 5 | -0.266316 | 0.789995852 | Receptor |
| 5-HTT | 10 | -0.349835 | 0.726462531 | Transporter |
| 5-HT6 | 9 | -0.381933 | 0.702511062 | Receptor |
| D2 | 2 | -0.403481 | 0.686594409 | Receptor |
| 5-HT2A | 7 | -0.783743 | 0.433190926 | Receptor |
| CB1 | 18 | -1.017558 | 0.308888056 | Receptor |
| VAChT | 13 | -1.268602 | 0.204583051 | Transporter |
| mGluR5 | 15 | -1.707917 | 0.087651744 | Receptor |
| NMDA | 14 | -2.295967 | 0.021677768 | Receptor |
| MOR | 19 | -2.452502 | 0.014186659 | Receptor |
| H3 | 17 | -2.686851 | 0.007212912 | Receptor |

**Table S6** Weights for receptors and transporters in the PLS2 component. Statistically significant level was set as p < .001 after Bonferroni-Hochberg correction.

For the gradient 3, we found that the PLS1 explain most 28.10 % variance in this model (p < .01), whilst PLS2 was found to explain 19.39 %. variance (p < .01) here. Further, we estimated the weight z value for each receptor/transporter in PLS1 and PLS2 loading by using Bootstrapping method, respectively. These cortical neurotransmitomic weights for third gradient were then mapped into Schaefer-100 atlas for PLS1 and PLS2 (Figure S9). PLS1 and PLS2 neurotransmitomic weights map was presented in the Figure S9. Rather, we found no correlation between the PLS1/PLS2 weights and case-control z values for third gradient (PLS1: r = .13, p = .17, uncorrected; PLS2: r = .16, p = .11). Weights for PLS1 and PLS2 loading have been tabulated into Table S7-8.

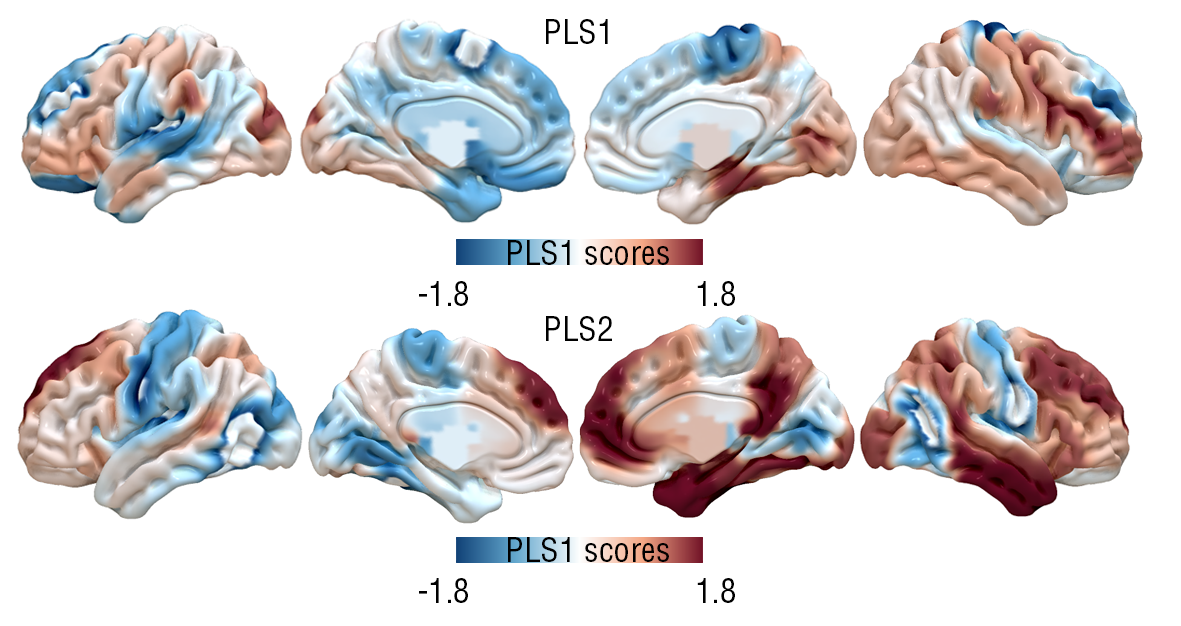

**Figure S9** PLS1 and PLS2 scores maps for PLS models in gradient 3.

| **Receptor/Transporter** | **ID** | **Z scores** | **P values** | **Type** |
| --- | --- | --- | --- | --- |
| NMDA | 14 | 0.535796 | 0.592099547 | Receptor |
| 5-HT6 | 9 | 0.481835 | 0.629923164 | Receptor |
| MI | 12 | 0.161274 | 0.8718776 | Receptor |
| 5-HT2A | 7 | 0.127053 | 0.89889845 | Receptor |
| 5-HT4 | 8 | 0.116201 | 0.907493244 | Receptor |
| GABABZ | 16 | 0.021139 | 0.983134774 | Receptor |
| VAChT | 13 | -0.098095 | 0.921856858 | Transporter |
| NET | 4 | -0.26545 | 0.790662823 | Transporter |
| D1 | 1 | -0.33545 | 0.737285656 | Receptor |
| DAT | 3 | -0.357926 | 0.720398693 | Transporter |
| 5-HT1A | 5 | -0.740089 | 0.459245993 | Receptor |
| 5-HTT | 10 | -0.789719 | 0.429691893 | Transporter |
| alpha4-beta2 | 11 | -1.006961 | 0.313953516 | Receptor |
| mGluR5 | 15 | -1.252003 | 0.210568771 | Receptor |
| D2 | 2 | -1.416053 | 0.156759987 | Receptor |
| CB1 | 18 | -1.449921 | 0.147080551 | Receptor |
| H3 | 17 | -1.562317 | 0.118213329 | Receptor |
| 5-HT1B | 6 | -2.282277 | 0.022472992 | Receptor |
| MOR | 19 | -2.567511 | 0.010243154 | Receptor |

**Table S7** Weights for receptors and transporters in the PLS1 component for third gradient. Statistically significant level was set as p < .001 after Bonferroni-Hochberg correction.

| **Receptor/Transporter** | **ID** | **Z scores** | **P values** | **Type** |
| --- | --- | --- | --- | --- |
| 5-HT6 | 9 | 2.330711 | 0.009884302 | Receptor |
| MI | 12 | 1.525135 | 0.063612714 | Receptor |
| NMDA | 14 | 1.249796 | 0.105687039 | Receptor |
| VAChT | 13 | 1.16228 | 0.122560874 | Transporter |
| 5-HT2A | 7 | 1.100278 | 0.135605507 | Receptor |
| GABABZ | 16 | 0.943863 | 0.172619831 | Receptor |
| 5-HT4 | 8 | 0.880081 | 0.189407716 | Receptor |
| alpha4-beta2 | 11 | 0.6629 | 0.253697301 | Receptor |
| D1 | 1 | 0.524668 | 0.299907003 | Receptor |
| 5-HT1A | 5 | 0.497672 | 0.309357623 | Receptor |
| NET | 4 | 0.125067 | 0.450235254 | Transporter |
| 5-HTT | 10 | 0.023839 | 0.490490516 | Transporter |
| mGluR5 | 15 | -0.081431 | 0.467549598 | Receptor |
| DAT | 3 | -0.209395 | 0.417069948 | Transporter |
| CB1 | 18 | -0.344261 | 0.365325006 | Receptor |
| H3 | 17 | -1.649225 | 0.049550774 | Receptor |
| D2 | 2 | -1.947804 | 0.02571921 | Receptor |
| 5-HT1B | 6 | -1.994783 | 0.023033276 | Receptor |
| MOR | 19 | -2.233502 | 0.012757929 | Receptor |

**Table S8** Weights for receptors and transporters in the PLS2 component for third gradient. Statistically significant level was set as p < .001 after Bonferroni-Hochberg correction.

6. **Transcriptomic signatures for gradient changes of ADHD**

By using the PLS model, we reveled that the PLS1 and PLS2 component explained 17.94% variances of principal gradient. Further, we used Bootstrapping method (n = 10,000) to estimate the weights (z scores) for each gene in these PLS loading. For screening associated genes, we re-ranked these genes by descending order, and calculated the p value for each weight with Bonferroni-Hochberg correction at p < .001 (Z > 5.2 or Z < -5.2). By using these criterion, a total of 134 genes (58 genes with positive weight, PLS1+; 76 genes with negative weight, PLS1-) in PLS1 component and 34 genes (6 genes with positive weight, PLS2+; 26 genes with negative weight, PLS2-) have been revealed. We have provided unfolded tables underneaths for reporting both gene lists.

| **ID** | **Gene Symbol** | **Z scores** | **P values** |
| --- | --- | --- | --- |
| 7908 | DYDC2 | 7.182961 | 6.82E-13 |
| 9318 | ATOH7 | 6.868112 | 6.51E-12 |
| 4931 | MMD | 6.857208 | 7.02E-12 |
| 148 | BIRC3 | 6.757306 | 1.41E-11 |
| 2164 | PKIA | 6.588036 | 4.46E-11 |
| 9746 | KCTD4 | 6.564263 | 5.23E-11 |
| 1063 | GLRA2 | 6.397678 | 1.58E-10 |
| 6497 | SULF2 | 6.369096 | 1.90E-10 |
| 8445 | TMEM200A | 6.299226 | 2.99E-10 |
| 8482 | LRRC56 | 6.278778 | 3.41E-10 |
| 2575 | SLA | 6.230166 | 4.66E-10 |
| 190 | ASCL2 | 6.212186 | 5.23E-10 |
| 6312 | TRIM36 | 6.194948 | 5.83E-10 |
| 2257 | PTGER3 | 6.183347 | 6.28E-10 |
| 216 | ATP2B4 | 6.172336 | 6.73E-10 |
| 8541 | DACH2 | 6.090811 | 1.12E-09 |
| 8089 | HIST1H2BK | 6.027722 | 1.66E-09 |
| 886 | F12 | 5.952975 | 2.63E-09 |
| 2617 | SLIT1 | 5.817381 | 5.98E-09 |
| 9926 | CPLX3 | 5.812645 | 6.15E-09 |
| 9996 | OST4 | 5.795297 | 6.82E-09 |
| 5351 | RRP7A | 5.644785 | 1.65E-08 |
| 1106 | GPR6 | 5.58285 | 2.37E-08 |
| 5774 | ZCCHC17 | 5.563766 | 2.64E-08 |
| 2497 | S100A10 | 5.541479 | 3.00E-08 |
| 8282 | DAPL1 | 5.498573 | 3.83E-08 |
| 9531 | RIIAD1 | 5.467808 | 4.56E-08 |
| 9693 | WDR86 | 5.452951 | 4.95E-08 |
| 1991 | PDYN | 5.434546 | 5.49E-08 |
| 6607 | NIT2 | 5.424635 | 5.81E-08 |
| 9248 | LYRM9 | 5.406895 | 6.41E-08 |
| 9920 | NBDY | 5.380681 | 7.42E-08 |
| 5157 | ERC2 | 5.379818 | 7.46E-08 |
| 1371 | IL13RA2 | 5.371958 | 7.79E-08 |
| 3800 | RIPOR2 | 5.37059 | 7.85E-08 |
| 7803 | KIRREL2 | 5.330366 | 9.80E-08 |
| 6116 | PID1 | 5.317341 | 1.05E-07 |
| 3070 | ZNF226 | 5.306918 | 1.11E-07 |
| 2131 | PPP1R1A | 5.305642 | 1.12E-07 |
| 6293 | TMEM176A | 5.30449 | 1.13E-07 |
| 7766 | TMTC1 | 5.276013 | 1.32E-07 |
| 3418 | CACNA1H | 5.27242 | 1.35E-07 |
| 5902 | GNG2 | 5.26952 | 1.37E-07 |
| 2521 | SCN9A | 5.26712 | 1.39E-07 |
| 9085 | AKAP14 | 5.264738 | 1.40E-07 |
| 9377 | SEMA3D | 5.261639 | 1.43E-07 |
| 2311 | RAB3B | 5.259272 | 1.45E-07 |
| 2338 | RAP2B | 5.258113 | 1.46E-07 |
| 1853 | NOV | 5.239631 | 1.61E-07 |
| 6823 | SH3RF1 | 5.224503 | 1.75E-07 |
| 2315 | RAB27B | 5.223228 | 1.76E-07 |
| 3598 | CPNE6 | 5.220939 | 1.78E-07 |
| 920 | GPC4 | 5.213146 | 1.86E-07 |
| 9380 | SLC17A8 | 5.187502 | 2.13E-07 |
| 1964 | PDE2A | 5.178578 | 2.24E-07 |
| 1208 | HBQ1 | 5.168563 | 2.36E-07 |
| 6199 | NUDT11 | 5.161822 | 2.45E-07 |
| 5300 | CPAMD8 | 5.150796 | 2.59E-07 |

**Table S9** Gene list for statistically significant association with principal gradients changes in the PLS1+ component. P values were estimated by using Z sampling distribution.

| **ID** | **Gene Symbol** | **Z scores** | **P values** |
| --- | --- | --- | --- |
| 8447 | RNF157 | 8447 | -5.179801 |
| 5634 | MYO15A | 5634 | -5.240591 |
| 6883 | C6orf47 | 6883 | -5.241308 |
| 6692 | SLC45A4 | 6692 | -5.250074 |
| 8733 | ACVR1C | 8733 | -5.250109 |
| 956 | FLT3 | 956 | -5.252495 |
| 3533 | VAPA | 3533 | -5.253716 |
| 6898 | ZBED5 | 6898 | -5.257201 |
| 778 | EDNRA | 778 | -5.261859 |
| 359 | CAMK2G | 359 | -5.289353 |
| 486 | CHGA | 486 | -5.314568 |
| 7831 | ARID5B | 7831 | -5.327025 |
| 5138 | OSBPL3 | 5138 | -5.334835 |
| 9973 | PCP4L1 | 9973 | -5.335447 |
| 5811 | MPC1 | 5811 | -5.340607 |
| 4358 | PPARGC1A | 4358 | -5.351762 |
| 9130 | DENND1B | 9130 | -5.467565 |
| 5305 | GLS2 | 5305 | -5.471878 |
| 1350 | IGFBP2 | 1350 | -5.47325 |
| 5165 | ABHD12 | 5165 | -5.484875 |
| 3907 | HS3ST1 | 3907 | -5.491524 |
| 4239 | CDC42EP3 | 4239 | -5.49882 |
| 2294 | PVALB | 2294 | -5.501187 |
| 2007 | PFKFB2 | 2007 | -5.530285 |
| 8757 | DCBLD2 | 8757 | -5.555034 |
| 2061 | PLCB4 | 2061 | -5.563411 |
| 217 | ATP4A | 217 | -5.57997 |
| 3062 | ZSCAN9 | 3062 | -5.601548 |
| 6654 | PLXDC1 | 6654 | -5.613183 |
| 508 | CKMT1B | 508 | -5.638111 |
| 9812 | ANKRD34C | 9812 | -5.641329 |
| 771 | ECM1 | 771 | -5.684385 |
| 9580 | INTS4P1 | 9580 | -5.692733 |
| 7290 | BHLHE41 | 7290 | -5.71931 |
| 3078 | ZYX | 3078 | -5.729601 |
| 7483 | PLEKHH3 | 7483 | -5.749256 |
| 4121 | SYCP2 | 4121 | -5.77284 |
| 8415 | SLC35A4 | 8415 | -5.778586 |
| 6552 | EIF5A2 | 6552 | -5.794553 |
| 3499 | P2RX6 | 3499 | -5.826287 |
| 7366 | ZMAT4 | 7366 | -5.859169 |
| 7713 | SCRT1 | 7713 | -5.87046 |
| 9131 | IFNLR1 | 9131 | -5.897463 |
| 924 | FGF9 | 924 | -5.907952 |
| 5937 | PRMT7 | 5937 | -5.925703 |
| 4410 | CCNI | 4410 | -5.951056 |
| 6007 | KCTD9 | 6007 | -6.067214 |
| 1147 | NR3C1 | 1147 | -6.090213 |
| 6534 | SERTAD4 | 6534 | -6.092176 |
| 2370 | RET | 2370 | -6.101001 |
| 3369 | FGF18 | 3369 | -6.111956 |
| 5364 | SLCO4A1 | 5364 | -6.115447 |
| 6442 | HR | 6442 | -6.204223 |
| 8401 | GLCCI1 | 8401 | -6.215447 |
| 5517 | ZBTB21 | 5517 | -6.235313 |
| 6228 | SLC47A1 | 6228 | -6.256673 |
| 4051 | RCAN2 | 4051 | -6.279235 |
| 1480 | KCNS1 | 1480 | -6.295519 |
| 4807 | KIAA1107 | 4807 | -6.296776 |
| 2513 | SCN1A | 2513 | -6.355923 |
| 1958 | PCSK1 | 1958 | -6.391459 |
| 5456 | FAM216A | 5456 | -6.393551 |
| 9398 | FBXO33 | 9398 | -6.41137 |
| 7510 | PIF1 | 7510 | -6.447735 |
| 6829 | MYH7B | 6829 | -6.558421 |
| 9265 | CMYA5 | 9265 | -6.640444 |
| 9605 | RAB37 | 9605 | -6.683739 |
| 8465 | FBXO32 | 8465 | -7.016892 |
| 1511 | LAG3 | 1511 | -7.070116 |
| 9656 | FMN1 | 9656 | -7.176766 |
| 3523 | KCNAB3 | 3523 | -7.422486 |
| 1428 | ITPR1 | 1428 | -7.591109 |
| 124 | ANK1 | 124 | -7.769391 |
| 303 | KLF9 | 303 | -7.848392 |
| 9860 | HAPLN4 | 9860 | -7.912125 |
| 8457 | OSBPL6 | 8457 | -8.054201 |

**Table S10** Gene list for statistically significant association with principal gradients changes in the PLS1- component. P values were estimated by using Z sampling distribution.

| **ID** | **Gene Symbol** | **Z scores** | **P values** |
| --- | --- | --- | --- |
| 3446 | NOL3 | 5.687012 | 1.29281E-08 |
| 5998 | FAM20A | 5.620732 | 1.9015E-08 |
| 5124 | MOXD1 | 5.518411 | 3.42079E-08 |
| 6592 | LXN | 5.503236 | 3.72883E-08 |
| 8163 | HS6ST2 | 5.306792 | 1.11571E-07 |
| 135 | ANXA6 | 5.171803 | 2.31846E-07 |

**Table S11** Gene list for statistically significant association with principal gradients changes in the PLS2+ component. P values were estimated by using Z sampling distribution.

| **ID** | **Gene Symbol** | **Z scores** | **P values** |
| --- | --- | --- | --- |
| 4525 | NXPH2 | -5.244292 | 1.56884E-07 |
| 2076 | PLTP | -5.259603 | 1.44367E-07 |
| 5288 | DKK2 | -5.298105 | 1.17011E-07 |
| 9334 | SLC39A12 | -5.310252 | 1.09474E-07 |
| 844 | EPHB2 | -5.35663 | 8.47885E-08 |
| 402 | CD6 | -5.38117 | 7.40033E-08 |
| 2818 | THBD | -5.381442 | 7.38915E-08 |
| 5051 | MGAT4C | -5.382555 | 7.34359E-08 |
| 6516 | PCDHB4 | -5.392064 | 6.96529E-08 |
| 4000 | TOB1 | -5.455329 | 4.88823E-08 |
| 9911 | CLEC18B | -5.510857 | 3.57091E-08 |
| 2380 | RFX3 | -5.560573 | 2.6889E-08 |
| 122 | ANGPT1 | -5.615241 | 1.96288E-08 |
| 633 | CTSH | -5.63165 | 1.78494E-08 |
| 3016 | ZIC1 | -5.717789 | 1.07919E-08 |
| 890 | FABP5 | -5.725956 | 1.02853E-08 |
| 1274 | HPCAL1 | -5.834014 | 5.41096E-09 |
| 3094 | SCG2 | -5.89139 | 3.82961E-09 |
| 8763 | RTP1 | -5.905038 | 3.52565E-09 |
| 8927 | CBLN2 | -6.067095 | 1.30245E-09 |
| 5215 | HSPB8 | -6.071726 | 1.26543E-09 |
| 1712 | MT1F | -6.331317 | 2.43077E-10 |
| 2090 | PRRX1 | -6.425161 | 1.3173E-10 |
| 5819 | ASB2 | -6.440427 | 1.19138E-10 |
| 80 | AEBP1 | -7.025917 | 2.12674E-12 |
| 445 | CDH13 | -7.454011 | 9.05942E-14 |

**Table S12** Gene list for statistically significant association with principal gradients changes in the PLS2- component. P values were estimated by using Z sampling distribution.

7. **Association of single-gene expression level to gradient changes of ADHD**

Rather gene lists in PLS components, we estimated the correlation between each gene that survived from Bonferroni-Hochberg correction at p < .001 (Z > 5.2 or Z < -5.2) and case-control z scores. Results revealed that the principal gradient changes of ADHD were associated with single-gene expression level. Full results have been sorted into Table S13-16.

| **ID** | **Gene Symbol** | **R values** | **P values** |
| --- | --- | --- | --- |
| 1106 | GPR6 | 0.327063775 | 2.10E-08 |
| 5902 | GNG2 | 0.296014804 | 4.54E-07 |
| 2164 | PKIA | 0.285400381 | 1.20E-06 |
| 5300 | CPAMD8 | 0.285054262 | 1.24E-06 |
| 2617 | SLIT1 | 0.282031209 | 1.62E-06 |
| 4931 | MMD | 0.273972334 | 3.27E-06 |
| 2497 | S100A10 | 0.260925428 | 9.71E-06 |
| 6312 | TRIM36 | 0.256117833 | 1.43E-05 |
| 6293 | TMEM176A | 0.255024216 | 1.56E-05 |
| 9248 | LYRM9 | 0.253754376 | 1.73E-05 |
| 2311 | RAB3B | 0.2529843 | 1.83E-05 |
| 7908 | DYDC2 | 0.247442607 | 2.82E-05 |
| 7766 | TMTC1 | 0.24728974 | 2.86E-05 |
| 8445 | TMEM200A | 0.24605937 | 3.14E-05 |
| 9693 | WDR86 | 0.2419781 | 4.28E-05 |
| 8282 | DAPL1 | 0.24047306 | 4.79E-05 |
| 8089 | HIST1H2BK | 0.240425409 | 4.81E-05 |
| 9920 | NBDY | 0.239472516 | 5.16E-05 |
| 190 | ASCL2 | 0.238271912 | 5.64E-05 |
| 6823 | SH3RF1 | 0.237748543 | 5.86E-05 |
| 9746 | KCTD4 | 0.235035714 | 7.15E-05 |
| 148 | BIRC3 | 0.23380083 | 7.83E-05 |
| 9318 | ATOH7 | 0.231460902 | 9.27E-05 |
| 3070 | ZNF226 | 0.228494049 | 1.15E-04 |
| 6497 | SULF2 | 0.227227891 | 1.25E-04 |
| 9531 | RIIAD1 | 0.227130505 | 1.26E-04 |
| 8541 | DACH2 | 0.224020153 | 1.57E-04 |
| 2338 | RAP2B | 0.223431105 | 1.63E-04 |
| 3598 | CPNE6 | 0.222679564 | 1.72E-04 |
| 6607 | NIT2 | 0.222059209 | 1.80E-04 |
| 9380 | SLC17A8 | 0.218196187 | 2.34E-04 |
| 9085 | AKAP14 | 0.217563014 | 2.44E-04 |
| 7803 | KIRREL2 | 0.217401653 | 2.47E-04 |
| 5774 | ZCCHC17 | 0.21722313 | 2.50E-04 |
| 1063 | GLRA2 | 0.216805604 | 2.57E-04 |
| 1964 | PDE2A | 0.214741623 | 2.95E-04 |
| 8482 | LRRC56 | 0.213491948 | 3.21E-04 |
| 2315 | RAB27B | 0.209307161 | 4.22E-04 |
| 2521 | SCN9A | 0.208381486 | 4.48E-04 |
| 2575 | SLA | 0.206650416 | 5.01E-04 |
| 3418 | CACNA1H | 0.20544692 | 5.41E-04 |
| 9926 | CPLX3 | 0.202342473 | 6.59E-04 |
| 2257 | PTGER3 | 0.201226291 | 7.07E-04 |
| 9996 | OST4 | 0.199650169 | 7.80E-04 |
| 1991 | PDYN | 0.19944899 | 7.90E-04 |
| 216 | ATP2B4 | 0.196531607 | 9.46E-04 |
| 886 | F12 | 0.194934784 | 1.04E-03 |
| 1853 | NOV | 0.193021884 | 1.17E-03 |
| 2131 | PPP1R1A | 0.192275087 | 1.22E-03 |
| 3800 | RIPOR2 | 0.189345065 | 1.46E-03 |
| 1371 | IL13RA2 | 0.1827776 | 2.14E-03 |
| 920 | GPC4 | 0.182228678 | 2.20E-03 |
| 5157 | ERC2 | 0.181504629 | 2.30E-03 |
| 5351 | RRP7A | 0.181492295 | 2.30E-03 |
| 9377 | SEMA3D | 0.179866112 | 2.52E-03 |
| 1208 | HBQ1 | 0.179419919 | 2.58E-03 |
| 6116 | PID1 | 0.178801427 | 2.68E-03 |
| 6199 | NUDT11 | 0.170257944 | 4.28E-03 |

**Table S13** Associations between single-gene expression level and case-control z values of principal gradients change in the PLS1+ component. The order of these genes have been descended by correlation strength.

| **ID** | **Gene Symbol** | **R values** | **P values** |
| --- | --- | --- | --- |
| 9265 | CMYA5 | -0.329084121 | 1.70E-08 |
| 4410 | CCNI | -0.303629767 | 2.21E-07 |
| 3369 | FGF18 | -0.293553148 | 5.71E-07 |
| 2007 | PFKFB2 | -0.288770455 | 8.85E-07 |
| 4051 | RCAN2 | -0.280328297 | 1.88E-06 |
| 8465 | FBXO32 | -0.277274571 | 2.46E-06 |
| 3499 | P2RX6 | -0.275323338 | 2.91E-06 |
| 7831 | ARID5B | -0.274898908 | 3.02E-06 |
| 8733 | ACVR1C | -0.273267767 | 3.47E-06 |
| 9812 | ANKRD34C | -0.265290642 | 6.79E-06 |
| 1511 | LAG3 | -0.263878216 | 7.63E-06 |
| 6007 | KCTD9 | -0.262683526 | 8.42E-06 |
| 303 | KLF9 | -0.261446049 | 9.31E-06 |
| 5937 | PRMT7 | -0.261426454 | 9.33E-06 |
| 5517 | ZBTB21 | -0.26084327 | 9.78E-06 |
| 9580 | INTS4P1 | -0.260740407 | 9.86E-06 |
| 9656 | FMN1 | -0.260637347 | 9.94E-06 |
| 6654 | PLXDC1 | -0.258619783 | 1.17E-05 |
| 3078 | ZYX | -0.249945295 | 2.33E-05 |
| 5811 | MPC1 | -0.24894834 | 2.51E-05 |
| 9398 | FBXO33 | -0.248254321 | 2.65E-05 |
| 1428 | ITPR1 | -0.245878833 | 3.18E-05 |
| 6829 | MYH7B | -0.243376789 | 3.85E-05 |
| 5165 | ABHD12 | -0.240967318 | 4.62E-05 |
| 6898 | ZBED5 | -0.240232073 | 4.88E-05 |
| 4807 | KIAA1107 | -0.239560867 | 5.13E-05 |
| 8447 | RNF157 | -0.239398878 | 5.19E-05 |
| 1350 | IGFBP2 | -0.239280301 | 5.23E-05 |
| 486 | CHGA | -0.23914234 | 5.29E-05 |
| 7510 | PIF1 | -0.238371737 | 5.60E-05 |
| 9130 | DENND1B | -0.237998165 | 5.76E-05 |
| 6692 | SLC45A4 | -0.2373226 | 6.05E-05 |
| 956 | FLT3 | -0.236188078 | 6.58E-05 |
| 8457 | OSBPL6 | -0.235229551 | 7.05E-05 |
| 9860 | HAPLN4 | -0.235198048 | 7.07E-05 |
| 5456 | FAM216A | -0.234275202 | 7.56E-05 |
| 508 | CKMT1B | -0.233829369 | 7.81E-05 |
| 9605 | RAB37 | -0.231532942 | 9.22E-05 |
| 3533 | VAPA | -0.22795949 | 0.000118981 |
| 2294 | PVALB | -0.226645306 | 0.000130542 |
| 7290 | BHLHE41 | -0.226596538 | 0.000130991 |
| 8415 | SLC35A4 | -0.225040669 | 0.000146086 |
| 3523 | KCNAB3 | -0.223611846 | 0.000161369 |
| 124 | ANK1 | -0.222887804 | 0.000169673 |
| 9131 | IFNLR1 | -0.221198012 | 0.000190634 |
| 5634 | MYO15A | -0.220842661 | 0.00019534 |
| 5138 | OSBPL3 | -0.220697011 | 0.000197299 |
| 1480 | KCNS1 | -0.219676684 | 0.000211553 |
| 4239 | CDC42EP3 | -0.214259909 | 0.000304733 |
| 7483 | PLEKHH3 | -0.213780939 | 0.000314592 |
| 359 | CAMK2G | -0.210757213 | 0.000384015 |
| 1147 | NR3C1 | -0.210679678 | 0.000385969 |
| 6228 | SLC47A1 | -0.209737817 | 0.000410462 |
| 6552 | EIF5A2 | -0.208036823 | 0.000458389 |
| 6883 | C6orf47 | -0.206940221 | 0.000491987 |
| 4358 | PPARGC1A | -0.206870687 | 0.000494193 |
| 2370 | RET | -0.206616726 | 0.000502327 |
| 924 | FGF9 | -0.205074307 | 0.000554452 |
| 4121 | SYCP2 | -0.204963091 | 0.000558398 |
| 6442 | HR | -0.204433778 | 0.000577536 |
| 1958 | PCSK1 | -0.204120534 | 0.000589147 |
| 2061 | PLCB4 | -0.202148086 | 0.000667362 |
| 771 | ECM1 | -0.202108377 | 0.000669031 |
| 6534 | SERTAD4 | -0.200666877 | 0.000732294 |
| 2513 | SCN1A | -0.200546583 | 0.000737815 |
| 8401 | GLCCI1 | -0.199485722 | 0.000788202 |
| 3907 | HS3ST1 | -0.197416941 | 0.000895714 |
| 7366 | ZMAT4 | -0.196525134 | 0.000946103 |
| 5364 | SLCO4A1 | -0.19514578 | 0.001029203 |
| 8757 | DCBLD2 | -0.193727233 | 0.001121628 |
| 217 | ATP4A | -0.191381523 | 0.001291352 |
| 7713 | SCRT1 | -0.19134715 | 0.001294005 |
| 5305 | GLS2 | -0.191003483 | 0.001320814 |
| 778 | EDNRA | -0.190652356 | 0.001348729 |
| 3062 | ZSCAN9 | -0.186437313 | 0.0017288 |
| 9973 | PCP4L1 | -0.176569434 | 0.003029955 |

**Table S14** Associations between single-gene expression level and case-control z values of principal gradients change in the PLS1- component. The order of these genes have been descended by correlation strength.

| **ID** | **Gene Symbol** | **R values** | **P values** |
| --- | --- | --- | --- |
| 5124 | MOXD1 | 0.339889149 | 5.33E-09 |
| 6592 | LXN | 0.250013399 | 2.31E-05 |
| 3446 | NOL3 | 0.218380561 | 2.31E-04 |
| 8163 | HS6ST2 | 0.117826509 | 4.89E-02 |
| 135 | ANXA6 | 0.091470738 | 1.27E-01 |
| 5998 | FAM20A | 0.068697667 | 2.52E-01 |

**Table S15** Associations between single-gene expression level and case-control z values of principal gradients change in the PLS2+ component. The order of these genes have been descended by correlation strength.

| **ID** | **Gene Symbol** | **R values** | **P values** |
| --- | --- | --- | --- |
| 844 | EPHB2 | -0.31290031 | 8.93E-08 |
| 4525 | NXPH2 | -0.282360081 | 1.57E-06 |
| 5215 | HSPB8 | -0.27894429 | 2.12E-06 |
| 3016 | ZIC1 | -0.263139022 | 8.11E-06 |
| 6516 | PCDHB4 | -0.261055531 | 9.61E-06 |
| 5819 | ASB2 | -0.236437787 | 6.46E-05 |
| 5288 | DKK2 | -0.235010726 | 7.17E-05 |
| 4000 | TOB1 | -0.226749887 | 1.30E-04 |
| 2090 | PRRX1 | -0.219201554 | 2.19E-04 |
| 445 | CDH13 | -0.219083279 | 2.20E-04 |
| 8927 | CBLN2 | -0.214429033 | 3.01E-04 |
| 80 | AEBP1 | -0.203471359 | 6.14E-04 |
| 2818 | THBD | -0.202778687 | 6.41E-04 |
| 3094 | SCG2 | -0.182178392 | 2.21E-03 |
| 122 | ANGPT1 | -0.163637672 | 6.06E-03 |
| 5051 | MGAT4C | -0.155026024 | 9.37E-03 |
| 633 | CTSH | -0.154854893 | 9.45E-03 |
| 9911 | CLEC18B | -0.153225126 | 1.02E-02 |
| 2380 | RFX3 | -0.151724364 | 1.10E-02 |
| 1274 | HPCAL1 | -0.145692088 | 1.47E-02 |
| 9334 | SLC39A12 | -0.113601441 | 5.76E-02 |
| 2076 | PLTP | -0.10185336 | 8.89E-02 |
| 8763 | RTP1 | -0.097840813 | 1.02E-01 |
| 890 | FABP5 | -0.088501846 | 1.40E-01 |
| 402 | CD6 | -0.075470186 | 2.08E-01 |
| 1712 | MT1F | -0.054148993 | 3.67E-01 |

**Table S16** Associations between single-gene expression level and case-control z values of principal gradients change in the PLS2- component. The order of these genes have been descended by correlation strength.

8. **Association of these PLS gene sets to other neurodevelopmental psychiatric disorders**

To examine whether these PLS gene sets were specific for ADHD patients, we extracted ISH genes relating to autism disorder, bipolar disorder and intellectual disability. Further, we overlapped the PLS gene sets of gradient-derived phenotype for ISH genes of these neurodevelopmental psychiatric disorders. However, Only GLRA2 and PVALB that were categorized in autism disorder overlapped in the PLS1+ (1/58, 1.72%) and PLS1- (1/76, 1.31%) gene set. Further, we correlated regional expression levels of these non-overlapping genes that categorized into autism disorder, bipolar disorder or intellectual disability to case-control z-scores for principal gradient in ADHD patients, but reveal the null associations for them (p < .001). Full results have been tabulated into the Table S17.

| **Gene Symbol** | **EntrezID** | **R values** | **P values** | **Disorders in ISH** |
| --- | --- | --- | --- | --- |
| EXT1 | 2131 | 0.24728974 | 2.86E-05 | autism |
| PVALB | 5816 | 0.229857823 | 0.000103966 | autism |
| PIK3CG | 5294 | 0.176561786 | 0.00303124 | autism |
| CXCL14 | 9547 | 0.17123862 | 0.00405555 | autism |
| GABRA5 | 2558 | 0.15722715 | 0.008399956 | autism |
| DLX5 | 1749 | 0.130306676 | 0.029257715 | autism |
| PLAUR | 5329 | 0.122198249 | 0.041024319 | autism |
| DVL1 | 1855 | 0.117108569 | 0.050279816 | autism |
| SLC1A2 | 6506 | 0.115462853 | 0.053621798 | autism |
| TSC2 | 7249 | 0.111524655 | 0.062373131 | autism |
| OXTR | 5021 | 0.109757709 | 0.066664954 | autism |
| CALB1 | 793 | 0.107970456 | 0.071249195 | autism |
| PRKCB | 5579 | 0.098626095 | 0.099565204 | autism |
| SST | 6750 | 0.098160682 | 0.101180908 | autism |
| MET | 4233 | 0.087812103 | 0.142747236 | autism |
| ACCN1 | 40 | 0.062719444 | 0.295638553 | autism |
| GRM8 | 2918 | 0.057760806 | 0.335545681 | autism |
| GABRA4 | 2557 | 0.047328606 | 0.430194764 | autism |
| TDO2 | 6999 | 0.047142953 | 0.432007778 | autism |
| LRP8 | 7804 | 0.046766761 | 0.435694989 | autism |
| GABRG1 | 2565 | 0.045248054 | 0.450762735 | autism |
| GLRB | 2743 | 0.038292599 | 0.523392043 | autism |
| ASL | 435 | 0.036878214 | 0.538859152 | autism |
| OMG | 4974 | 0.030090611 | 0.616108406 | autism |
| DLX1 | 1745 | 0.024844837 | 0.678919681 | autism |
| VIP | 7432 | 0.022061711 | 0.713204087 | autism |
| HTR7 | 3363 | 0.019225647 | 0.748743142 | autism |
| CACNA1C | 775 | 0.010820612 | 0.856949427 | autism |
| GLO1 | 2739 | 0.006627747 | 0.9120854 | autism |
| SLC6A12 | 6539 | 0.000557055 | 0.992596046 | autism |
| INPP1 | 3628 | -0.000885812 | 0.988226709 | autism |
| ADAM23 | 8745 | -0.01113072 | 0.852895612 | autism |
| GABRG3 | 2567 | -0.020560102 | 0.731949448 | autism |
| ADCYAP1 | 116 | -0.021913175 | 0.715050885 | autism |
| GRPR | 2925 | -0.022406353 | 0.70892545 | autism |
| DHCR7 | 1717 | -0.024815013 | 0.679283744 | autism |
| TSC1 | 7248 | -0.029078915 | 0.628024634 | autism |
| NEFL | 4747 | -0.030450638 | 0.611891785 | autism |
| GAD1 | 2571 | -0.030916279 | 0.606457152 | autism |
| GLRA2 | 2742 | -0.034701595 | 0.563098504 | autism |
| MBP | 4155 | -0.038843228 | 0.517432473 | autism |
| RORB | 6096 | -0.039740707 | 0.507794296 | autism |
| GABRB1 | 2560 | -0.053907064 | 0.368835881 | autism |
| PDE1A | 5136 | -0.059293762 | 0.322855945 | autism |
| SLC6A1 | 6529 | -0.078297749 | 0.191444709 | autism |
| EN2 | 2020 | -0.090634116 | 0.1302975 | autism |
| RELN | 5649 | -0.106879835 | 0.074170689 | autism |
| AVPR1A | 552 | -0.108537697 | 0.069767196 | autism |
| GABRA2 | 2555 | -0.126037476 | 0.035031945 | autism |
| NRCAM | 4897 | -0.127212304 | 0.033353792 | autism |
| AIF1 | 199 | -0.127369274 | 0.033134839 | autism |
| PTEN | 5728 | -0.129642593 | 0.030098504 | autism |
| CA3 | 761 | -0.137237367 | 0.021617272 | autism |
| PCP4 | 5121 | -0.137428743 | 0.021433485 | autism |
| CALB2 | 794 | -0.139369838 | 0.019644553 | autism |
| GLRA3 | 8001 | -0.142251217 | 0.017228509 | autism |
| MFGE8 | 4240 | -0.144873744 | 0.01525913 | autism |
| NPY1R | 4886 | -0.145916986 | 0.014532294 | autism |
| PTGS2 | 5743 | -0.150914849 | 0.011455451 | autism |
| VIPR2 | 7434 | -0.156248376 | 0.008820245 | autism |
| CTGF | 1490 | -0.157127647 | 0.008441854 | autism |
| NRXN1 | 9378 | -0.194095202 | 0.001096947 | autism |
| WNT2 | 7472 | -0.199176748 | 0.000803464 | autism |
| CLOCK | 9575 | 0.081343543 | 0.174686213 | bipolar |
| XBP1 | 7494 | 0.002326824 | 0.969080874 | bipolar |
| ADRBK2 | 157 | -0.010693827 | 0.858607889 | bipolar |
| AMMECR1 | 9949 | 0.135030307 | 0.023837368 | Intell Disab |
| DLG3 | 1741 | 0.114786427 | 0.055047929 | Intell Disab |
| PDHX | 8050 | 0.106576866 | 0.074999323 | Intell Disab |
| UBE3A | 7337 | 0.084175541 | 0.160104737 | Intell Disab |
| ACTB | 60 | 0.082723676 | 0.1674615 | Intell Disab |
| SMS | 6611 | 0.057159994 | 0.340605134 | Intell Disab |
| GDI1 | 2664 | 0.046737042 | 0.435987051 | Intell Disab |
| DCX | 1641 | 0.039703292 | 0.508194223 | Intell Disab |
| FGD1 | 2245 | 0.02797443 | 0.641144686 | Intell Disab |
| GRIK2 | 2898 | 0.021086454 | 0.725359928 | Intell Disab |
| ZNF41 | 7592 | 0.00279255 | 0.962896307 | Intell Disab |
| VLDLR | 7436 | -0.012588626 | 0.833890905 | Intell Disab |
| SMCX | 8242 | -0.01980012 | 0.741498519 | Intell Disab |
| SYP | 6855 | -0.021568442 | 0.719343477 | Intell Disab |
| PRSS12 | 8492 | -0.024336886 | 0.685130373 | Intell Disab |
| OPHN1 | 4983 | -0.02951564 | 0.622868607 | Intell Disab |
| PAK3 | 5063 | -0.04110665 | 0.49330709 | Intell Disab |
| MECP2 | 4204 | -0.046761593 | 0.435745767 | Intell Disab |
| FGFR1 | 2260 | -0.051435667 | 0.391223428 | Intell Disab |
| FMR1 | 2332 | -0.058430505 | 0.329963161 | Intell Disab |
| IGF1 | 3479 | -0.063484603 | 0.289774419 | Intell Disab |
| ATRX | 546 | -0.074251655 | 0.2154862 | Intell Disab |
| L1CAM | 3897 | -0.081678462 | 0.17291207 | Intell Disab |
| RPS6KA3 | 6197 | -0.097993768 | 0.101765393 | Intell Disab |
| SLC6A8 | 6535 | -0.104423054 | 0.081109078 | Intell Disab |
| LARGE | 9215 | -0.112575557 | 0.05993017 | Intell Disab |
| CTNND2 | 1501 | -0.121112372 | 0.042868221 | Intell Disab |
| IGBP1 | 3476 | -0.122810663 | 0.040014191 | Intell Disab |
| AGTR2 | 186 | -0.124009906 | 0.038096845 | Intell Disab |
| ARHGEF6 | 9459 | -0.152908362 | 0.010398584 | Intell Disab |
| CA2 | 760 | -0.198311344 | 0.000847674 | Intell Disab |
| ZFHX1B | 9839 | -0.223369161 | 0.000164109 | Intell Disab |

**Table S17** Correlation of non-overlapping genes in ISH dataset relating to other neurodevelopmental psychiatric disorders and regional differences of principal gradient in the ADHD. Genes for each disorder were re-ranked by descending order. Intell Disab = Intelligence Disability

9. **Decoding the brain network associations with PLS gene sets**

On the basis of GAMBA tool, we decoded PLS gene sets for ADHD-specific gradient differences into brain networks that defined by Yeo-7 atlas^12^, including visual network (VIS), sensory/motor network (SMN), dorsal attention network (DAN), ventral attention network (VAN), limbic network (LIB), frontoparital network (FPN) and default model network (DMN). The general linear regression models were used to fit the PLS gene sets to brain network properties, respectively. To control the inflation of false-positive error, the Bonferroni-Holm FDR corrections have been performed. The standardized beta values indicated the slope of corresponding models. Full results have been sorted into Table S18-19.

| **Components** | **Network** | **Standardized Beta** | **Significance** |
| --- | --- | --- | --- |
| PLS1+ | Visual | -0.502738089 | *** |
| PLS1+ | Somatomotor | -0.273899029 | - |
| PLS1+ | Dorsal_attention | -0.105894459 | - |
| PLS1+ | Ventral_attention | 0.253915953 | - |
| PLS1+ | Limbic | 0.494423913 | *** |
| PLS1+ | Frontal_parietal | 0.05501908 | - |
| PLS1+ | Default_mode | 0.293883665 | - |
| PLS1- | Visual | 0.521397833 | *** |
| PLS1- | Somatomotor | 0.307009181 | - |
| PLS1- | Dorsal_attention | 0.179777108 | - |
| PLS1- | Ventral_attention | -0.213869162 | - |
| PLS1- | Limbic | -0.5610615 | *** |
| PLS1- | Frontal_parietal | -0.125150794 | - |
| PLS1- | Default_mode | -0.324448261 | - |

**Table S18** Association of gene set in the PLS1 component for brain network properties in healthy brain. Statistical significance was set as p < .05 with Bonferroni-Holm FDR correction. *** p < .01; - not reach significant level.

| **Components** | **Network** | **Standardized Beta** | **Significance** |
| --- | --- | --- | --- |
| PLS1+ | Visual | 0.136428184 | - |
| PLS1+ | Somatomotor | 0.213890298 | - |
| PLS1+ | Dorsal_attention | 0.268309554 | - |
| PLS1+ | Ventral_attention | -0.135594544 | - |
| PLS1+ | Limbic | 0.012724558 | - |
| PLS1+ | Frontal_parietal | -0.380804191 | *** |
| PLS1+ | Default_mode | -0.284015825 | - |
| PLS1- | Visual | -0.522499534 | *** |
| PLS1- | Somatomotor | -0.103394804 | - |
| PLS1- | Dorsal_attention | -0.206554892 | - |
| PLS1- | Ventral_attention | 0.341218657 | *** |
| PLS1- | Limbic | 0.111579815 | - |
| PLS1- | Frontal_parietal | 0.262378052 | - |
| PLS1- | Default_mode | 0.367425603 | *** |

**Table S19** Association of gene set in the PLS2 component for brain network properties in healthy brain. Statistical significance was set as p < .05 with Bonferroni-Holm FDR correction. *** p < .01; - not reach significant level.

10. **Decoding the brain cognitive ontology associations with PLS gene sets**

Yeo et al. (2015)^13^ have provided an ontology system to annotate brain cognitive functions into 12 components. We labeled the ontology for corresponding cognitive component by using most probability of corresponding task. More details can be found in the original paper. Ontology in the current study included as followed: Component 1, Vibrotactile Mon/Discrim; Component 2, Recitation/Repetition; Component 3, Pitch Mon/Discrim; Component 4, Visual Pursuit/Tracking; Component 5, Naming; Component 6, Saccades; Component 7, Micturition; Component 8, Flanker; Component 9, WCST; Component 10, Theory of Mind; Component 11, Face Mon/Discrim; Component 12, Reward Task. Full results have been tabulated into Table S20-21 for both PLS1 and PLS2 gene sets.

| **PLS** | **Components** | **Standardized Beta** | **Significance** |
| --- | --- | --- | --- |
| PLS1+ | Comp01 | -0.282429963 | - |
| PLS1+ | Comp02 | -0.062945239 | - |
| PLS1+ | Comp03 | 0.018547576 | - |
| PLS1+ | Comp04 | -0.339585193 | *** |
| PLS1+ | Comp05 | 0.023344023 | - |
| PLS1+ | Comp06 | -0.339267597 | *** |
| PLS1+ | Comp07 | 0.182144831 | - |
| PLS1+ | Comp08 | 0.234151567 | - |
| PLS1+ | Comp09 | -0.100035974 | - |
| PLS1+ | Comp10 | 0.011118617 | - |
| PLS1+ | Comp11 | 0.553758275 | *** |
| PLS1+ | Comp12 | 0.263268674 | - |
| PLS1- | Comp01 | 0.317497036 | *** |
| PLS1- | Comp02 | 0.133144417 | - |
| PLS1- | Comp03 | -0.029568398 | - |
| PLS1- | Comp04 | 0.409810001 | *** |
| PLS1- | Comp05 | 0.001799554 | - |
| PLS1- | Comp06 | 0.420528243 | *** |
| PLS1- | Comp07 | -0.183416792 | - |
| PLS1- | Comp08 | -0.271632972 | - |
| PLS1- | Comp09 | 0.1127442 | - |
| PLS1- | Comp10 | 0.02131915 | - |
| PLS1- | Comp11 | -0.58075032 | *** |
| PLS1- | Comp12 | -0.352797658 | *** |

**Table S20** Association of gene set in the PLS1 set for brain cognitive ontology (component). Statistical significance was set as p < .05 with Bonferroni-Holm FDR correction. *** p < .01; - not reach significant level.

| **PLS** | **Components** | **Standardized Beta** | **Significance** |
| --- | --- | --- | --- |
| PLS1+ | Comp01 | 0.176685225 | - |
| PLS1+ | Comp02 | 0.044590965 | - |
| PLS1+ | Comp03 | 0.017960419 | - |
| PLS1+ | Comp04 | 0.249992408 | - |
| PLS1+ | Comp05 | -0.058246861 | - |
| PLS1+ | Comp06 | 0.191637639 | - |
| PLS1+ | Comp07 | -0.050685074 | - |
| PLS1+ | Comp08 | -0.466846767 | *** |
| PLS1+ | Comp09 | -0.071769055 | - |
| PLS1+ | Comp10 | 0.075287099 | - |
| PLS1+ | Comp11 | -0.181662629 | - |
| PLS1+ | Comp12 | -0.578485118 | *** |
| PLS1- | Comp01 | -0.082911122 | - |
| PLS1- | Comp02 | -0.030039804 | - |
| PLS1- | Comp03 | -0.219761564 | - |
| PLS1- | Comp04 | -0.474572187 | *** |
| PLS1- | Comp05 | -0.067749174 | - |
| PLS1- | Comp06 | -0.115587084 | - |
| PLS1- | Comp07 | 0.214763497 | - |
| PLS1- | Comp08 | 0.597527658 | *** |
| PLS1- | Comp09 | 0.020308783 | - |
| PLS1- | Comp10 | -0.187314543 | - |
| PLS1- | Comp11 | 0.356684837 | *** |
| PLS1- | Comp12 | 0.605229286 | *** |

**Table S21** Association of gene set in the PLS2 set for brain cognitive ontology (component). Statistical significance was set as p < .05 with Bonferroni-Holm FDR correction. *** p < .01; - not reach significant level.

11. **Decoding the cortical metabolisms of gradient-derived PLS components for ADHD**

Vaishnavi and colleagues (2010) have revealed the regional aerobic glycolysis in the cortical areas and provided an atlas to quantify cortical metabolisms, including glycolytic index (GI), oxygen-glucose index (OGI), cerebral metabolic rate of oxygen/glucose (CMRO_2_/GMRGlu) and cerebral blood flow (CBF). Thus, to uncover whether these gene sets in PLS1 and PLS2 were associated with cortical metabolisms, the general linear models were built as well. Full results have been sorted into Table 22-23 as underneath.

| **PLS** | **Cortical metabolism** | **Standardized Beta** | **Significance** |
| --- | --- | --- | --- |
| PLS1+ | GI | -0.125159751 | - |
| PLS1+ | OGI | 0.111624035 | - |
| PLS1+ | CMRO2 | -0.636822586 | *** |
| PLS1+ | CMRGlu | -0.48557228 | *** |
| PLS1+ | CBF | -0.268486228 | - |
| PLS1- | GI | 0.05799824 | - |
| PLS1- | OGI | -0.037502097 | - |
| PLS1- | CMRO2 | 0.658603191 | *** |
| PLS1- | CMRGlu | 0.464496947 | *** |
| PLS1- | CBF | 0.243262562 | - |

**Table S22** Association of gene set in the PLS1 set for cortical metabolism. Statistical significance was set as p < .05 with Bonferroni-Holm FDR correction. *** p < .01; - not reach significant level.

| **PLS** | **Cortical metabolism** | **Standardized Beta** | **Significance** |
| --- | --- | --- | --- |
| PLS1+ | GI | -0.550774286 | *** |
| PLS1+ | OGI | 0.554244289 | *** |
| PLS1+ | CMRO2 | -0.192848032 | - |
| PLS1+ | CMRGlu | -0.422066928 | *** |
| PLS1+ | CBF | -0.441607207 | *** |
| PLS1- | GI | 0.504421764 | *** |
| PLS1- | OGI | -0.4986237 | *** |
| PLS1- | CMRO2 | -0.149947226 | - |
| PLS1- | CMRGlu | 0.167144476 | - |
| PLS1- | CBF | 0.321899581 | *** |

**Table S23** Association of gene set in the PLS2 set for cortical metabolism. Statistical significance was set as p < .05 with Bonferroni-Holm FDR correction. *** p < .01; - not reach significant level.

13. **Decoding cognitive terms of these PLS gene sets at NeuroSynth**

As mentioned above, the NeuroSynth platform provided an automatic meta-analytic tool to decode what cognitive terms are correlate with gene sets that we provided. In this vein, we decoded cognitive terms of gene set for PLS1+, PLS1-, PLS2+, and PLS2-, respectively. Full results have been provided into the Figure S10, with dark blue for significant association (p < .05, Bonferroni-Holm FDR corrected).

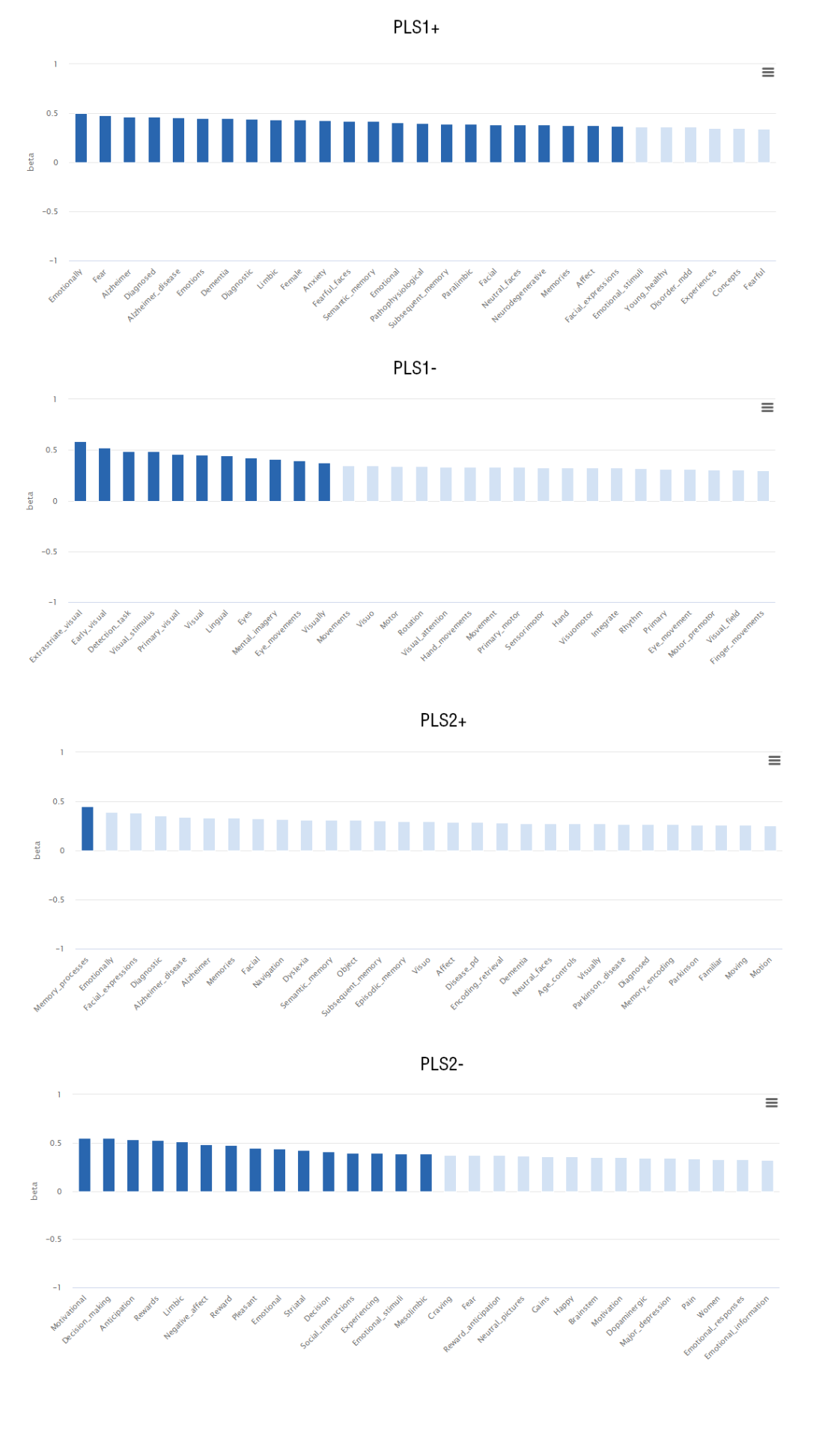

**Figure S10** Cognitive terms for PLS gene lists at NeuroSynth.

14. **Decoding neurological and psychiatric diseases at BrainMap**

Rather single-gene correlation, we decoded gene sets with PLS1 and PLS2 components at BrainMap to reveal the association between gene expression pattern and structural/functional abnormalities of 19 diseases in BrainMap dataset. BrainMap provided a outlet to do an online meta-analytic decoding in both VBM and functional MRI dataset. Full results have been documented into following Table S24-25.

| **Modality** | **Diseases** | **Standardized Beta** | **Significance** |
| --- | --- | --- | --- |
| VBM | ADHD | 0.070885793 | - |
| VBM | ALS | -0.14132548 | - |
| VBM | ASD | -0.096569226 | - |
| VBM | FTD | -0.336756821 | - |
| VBM | MCI | -0.180901465 | - |
| VBM | MS | 0.084484822 | * |
| VBM | OCD | -0.27570234 | - |
| VBM | PTSD | -0.026346629 | - |
| VBM | Alzheimers | -0.103448966 | - |
| VBM | Anxiety | -0.342734463 | - |
| VBM | Asperger | -0.060136288 | - |
| VBM | Bipolar | -0.27672006 | - |
| VBM | Dementia | -0.067555079 | - |
| VBM | Depression | -0.022635381 | - |
| VBM | Dyslexia | -0.057772047 | * |
| VBM | Huntington | 0.182756311 | * |
| VBM | Obesity | -0.063540086 | - |
| VBM | Parkinson | -0.080409798 | - |
| VBM | Psychosis | -0.118183804 | - |
| VBM | Schizophrenia | -0.343803194 | - |
| VBM | SementicDementia | -0.164201566 | - |
| VBM | Stroke | -0.072469242 | - |
| fMRI | ADHD | 0.070885793 | - |
| fMRI | ASD | 0.251908889 | *** |
| fMRI | MCI | -0.112453517 | - |
| fMRI | MDD | 0.141629602 | - |
| fMRI | OCD | -0.109707744 | - |
| fMRI | PTSD | -0.280033702 | - |
| fMRI | Alzheimers | 0.051524612 | - |
| fMRI | Anxiety | -0.276089282 | - |
| fMRI | Asperger | 0.317786145 | *** |
| fMRI | Bipolar | -0.168313731 | - |
| fMRI | Depression | -0.088994849 | - |
| fMRI | Dyslexia | 0.241381198 | *** |
| fMRI | Obesity | 0.075353004 | - |
| fMRI | Parkinson | 0.232744145 | *** |
| fMRI | Schizophrenia | 0.119380614 | - |
| fMRI | Stroke | 0.174196734 | *** |

**Table S24** Association of gene set in the PLS1 set for neurological and psychiatric diseases. Statistical significance was set as p < .05 with Bonferroni-Holm FDR correction. Each * represents that this p value reached statistical significance in one test. A total of five tests were used here: linear regression model test, null-spatial permutation, null-random-gene permutation, null-brain-gene permutation and null-coexpressed-gene permutation.

| **Modality** | **Diseases** | **Standardized Beta** | **Significance** |
| --- | --- | --- | --- |
| VBM | ADHD | 0.062714085 | - |
| VBM | ALS | 0.033036342 | - |
| VBM | ASD | 0.127415546 | - |
| VBM | FTD | 0.396852085 | * |
| VBM | MCI | 0.035839454 | - |
| VBM | MS | 0.139979391 | - |
| VBM | OCD | 0.443199756 | **** |
| VBM | PTSD | 0.096348586 | - |
| VBM | Alzheimers | 0.096705429 | - |
| VBM | Anxiety | 0.31331625 | *** |
| VBM | Asperger | 0.054393867 | - |
| VBM | Bipolar | 0.408324732 | **** |
| VBM | Dementia | 0.205470275 | - |
| VBM | Depression | 0.381611471 | - |
| VBM | Dyslexia | -0.230627972 | - |
| VBM | Huntington | 0.030660184 | - |
| VBM | Obesity | 0.046326205 | - |
| VBM | Parkinson | 0.028605402 | - |
| VBM | Psychosis | -0.176188777 | - |
| VBM | Schizophrenia | 0.396624279 | **** |
| VBM | SementicDementia | -0.028944837 | - |
| VBM | Stroke | 0.173876802 | ** |
| fMRI | ADHD | 0.388636486 | **** |
| fMRI | ASD | -0.083859782 | - |
| fMRI | MCI | 0.287881175 | *** |
| fMRI | MDD | 0.146257951 | - |
| fMRI | OCD | 0.311617325 | ** |
| fMRI | PTSD | 0.537533508 | **** |
| fMRI | Alzheimers | -0.088472866 | - |
| fMRI | Anxiety | 0.366418115 | **** |
| fMRI | Asperger | -0.153785865 | - |
| fMRI | Bipolar | 0.280670385 | *** |
| fMRI | Depression | 0.469255686 | **** |
| fMRI | Dyslexia | -0.039040309 | - |
| fMRI | Obesity | 0.151843697 | ** |
| fMRI | Parkinson | -0.110146446 | - |
| fMRI | Schizophrenia | 0.104395375 | - |
| fMRI | Stroke | 0.10386253 | - |

**Table S25** Association of gene set in the PLS2 set for neurological and psychiatric diseases. Statistical significance was set as p < .05 with Bonferroni-Holm FDR correction. Each * represents that this p value reached statistical significance in one test. A total of five tests were used here: linear regression model test, null-spatial permutation, null-random-gene permutation, null-brain-gene permutation and null-coexpressed-gene permutation.

15. **Enrichment analysis for PLS1 components**

To reveal the biological process and enrichment pathways, we used the Metascape tool by meta-analyzing PLS1 single-gene list for ADHD. For quality control, we limited the databsets into Gene Ontology (GO) and KGGE pathways only. All the annotations and datasets were updated recently (01-10-2022). All genes in the genome have been used as the enrichment background. P values for all the GO terms and pathways were corrected by Benjamini-Hochberg FDR method. Full results for enrichment analysis for PLS1 have been structured into Table S26-27 and Figure S11.

| **GO** | **Category** | **Description** | **Count** | **%** | **Log10(P)** | **Log10(q)** |
| --- | --- | --- | --- | --- | --- | --- |
| GO:0051046 | GO Biological Processes | regulation of secretion | 8 | 13.79 | -4.61 | -0.34 |
| GO:0050920 | GO Biological Processes | regulation of chemotaxis | 5 | 8.62 | -4.09 | -0.34 |
| GO:0099504 | GO Biological Processes | synaptic vesicle cycle | 4 | 6.9 | -4.04 | -0.34 |
| GO:0040013 | GO Biological Processes | negative regulation of locomotion | 6 | 10.34 | -3.82 | -0.31 |
| GO:0006939 | GO Biological Processes | smooth muscle contraction | 3 | 5.17 | -3.67 | -0.23 |
| GO:0003407 | GO Biological Processes | neural retina development | 3 | 5.17 | -3.4 | -0.05 |
| hsa05034 | KEGG Pathway | Alcoholism | 4 | 6.9 | -3.33 | -0.02 |
| GO:1903828 | GO Biological Processes | negative regulation of protein localization | 4 | 6.9 | -3.2 | -0.01 |
| R-HSA-418346 | Reactome Gene Sets | Platelet homeostasis | 3 | 5.17 | -3.18 | -0.01 |
| GO:0007188 | GO Biological Processes | adenylate cyclase-modulating G protein-coupled receptor signaling pathway | 4 | 6.9 | -2.98 | 0 |
| GO:0050727 | GO Biological Processes | regulation of inflammatory response | 5 | 8.62 | -2.92 | 0 |
| GO:1904375 | GO Biological Processes | regulation of protein localization to cell periphery | 3 | 5.17 | -2.74 | 0 |
| GO:0035725 | GO Biological Processes | sodium ion transmembrane transport | 3 | 5.17 | -2.68 | 0 |
| M5885 | Canonical Pathways | NABA MATRISOME ASSOCIATED | 6 | 10.34 | -2.5 | 0 |
| GO:0043086 | GO Biological Processes | negative regulation of catalytic activity | 6 | 10.34 | -2.38 | 0 |
| GO:0071248 | GO Biological Processes | cellular response to metal ion | 3 | 5.17 | -2.19 | 0 |
| GO:1901888 | GO Biological Processes | regulation of cell junction assembly | 3 | 5.17 | -2.15 | 0 |

**Table S26** Top 17 clusters with their representative enriched terms (one per cluster) for PLS1+ single-list component. "Count" is the number of genes in the user-provided lists with membership in the given ontology term. "%" is the percentage of all of the user-provided genes that are found in the given ontology term (only input genes with at least one ontology term annotation are included in the calculation). "Log10(P)" is the p-value in log base 10. "Log10(q)" is the multi-test adjusted p-value in log base 10.

| **GO** | **Category** | **Description** | **Count** | **%** | **Log10(P)** | **Log10(q)** |
| --- | --- | --- | --- | --- | --- | --- |
| hsa04020 | KEGG Pathway | Calcium signaling pathway | 8 | 10.53 | -6.77 | -2.42 |
| GO:0051384 | GO Biological Processes | response to glucocorticoid | 6 | 7.89 | -5.79 | -1.74 |
| R-HSA-9615017 | Reactome Gene Sets | FOXO-mediated transcription of oxidative stress, metabolic and neuronal genes | 3 | 3.95 | -4.27 | -0.77 |
| R-HSA-5683057 | Reactome Gene Sets | MAPK family signaling cascades | 6 | 7.89 | -3.76 | -0.49 |
| GO:0098655 | GO Biological Processes | cation transmembrane transport | 8 | 10.53 | -3.54 | -0.49 |
| GO:0035270 | GO Biological Processes | endocrine system development | 4 | 5.26 | -3.52 | -0.49 |
| GO:0010817 | GO Biological Processes | regulation of hormone levels | 7 | 9.21 | -3.48 | -0.49 |
| GO:0042886 | GO Biological Processes | amide transport | 4 | 5.26 | -3.36 | -0.44 |
| hsa05230 | KEGG Pathway | Central carbon metabolism in cancer | 3 | 3.95 | -3.13 | -0.35 |
| GO:0015918 | GO Biological Processes | sterol transport | 3 | 3.95 | -2.96 | -0.26 |
| GO:0043583 | GO Biological Processes | ear development | 4 | 5.26 | -2.64 | -0.07 |
| GO:0003015 | GO Biological Processes | heart process | 3 | 3.95 | -2.56 | -0.02 |
| hsa04919 | KEGG Pathway | Thyroid hormone signaling pathway | 3 | 3.95 | -2.45 | 0 |
| GO:0046661 | GO Biological Processes | male sex differentiation | 3 | 3.95 | -2.06 | 0 |

**Table S27** Top 17 clusters with their representative enriched terms (one per cluster) for PLS1- single-list component. "Count" is the number of genes in the user-provided lists with membership in the given ontology term. "%" is the percentage of all of the user-provided genes that are found in the given ontology term (only input genes with at least one ontology term annotation are included in the calculation). "Log10(P)" is the p-value in log base 10. "Log10(q)" is the multi-test adjusted p-value in log base 10.

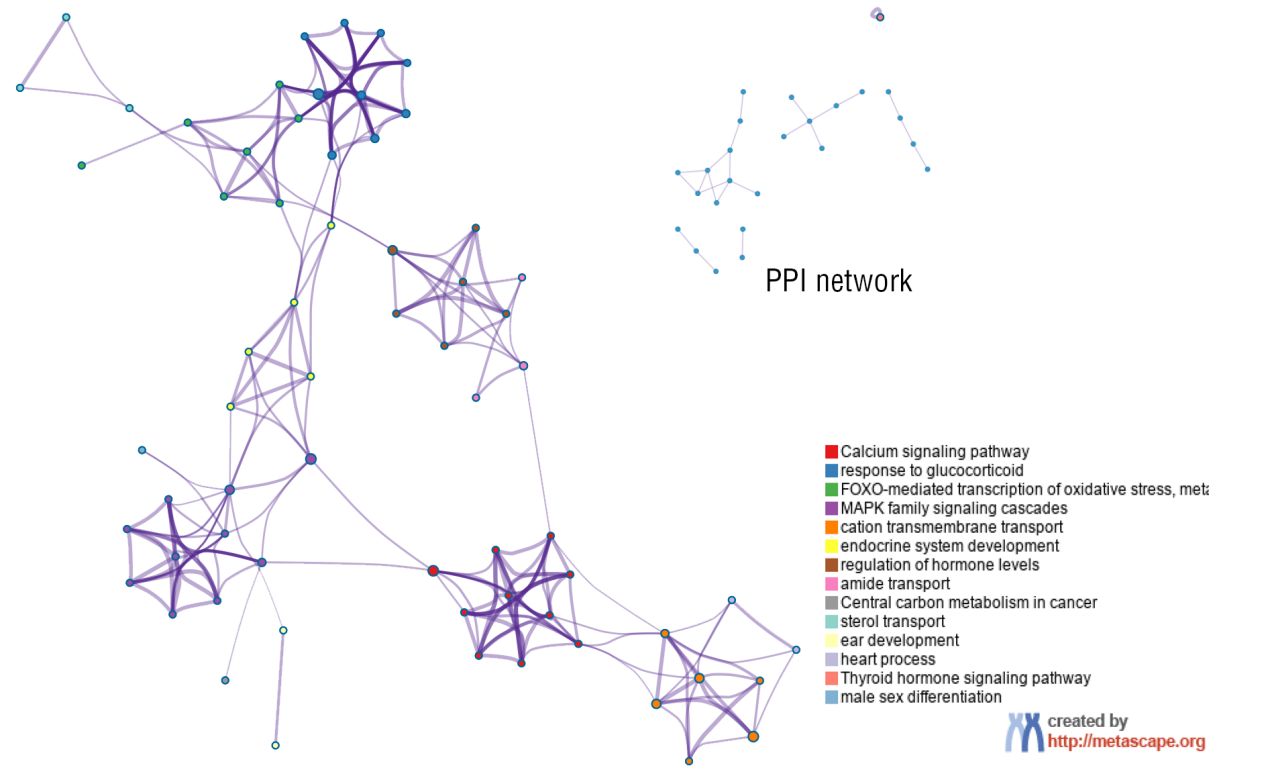

**Table S11** Network of enriched terms that colored by cluster ID. Nodes shared the same cluster ID are typically close to each other. Protein-protein interaction network has been provided as well. The network is visualized using Cytoscape.

16. **Muti-gene-list enrichment between GWAS and PLS-derived genes**

To determine whether the PLS-derived transcriptomic signatures shared with canonical polygenic risks of ADHD, we extracted risky genes from recent reviews with regard to the results of GWAS for ADHD. Further, we used multi-gene-list enrichment analysis to examine whether there were unique or shared biological process or pathways at Metascape as well. We found no shared genes between them, which indicated the PLS1- gene set that we revealed in the current study was unique for ADHD. Enrichment biological process and pathways have been reported in the Table S28 and Figure S12.

| **GO** | **Category** | **Description** | **Count** | **%** | **Log10(P)** | **Log10(q)** |
| --- | --- | --- | --- | --- | --- | --- |
| GO:0051588 | GO Biological Processes | regulation of neurotransmitter transport | 5 | 8.06 | -5.59 | -1.5 |
| GO:0051046 | GO Biological Processes | regulation of secretion | 9 | 14.52 | -5.31 | -1.5 |
| GO:0050920 | GO Biological Processes | regulation of chemotaxis | 6 | 9.68 | -5.1 | -1.45 |
| R-HSA-418346 | Reactome Gene Sets | Platelet homeostasis | 4 | 6.45 | -4.48 | -1.13 |
| GO:0051051 | GO Biological Processes | negative regulation of transport | 7 | 11.29 | -4.25 | -0.98 |
| GO:0006936 | GO Biological Processes | muscle contraction | 5 | 8.06 | -3.95 | -0.75 |
| GO:0040013 | GO Biological Processes | negative regulation of locomotion | 6 | 10.34 | -3.82 | -0.31 |
| GO:0035725 | GO Biological Processes | sodium ion transmembrane transport | 4 | 6.45 | -3.81 | -0.68 |
| GO:0043086 | GO Biological Processes | negative regulation of catalytic activity | 8 | 12.9 | -3.66 | -0.67 |
| GO:0003407 | GO Biological Processes | neural retina development | 3 | 5.17 | -3.4 | -0.05 |
| hsa05034 | KEGG Pathway | Alcoholism | 4 | 6.9 | -3.33 | -0.02 |
| GO:1904375 | GO Biological Processes | regulation of protein localization to cell periphery | 3 | 5.17 | -2.74 | 0 |
| GO:0042391 | GO Biological Processes | regulation of membrane potential | 5 | 8.06 | -2.71 | 0 |
| M5885 | Canonical Pathways | NABA MATRISOME ASSOCIATED | 6 | 10.34 | -2.5 | 0 |
| GO:1901888 | GO Biological Processes | regulation of cell junction assembly | 3 | 5.17 | -2.15 | 0 |

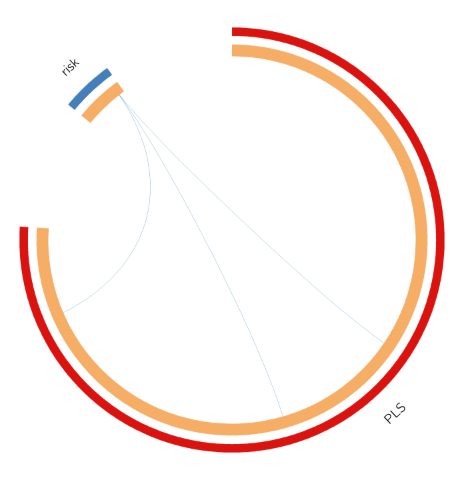
**Table S28** Top 15 clusters with their representative enriched terms (one per cluster) for multi-gene PLS1- component. "Count" is the number of genes in the user-provided lists with membership in the given ontology term. "%" is the percentage of all of the user-provided genes that are found in the given ontology term (only input genes with at least one ontology term annotation are included in the calculation). "Log10(P)" is the p-value in log base 10. "Log10(q)" is the multi-test adjusted p-value in log base 10.

**Figure S12** Overlap between gene lists. This plot including the shared term level, where blue curves link genes that belong to the same enriched ontology term. The inner circle represents gene lists, where hits are arranged along the arc. Genes that hit multiple lists are colored in dark orange, and genes unique to a list are shown in light orange. One gene that belong to same biological processes between them.

17. **Cell type-specific enrichment**

We have also analyzed PLS1+ gene-list enrichment by using Cell Type Signatures dataset^14^. All genes in the genome have been used as the enrichment background. Terms with a p-value < 0.01, a minimum count of 3, and an enrichment factor > 1.5 (the enrichment factor is the ratio between the observed counts and the counts expected by chance) are collected and grouped into clusters based on their membership similarities. We found 20 cell-type enrichment for PLS1+, such as HU FETAL RETINA RGC and MANNO MIDBRAIN NEUROTYPES HDA. Full list can be found in the Table S29.

| **GO** | **Description** | **Count** | **%** | **Log10(P)** | **Log10(q)** |
| --- | --- | --- | --- | --- | --- |
| [M39269](https://www.gsea-msigdb.org/gsea/msigdb/geneset_page.jsp?systematicName=M39269" \o "https://www.gsea-msigdb.org/gsea/msigdb/geneset_page.jsp?systematicName=M39269) | HU FETAL RETINA RGC | 11 | 19 | -9.1 | -4.6 |
| [M39068](https://www.gsea-msigdb.org/gsea/msigdb/geneset_page.jsp?systematicName=M39068" \o "https://www.gsea-msigdb.org/gsea/msigdb/geneset_page.jsp?systematicName=M39068) | MANNO MIDBRAIN NEUROTYPES HDA1 | 9 | 16 | -5.8 | -1.6 |
| [M39067](https://www.gsea-msigdb.org/gsea/msigdb/geneset_page.jsp?systematicName=M39067" \o "https://www.gsea-msigdb.org/gsea/msigdb/geneset_page.jsp?systematicName=M39067) | MANNO MIDBRAIN NEUROTYPES HDA | 8 | 14 | -5.3 | -1.3 |
| [M39069](https://www.gsea-msigdb.org/gsea/msigdb/geneset_page.jsp?systematicName=M39069" \o "https://www.gsea-msigdb.org/gsea/msigdb/geneset_page.jsp?systematicName=M39069) | MANNO MIDBRAIN NEUROTYPES HDA2 | 8 | 14 | -5.2 | -1.3 |
| [M39070](https://www.gsea-msigdb.org/gsea/msigdb/geneset_page.jsp?systematicName=M39070" \o "https://www.gsea-msigdb.org/gsea/msigdb/geneset_page.jsp?systematicName=M39070) | MANNO MIDBRAIN NEUROTYPES HNBGABA | 9 | 16 | -5.1 | -1.3 |
| [M39072](https://www.gsea-msigdb.org/gsea/msigdb/geneset_page.jsp?systematicName=M39072" \o "https://www.gsea-msigdb.org/gsea/msigdb/geneset_page.jsp?systematicName=M39072) | MANNO MIDBRAIN NEUROTYPES HSERT | 7 | 12 | -4.6 | -0.88 |
| [M41670](https://www.gsea-msigdb.org/gsea/msigdb/geneset_page.jsp?systematicName=M41670" \o "https://www.gsea-msigdb.org/gsea/msigdb/geneset_page.jsp?systematicName=M41670) | TRAVAGLINI LUNG LYMPHATIC CELL | 5 | 8.6 | -4.3 | -0.62 |
| [M39073](https://www.gsea-msigdb.org/gsea/msigdb/geneset_page.jsp?systematicName=M39073" \o "https://www.gsea-msigdb.org/gsea/msigdb/geneset_page.jsp?systematicName=M39073) | MANNO MIDBRAIN NEUROTYPES HOMTN | 6 | 10 | -4 | -0.42 |
| [M39168](https://www.gsea-msigdb.org/gsea/msigdb/geneset_page.jsp?systematicName=M39168" \o "https://www.gsea-msigdb.org/gsea/msigdb/geneset_page.jsp?systematicName=M39168) | MURARO PANCREAS ALPHA CELL | 7 | 12 | -4 | -0.42 |
| [M41691](https://www.gsea-msigdb.org/gsea/msigdb/geneset_page.jsp?systematicName=M41691" \o "https://www.gsea-msigdb.org/gsea/msigdb/geneset_page.jsp?systematicName=M41691) | TRAVAGLINI LUNG PLATELET MEGAKARYOCYTE CELL | 7 | 12 | -3.8 | -0.3 |
| [M39064](https://www.gsea-msigdb.org/gsea/msigdb/geneset_page.jsp?systematicName=M39064" \o "https://www.gsea-msigdb.org/gsea/msigdb/geneset_page.jsp?systematicName=M39064) | MANNO MIDBRAIN NEUROTYPES HNBML1 | 5 | 8.6 | -3.4 | 0 |
| [M39270](https://www.gsea-msigdb.org/gsea/msigdb/geneset_page.jsp?systematicName=M39270" \o "https://www.gsea-msigdb.org/gsea/msigdb/geneset_page.jsp?systematicName=M39270) | HU FETAL RETINA RPC | 3 | 5.2 | -2.9 | 0 |
| [M39290](https://www.gsea-msigdb.org/gsea/msigdb/geneset_page.jsp?systematicName=M39290" \o "https://www.gsea-msigdb.org/gsea/msigdb/geneset_page.jsp?systematicName=M39290) | DURANTE ADULT OLFACTORY NEUROEPITHELIUM OLFACTORY ENSHEATHING GLIA | 3 | 5.2 | -2.8 | 0 |
| [M39206](https://www.gsea-msigdb.org/gsea/msigdb/geneset_page.jsp?systematicName=M39206" \o "https://www.gsea-msigdb.org/gsea/msigdb/geneset_page.jsp?systematicName=M39206) | HAY BONE MARROW PLATELET | 4 | 6.9 | -2.7 | 0 |
| [M40223](https://www.gsea-msigdb.org/gsea/msigdb/geneset_page.jsp?systematicName=M40223" \o "https://www.gsea-msigdb.org/gsea/msigdb/geneset_page.jsp?systematicName=M40223) | DESCARTES FETAL KIDNEY METANEPHRIC CELLS | 3 | 5.2 | -2.5 | 0 |
| [M40214](https://www.gsea-msigdb.org/gsea/msigdb/geneset_page.jsp?systematicName=M40214" \o "https://www.gsea-msigdb.org/gsea/msigdb/geneset_page.jsp?systematicName=M40214) | DESCARTES FETAL INTESTINE ENS NEURONS | 3 | 5.2 | -2.4 | 0 |
| [M39055](https://www.gsea-msigdb.org/gsea/msigdb/geneset_page.jsp?systematicName=M39055" \o "https://www.gsea-msigdb.org/gsea/msigdb/geneset_page.jsp?systematicName=M39055) | MANNO MIDBRAIN NEUROTYPES HRGL2A | 5 | 8.6 | -2.2 | 0 |
| [M40019](https://www.gsea-msigdb.org/gsea/msigdb/geneset_page.jsp?systematicName=M40019" \o "https://www.gsea-msigdb.org/gsea/msigdb/geneset_page.jsp?systematicName=M40019) | BUSSLINGER GASTRIC X CELLS | 3 | 5.2 | -2.2 | 0 |
| [M39321](https://www.gsea-msigdb.org/gsea/msigdb/geneset_page.jsp?systematicName=M39321" \o "https://www.gsea-msigdb.org/gsea/msigdb/geneset_page.jsp?systematicName=M39321) | CUI DEVELOPING HEART VASCULAR ENDOTHELIAL CELL | 3 | 5.2 | -2.2 | 0 |
| [M39303](https://www.gsea-msigdb.org/gsea/msigdb/geneset_page.jsp?systematicName=M39303" \o "https://www.gsea-msigdb.org/gsea/msigdb/geneset_page.jsp?systematicName=M39303) | CUI DEVELOPING HEART C6 EPICARDIAL CELL | 3 | 5.2 | -2.2 | 0 |

**Table S29** Summary of enrichment analysis in Cell Type Signatures.

18. **Validation results**

To validate the reproducibility of key findings that the current study drawn, we included an independent homogenous sample (n = 22, ADHD/TD = 11/11) here. By replicating these analytic processes, we found the highly consistent pattern for explained ratio and cumulative explained variances between the original sample and validation sample (Figure S13).

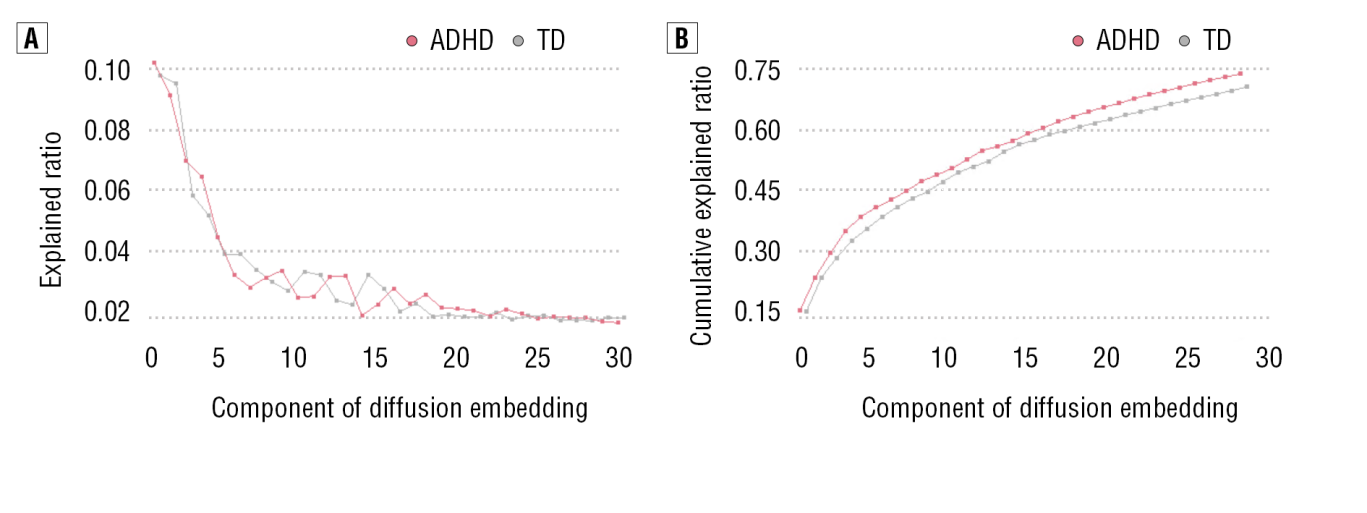
**Figure S13** Scree plot for component of diffusion embedding maps in the validation sample.

Further, we found the same results to differences of the explained ratio between ADHD and TD in validation, with no any significant differences for first-to-third gradients on explained ratio (Table S30).

| **Group** | **Mean** | **SD** | **P value** |
| --- | --- | --- | --- |
| Principal gradient | | | |
| ADHD | 9.95 | 4.32 | 0.773 ^corrected^ |
| TD | 9.97 | 4.60 |  |
| Second gradient | | | |
| ADHD | 9.21 | 4.60 | 0.845 ^corrected^ |
| TD | 8.82 | 6.99 |  |
| Third gradient | | | |
| ADHD | 5.29 | 6.63 | 0.627 ^corrected^ |
| TD | 6.50 | 6.99 |  |

**Table S30** Comparison between ADHD and TD for explained ratio in the first-to-third gradient.

With regard to other globally topological properties (i.e., gradient range and gradient variation), we fully replicated these findings that drawn from original sample by showing no significantly topological differences between ADHD and TD (Table S31-32).

| **Group** | **Mean** | **SD** | **P value** |
| --- | --- | --- | --- |
| Principal gradient | | | |
| ADHD | 0.12 | 0.02 | 0.177 ^corrected^ |
| TD | 0.14 | 0.03 |  |
| Second gradient | | | |
| ADHD | 0.11 | 0.03 | 0.644 ^corrected^ |
| TD | 0.12 | 0.04 |  |
| Third gradient | | | |
| ADHD | 0.09 | 0.01 | 0.312 ^corrected^ |
| TD | 0.10 | 0.04 |  |

**Table S31** Comparison between ADHD and TD for gradient range in the first-to-third gradient.

| **Group** | **Mean** | **SD** | **P value** |
| --- | --- | --- | --- |
| Principal gradient | | | |
| ADHD | 0.02 | 0.008 | 0.291 ^corrected^ |
| TD | 0.03 | 0.010 |  |
| Second gradient | | | |
| ADHD | 0.02 | 0.008 | 0.441 ^corrected^ |
| TD | 0.02 | 0.015 |  |
| Third gradient | | | |
| ADHD | 0.01 | 0.003 | 0.307 ^corrected^ |
| TD | 0.02 | 0.019 |  |

**Table S32** Comparison between ADHD and TD for gradient variant in the first-to-third gradient.

In addition to the globally topological properties, we still replicated the same analyses for system-based gradient perturbations of ADHD to TD. Results showed the moderately parallel system-based gradient patterns in validation sample, which indicated reproducibility in the current study. Full results have been documented in the Table S33-35.

| **System** | **Cohen d** | **t value** | **p value** | **q value ^FDR^** | **replication** |
| --- | --- | --- | --- | --- | --- |
| Visual | 0.27 | 14.49 | 0.000 | 0.000 | √ |
| Sensormotor | -0.01 | -1.04 | 0.293 | 0.470 | X |
| Dorsal attention | -0.40 | -18.05 | 0.000 | 0.000 | √ |
| Ventral attention | -0.04 | -1.86 | 0.030 | 0.061 | √ |
| Limbic | -0.07 | -8.95 | 0.003 | 0.042 | √ |
| Frontoparietal | -0.04 | -2.14 | 0.030 | 0.042 | X |
| Default mode | -0.40 | -26.45 | 0.000 | 0.000 | √ |
| Subcortical | -0.55 | -16.13 | 0.000 | 0.000 | √ |

**Table S33** Replication results for system-based principal gradient perturbations of ADHD in validation sample.

|  | VIS | SMN | DAN | VAN | LIM | FPN | DMN | SUB |
| --- | --- | --- | --- | --- | --- | --- | --- | --- |
| q (FDR) | 0.0000 | 0.0000 | 0.0198 | 0.0000 | 0.0000 | 0.0000 | 0.0000 | 0.0000 |
| Replication | √ | √ | √ | √ | √ | √ | √ | √ |

**Table S34** Replication results for system-based second gradient perturbations of ADHD in validation sample.

|  | VIS | SMN | DAN | VAN | LIM | FPN | DMN | SUB |
| --- | --- | --- | --- | --- | --- | --- | --- | --- |
| q (FDR) | 0.0033 | 0.9184 | 0.0000 | 0.0000 | 0.0001 | 0.6583 | 0.0199 | 0.0001 |
| Replication | √ | √ | √ | √ | √ | √ | √ | √ |

**Table S35** Replication results for system-based third gradient perturbations of ADHD in validation sample.

Further, we examined the map similarity between main analysis and replication analysis for vextex-based gradient. Results showed highly spatial correlation of them for mean map in principal gradient (r = .74, p < .01). We also checked these spatial similarities by comparing main findings to validation findings for ADHD and TD, respectively. We found moderate spatial correlation between them in the principal gradient (r = .67, p < .01 for HC; r = 0.73, p < .01 for ADHD). In this vein, these validations implied an acceptable reproducibility for our main findings.

To quantify reproducibility, the Jaccard index was used to estimate the comprehensive similarity between these samples pertaining to all gradient-related properties (e.g., global properties, system-based properties and vextex-based statistics). The Jaccard coefficient was mathematically described as ratio of intersection of A and B “areas” to union “areas”:

$$Ja (A, B)=\frac{\left| A\cap B \right|}{\left| A\cup B \right|}$$

where A represented binary results of main dataset and B meant the replicated results in validation dataset by referring A. Results showed high results similarity that was quantified by Jaccard coefficient (Ja = 0.785, 95% CI: 0.75 - 0.96), which indicated a favorable reproducibility for the current study.
